# A new gene family diagnostic for intracellular biomineralization of amorphous Ca-carbonates by cyanobacteria

**DOI:** 10.1101/2021.11.01.465933

**Authors:** Benzerara Karim, Duprat Elodie, Bitard-Feildel Tristan, Caumes Géraldine, Cassier-Chauvat Corinne, Chauvat Franck, Dezi Manuela, Diop Seydina Issa, Gaschignard Geoffroy, Görgen Sigrid, Gugger Muriel, López-García Purificación, Skouri-Panet Fériel, Moreira David, Callebaut Isabelle

## Abstract

Cyanobacteria have massively contributed to carbonate deposit formation over the geological history. They are traditionally thought to biomineralize CaCO_3_ extracellularly as an indirect byproduct of photosynthesis. However, the recent discovery of freshwater cyanobacteria forming intracellular amorphous calcium carbonates (iACC) challenges this view. Despite the geochemical interest of such a biomineralization process, its molecular mechanisms and evolutionary history remain elusive. Here, using comparative genomics, we identify a new gene (*ccyA*) and protein (calcyanin) family specifically associated with cyanobacterial iACC biomineralization. Calcyanin is composed of a conserved C-terminal domain, which likely adopts an original fold, and a variable N-terminal domain whose structure allows differentiating 4 major types among the 35 known calcyanin homologues. Calcyanin lacks detectable full-length homologs with known function. Yet, genetic and comparative genomic analyses suggest a possible involvement in Ca homeostasis, making this gene family a particularly interesting target for future functional studies. Whatever its function, this new gene family appears as a gene diagnostic of intracellular calcification in cyanobacteria. By searching for *ccyA* in publicly available genomes, we identified 13 additional cyanobacterial strains forming iACC. This significantly extends our knowledge about the phylogenetic and environmental distribution of cyanobacterial iACC biomineralization, especially with the detection of multicellular genera as well as a marine species. Phylogenetic analyses indicate that iACC biomineralization is ancient, with independent losses in various lineages and some HGT cases that resulted in the broad but patchy distribution of calcyanin across modern cyanobacteria. Overall, iACC biomineralization emerges as a new case of genetically controlled biomineralization in bacteria.

**Significance statement:** Few freshwater species of Cyanobacteria have been known to mineralize amorphous CaCO3 (ACC) intracellularly. Despite the geochemical interest of this biomineralization, its evolutionary history and molecular mechanism remain poorly known. Here, we report the discovery of a new gene family that has no homolog with known function, which proves to be a good diagnostic marker of this process. It allowed to find cyanobacteria in several phyla and environments such as seawater, where ACC biomineralization had not been reported before. Moreover, this gene is ancient and was independently lost in various lineages with some later horizontal transfers, resulting in a broad and patchy phylogenetic distribution in modern cyanobacteria.

## INTRODUCTION

The formation of mineral phases by living organisms is widespread in both eukaryotes and prokaryotes (Weiner and Dove 2003). While many cases of biomineralization in eukaryotes involve specific genes (Marron et al. 2016; Wang et al. 2021; Yarra et al. 2021), there is presently only one documented case of genetically controlled biomineralization in bacteria: the intracellular magnetite formation by magnetotactic bacteria (Lefevre and Bazylinski 2013). A genetic control of iACC biomineralization by several species of cyanobacteria has been recently hypothesized (Benzerara et al. 2014), but not yet proven. Interestingly, the involvement of ACC has been widely documented and studied in the formation of eukaryotic skeletons (Blue et al. 2017). By contrast and although a growing number of bacterial occurrences are described (Monteil et al. 2020), the determinants of ACC formation in prokaryotes remain poorly understood.

These iACC-biomineralizing cyanobacteria are geographically widespread in freshwater, hotspring or karstic terrestrial systems (Ragon et al. 2014) and sometimes locally abundant (Bradley et al. 2017). They received particular attention since they challenge the usual paradigm that cyanobacteria biomineralize CaCO_3_ extracellularly as an indirect byproduct of photosynthesis only (Altermann et al. 2006). Moreover, the geological history of iACC biomineralization remains mysterious since the fossilization potential of these bacteria appears uncertain (Couradeau et al. 2012; Riding 2012). They can form iACC even under thermodynamically unfavorable conditions, indicating that they consume energy to perform this process, possibly in relation with active sequestration of alkaline earth elements (AEEs) (Cam et al. 2018). An envelope of undetermined composition, either a lipid monolayer and/or proteins, surrounds the iACC granules (Blondeau, Sachse, et al. 2018) and may be instrumental for the achievement of local Ca concentrations that are high enough for the formation of iACC. Furthermore, some iACC-forming species require higher Ca amounts for optimal growth than iACC-lacking ones, indicating that they possess an unusual Ca homeostasis (De Wever et al. 2019). Interestingly, by forming iACC granules, these cyanobacteria accumulate very high Ca amounts, as well as other AEEs such as strontium (Sr) and barium (Ba) (Cam et al. 2016; Blondeau, Benzerara, et al. 2018) and may impact the geochemical cycles of these trace elements (Blondeau, Benzerara, et al. 2018). Indeed, by normalizing the uptake to their cell mass, they are among the highest Sr and Ba-scavenging organisms known (Cam et al. 2016). They can also very efficiently sequester radioisotopes such as ^90^Sr or radium (Ra) isotopes, a capability that may be used for bioremediation (Cam et al. 2016; Blondeau, Benzerara, et al. 2018; Mehta et al. 2019).

All the members of some clades of cyanobacteria, such as the one including *Thermosynechococcus elongatus* BP-1, share this capability to form iACC, suggesting the genetic heritability of this trait in this specific group (Benzerara et al. 2014). Yet, despite the geochemical relevance of this process, the genetic control of iACC formation has not been identified. Moreover, whether the presently known iACC-forming cyanobacteria share ancestral genetic traits related to this biomineralization process or they convergently developed this capability to form iACC during cyanobacterial evolution remains unknown. In the absence of a fossil record, investigating the genetic basis of this biomineralization process appears as the only way to track its geological history.

## Results and Discussion

### Detection of a gene family diagnostic of iACC biomineralization

We applied comparative genomics to search for putative genes exclusively shared by iACC-forming cyanobacteria, and therefore absent in non-iACC forming species. We analyzed the genomes of 56 cyanobacterial strains (supplementary table 1), in which the presence or absence of iACC was previously determined by electron microscopy (EM) (Benzerara et al. 2014). Fifty strains were lacking iACC and 6 were shown to form iACC: *Synechococcus* sp. PCC 6312, *Synechococcus calcipolaris* PCC 11701, *Thermosynechococcus elongatus* BP-1, *Cyanothece* sp. PCC 7425, *Chroococcidiopsis thermalis* PCC 7203, and *Gloeomargarita lithophora* D10. Among the 523 680 translated coding sequences (CDSs) retrieved from the 56 genomes, only one group of orthologous sequences was shared by all 6 iACC-forming strains and absent in all 50 iACC-lacking strains. The corresponding gene and its predicted protein product were named *ccyA* and calcyanin, respectively. Conversely, we found no group of orthologous sequences shared by all 50 iACC-lacking strains and absent in all 6 iACC-forming strains. No functional annotation of calcyanin could be achieved using profiles of known protein domain families. We first investigated the architecture of calcyanin by Hydrophobic Cluster Analysis (HCA) (Bitard-Feildel et al. 2018). Calcyanin was composed of two domains (fig. 1). While the C-terminal domain was highly conserved in the six different calcyanin sequences, the N-terminal domain appeared to be conserved in five sequences only, and exhibited significant differences in *G. lithophora*. Therefore, we used the conserved C-terminal domain to search for additional homologs in a comprehensive set of 594 cyanobacterial genomes, available in databases. We found additional *ccyA* homologs in 27 strains (supplementary table 2; supplementary fig. 1). Among them, we inspected 17 strains available to us, by EM coupled with energy dispersive x-ray spectrometry (EDXS), which allowed chemical mapping at the submicrometer-scale. We detected iACC in 13 of them (fig. 2), thereby increasing the number of known iACC-forming cyanobacterial species from 6 to 19. Moreover, we confirmed the presence of *ccyA* in the two recently sequenced genomes of *Synechococcus* sp. PCC 6716 and PCC 6717 that were previously shown to form iACC (Benzerara et al. 2014).

**Figure 1:**
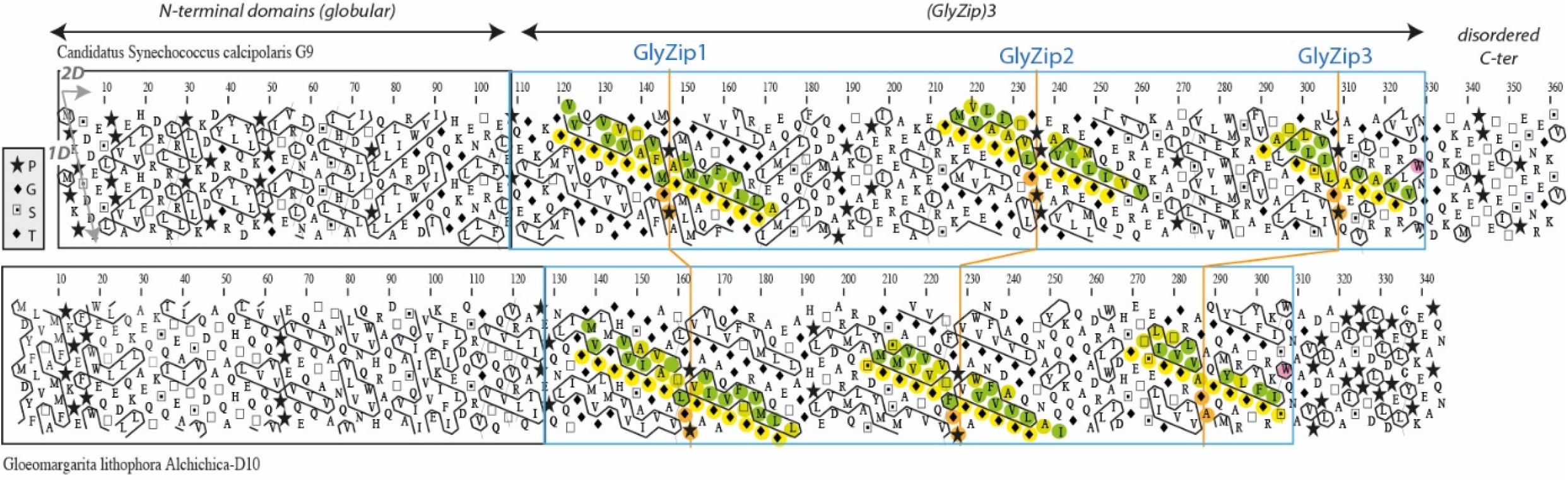
Domain architecture of calcyanins, as viewed by Hydrophobic Cluster Analysis. HCA plots of the calcyanin sequences of *Synechococcus calcipolaris* PCC 11701 and *Gloeomargarita lithophora* D10. The protein sequences are displayed on a duplicated alpha-helical net, on which the strong hydrophobic amino acids (V I L F M Y W) are contoured. The latter form clusters, which mainly correspond to the internal faces of regular secondary structures. The way to read the sequence (1D) and secondary structures (2D) are shown with arrows, whereas special symbols are described in the inset. After analysis of the two calcyanin sequences, two different domains can be highlighted, based on the identification of a common C-terminal domain composed of three long repeats (called GlyZips) of a periodic pattern, including glycine (or small amino acid - yellow) and hydrophobic amino acids (green) every four residues. These patterns give rise to long, horizontal hydrophobic clusters, which do not correspond to any known 3D structures and are likely to adopt an original fold, built around this repeated structure. This C-terminal domain is clearly distinct from the N-terminal domains, possessing smaller hydrophobic clusters usually encountered in current globular domains. No obvious sequence similarity could however be found between the *Gloeomargarita lithophora* calcyanin and the other group of sequences.

**Figure 2.**
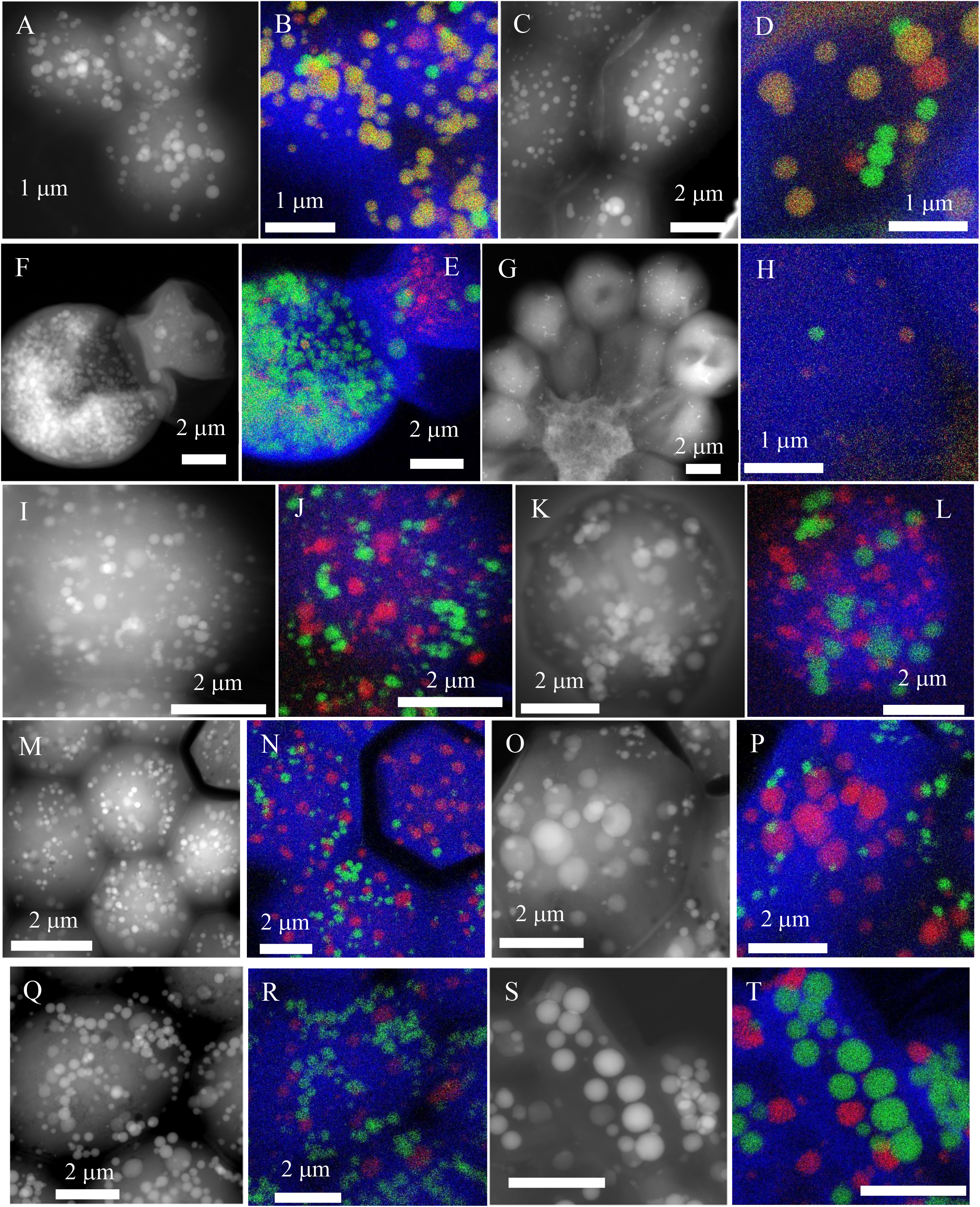
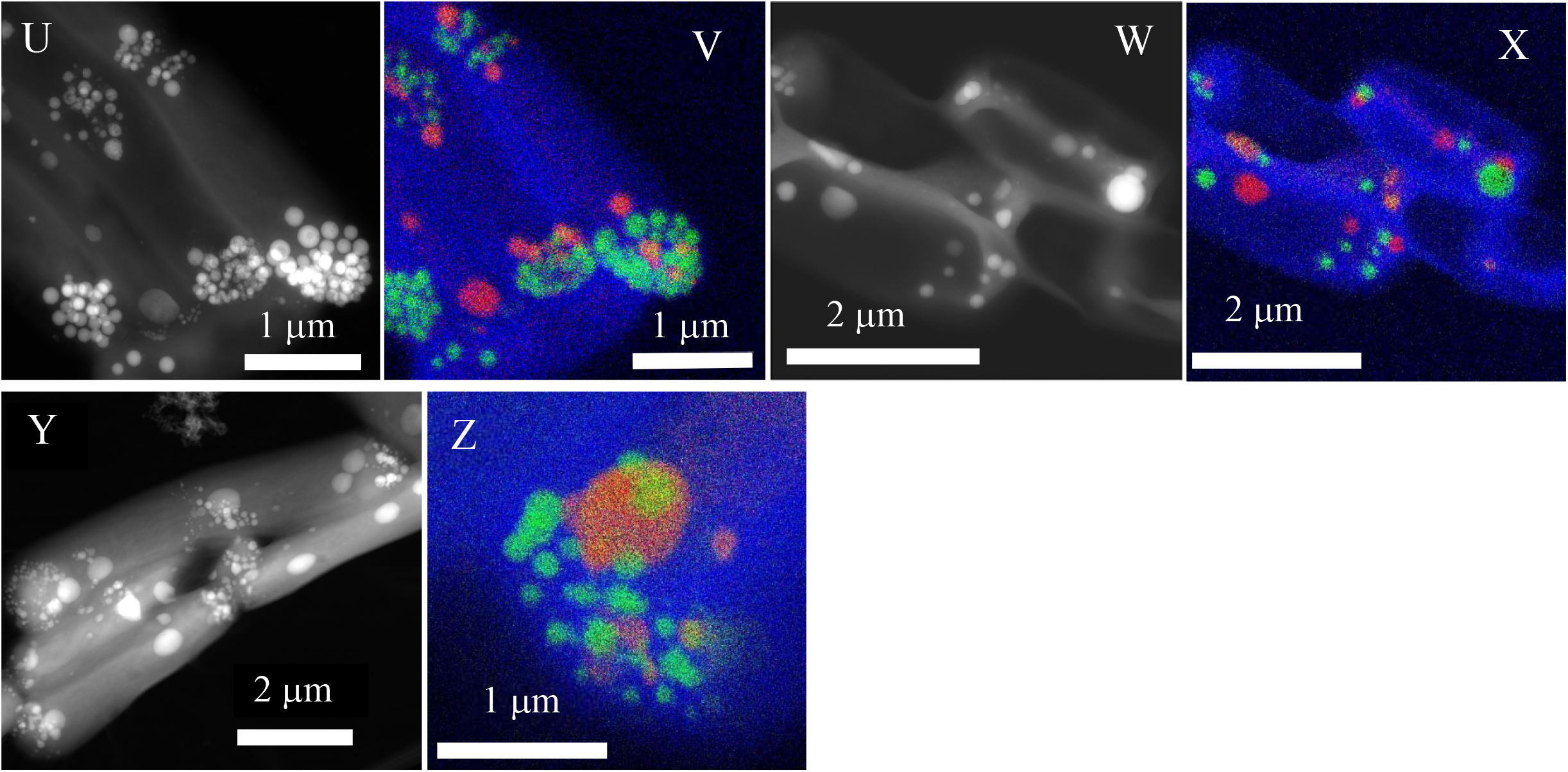
Electron microscopy detection of iACC in 13 calcyanin-bearing cyanobacterial strains not previously known to biomineralize carbonates. STEM-HAADF images of the 13 newly-identified iACC-forming strains and overlays of C (blue), Ca (green) and P (red) chemical maps as obtained by energy dispersive x-ray spectroscopy. A and B: *Chlorogloeopsis fritschii* PCC 9212; C and D: *Fischerella muscicola* PCC 7414; E and F: *Fischerella* sp. NIES-4106; G and H: *Microcystis aeruginosa* PCC 7806; I and J: *M. aeruginosa* PCC 7941; K and L: *M. aeruginosa* PCC 9443; M and N: *M. aeruginosa* PCC 9806; O and P: *M. aeruginosa* PCC 9807; Q and R: *M. aeruginosa* PCC 9808; S and T: *Neosynechococcus sphagnicola* sy1; U and V: *Synechococcus lividus* PCC 6715; W and X: *Synechococcus* sp. RS9917; Y and Z: *Thermosynechococcus* sp. NK55.

In some strains (e.g., *Fischerella* sp. NIES-4106, *Neosynechococcus sphagnicola* sy1), most of the cells exhibited abundant iACC granules. By contrast, only few cells contained few iACC granules for strains such as *Microcystis aeruginosa* PCC 7806. In other strains (e.g., *Chlorogloeopsis fritschii* PCC 9212), the cells contained few iACC granules and many Ca-rich polyphosphate inclusions that could be confused with iACC by EM alone (fig. 2). The four strains possessing *ccyA* but lacking iACC (*C. fritschii* PCC 6912; *Fischerella* sp. NIES-3754; *M. aeruginosa* PCC 9432 and PCC 9717; fig. 3) were phylogenetically very close to iACC-forming relatives. For example, *C. fritschii* PCC 9212 (iACC-forming) and PCC 6912 (no observed iACC) had only few differences in their gene repertoires (supplementary fig. 2; supplementary table 3) and the nucleotide sequences of the genomic regions containing *ccyA* were strictly identical in both strains. However, 57 genes of *C. fritschii* PCC 9212 had no homolog in *C. fritschii* PCC 6912. Their COG functional categories mostly corresponded to unknown functions (46 without COG hit, 2 genes with COG category X indicating an unknown function) or inorganic ion transport (4 genes, COG category P; supplementary table 3). Moreover, although we did not observe iACC in *C. fritschii* PCC 6912 and *Fischerella* sp. NIES-3754 cells, they both showed Ca- and P-rich inclusions morphologically similar to iACC (fig. 3). This may reflect the lack of expression of *ccyA* in these two strains under our culture conditions or the absence of other genes necessary for iACC biomineralization. For the two Microcystis strains possessing ccyA but lacking iACC, the extensive methylation observed in *Microcystis* genomes (Zhao et al. 2018) might add a potential epigenetic control on gene expression and genome rearrangements, also abundant, might disrupt gene dosage effects and/or impede gene co-expression (Frangeul et al. 2008; Humbert et al. 2013). At any rate, the search for *ccyA* in available cyanobacterial genome sequences allowed the detection of 13 additional iACC-forming strains among the 17 strains whose genomes contained *ccyA*, largely extending and optimizing the initial detection of 8 iACC-forming strains among 58 randomly selected, phylogenetically diverse cyanobacteria (Benzerara et al. 2014). Therefore, the search for *ccyA* occurrence significantly increased the probability of success in finding iACC-forming strains (Binomial exact test, p=9.0e-09), indicating that *ccyA* can be used as diagnostic marker of intracellular biomineralization.

**Figure 3:**
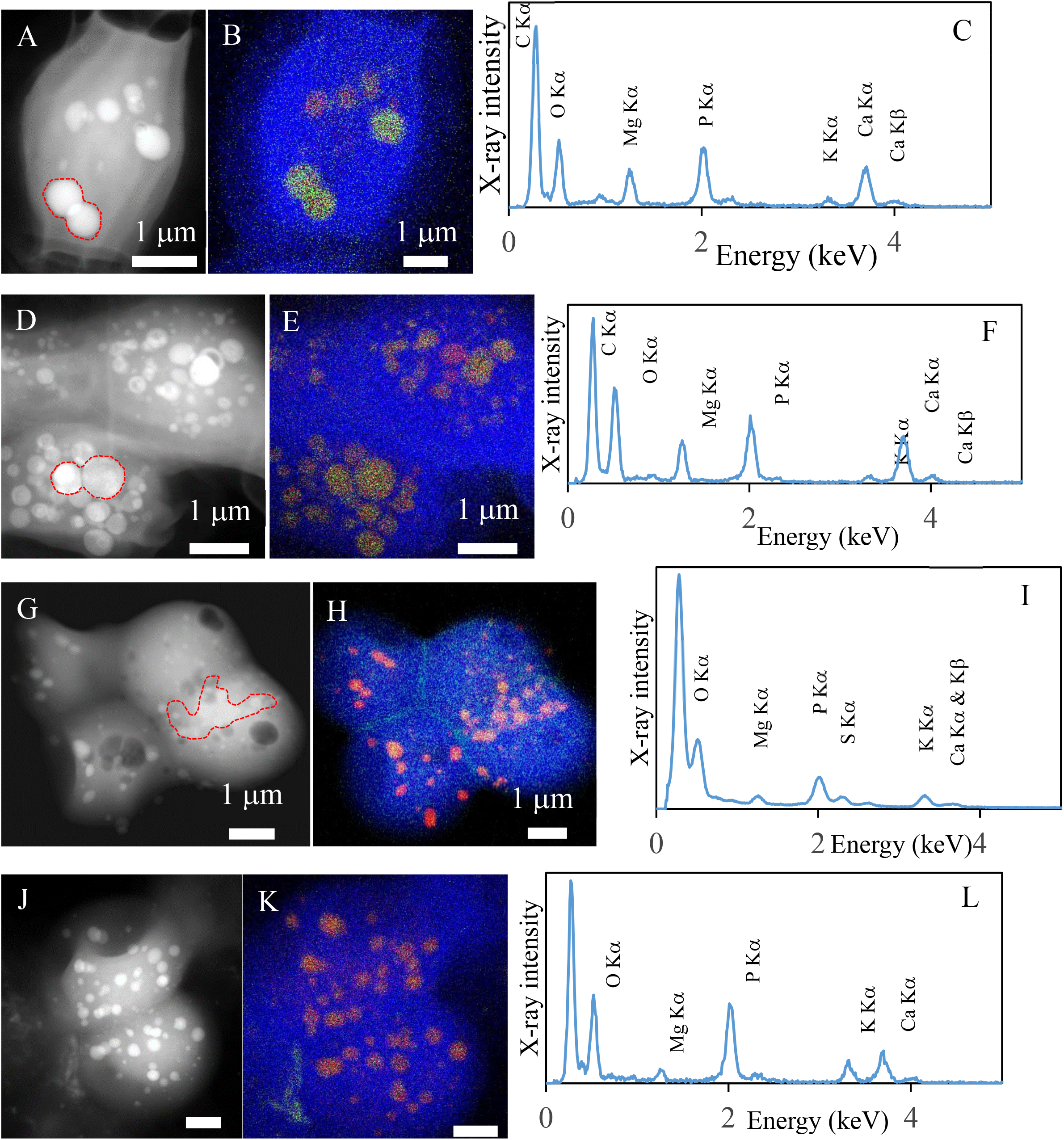
TEM analyses of ccyA-harbouring strains not forming iACC. Four strains harboring ccyA in their genomes but not showing iACC and overlays of C, Ca and P EDXS maps. A) STEM-HAADF image, B) overlays of C (blue), Ca (green) and P (red) EDXS maps and C) EDXS spectrum of *Fischerella* sp. NIES-3754. EDXS spectrum is extracted from the area indcated in A) by a dashed line; D, E and F: *Chlorogloeopsis fritschii* PCC 6912. G, H and I: *Microcystis aeruginosa* PCC 9432; J, K and L: *M. aeruginosa* PCC 9717.

Thanks to this approach, we detected both *ccyA* and iACC in several cyanobacterial genera where iACC biomineralization had not been observed before, thereby expanding considerably the phylogenetic diversity of iACC-forming cyanobacteria (fig. 4a). While iACC biomineralization had so far been reported in unicellular cyanobacteria only (Benzerara et al. 2014), we presently report it in multicellular genera belonging to the most complex morphotypes of the cyanobacterial phylum with cellular differentiation and ramifications (*Chlorogloeopsis, Fischerella* and *Mastigocladus*). Moreover, we also discovered iACC in *Microcystis aeruginosa*, one of the most common, worldwide-distributed bloom-forming cyanobacteria (Humbert et al. 2013). *Microcystis* shows a life cycle with a benthic phase in winter and a planktonic phase in warmer seasons when cells produce gas vesicles to float in the water column (Reynolds and Rogers 1976; Latour et al. 2007). Considering the high density of ACC relative to cells, a controlled production of dense iACC granules might favor a shift to benthic life. Interestingly, *ccyA* is present in the genome of some *M. aeruginosa* strains but absent from others. This finding may be consistent with the high genome plasticity detected in this species, reflecting frequent horizontal gene transfers (Frangeul et al. 2008; Humbert et al. 2013).

**Figure 4:**
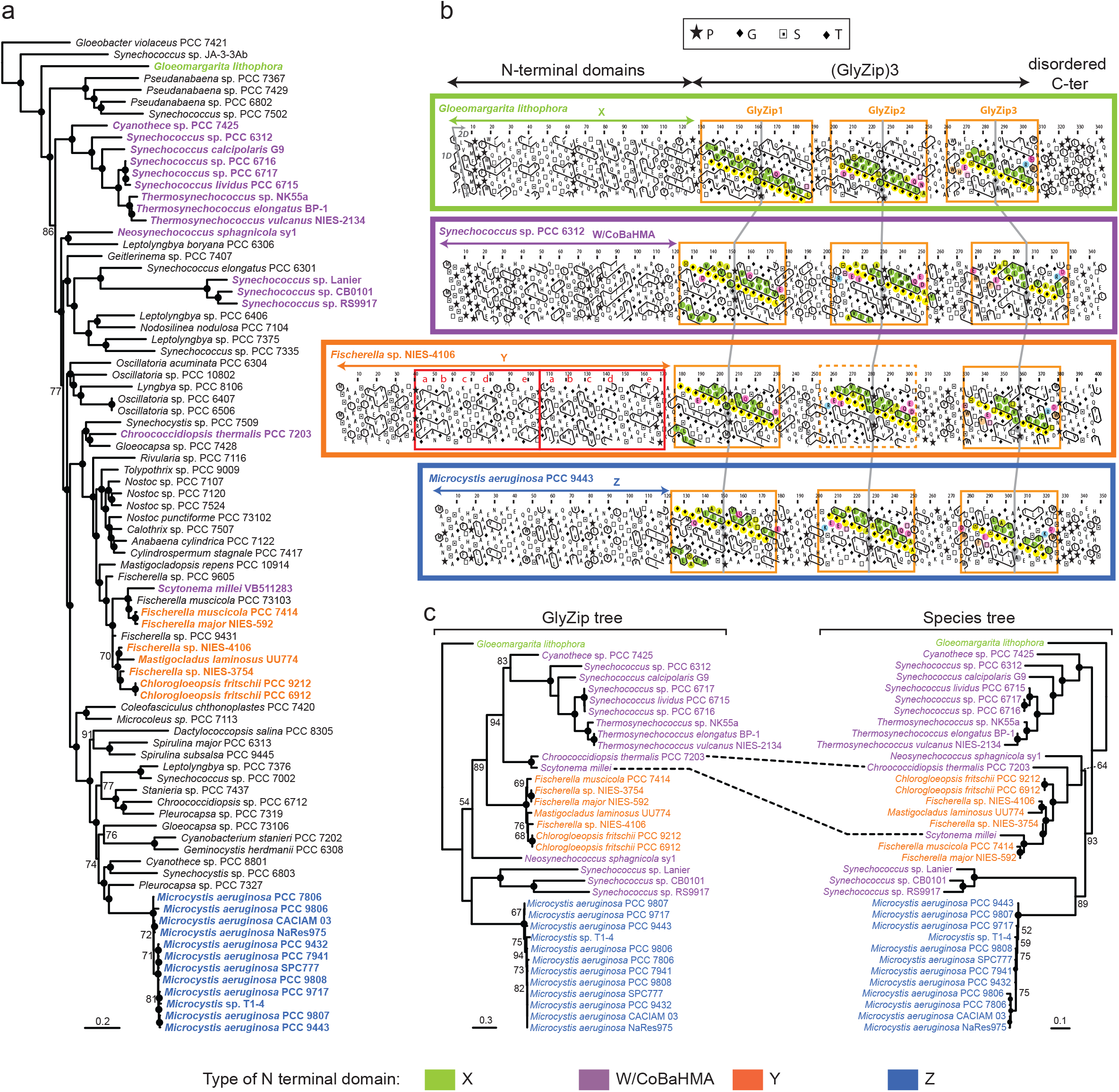
Phylogenetic analysis and domain architecture of the calcyanin protein family. **a**, Maximum likelihood phylogenetic tree of Cyanobacteria based on 58 conserved proteins; the strains containing the ccyA gene are highlighted in bold and color. **b**, HCA plots of representative calcyanin sequences. The protein sequences are displayed on a duplicated alpha-helical net, on which the strongly hydrophobic amino acids (V I L F M Y W) are contoured. The latter form clusters, which mainly correspond to the internal faces of regular secondary structures. The way to read the sequence is shown with an arrow, whereas special symbols are described in the inset. The positions of the domains are indicated, with the duplicated sub-module composing domain Y. The periodic pattern, made of glycine (or small amino acids – yellow) and hydrophobic amino acids (green) are highlighted for each GlyZip, with conserved signatures specific of each GlyZip shown with other colors. GlyZip2, which is present in only one species in the Y family, is indicated with dotted box. **c**, Maximum likelihood phylogenetic tree of the GlyZip domain of calcyanin (left) compared with the species tree based on 58 conserved proteins (right); the dashed lines indicate a probable case of recent horizontal ccyA gene transfer. Numbers on branches indicate bootstrap support (BS, only values >50% are shown), BS of 100% are indicated by black circles. The species names and HCA profiles are color-coded according to the type of N-terminal domain of calcyanins (the code is shown at the bottom of the figure).

The *ccyA* gene and iACC biomineralization were also found in four *Synechococcus* sp. strains previously not known to produce iACC (*Neosynechococcus sphagnicola* sy1, *Synechococcus* sp. RS9917, *Synechococcus lividus* PCC 6715, *Thermosynechococcus* sp. NK55a). *Synechococcus* is a polyphyletic genus, grouping strains isolated from significantly different environments (Komarek et al. 2020). We previously reported thermophilic and mesophilic freshwater iACC-biomineralizing *Synechococcus* representatives (Benzerara et al. 2014). Here, we significantly expanded this environmental distribution especially with the inclusion of the first marine (*Synechococcus* sp. RS9917) iACC-forming strain.

### Sequence-based analysis of the calcyanin structure

With the exception of the *Thermosynechococcus* sp. NK55a calcyanin, fused with a polypeptide containing a PIN-TRAM domain, the other 34 calcyanin family homologs contained 264 to 375 amino acids (average 336±25; supplementary table 2). All showed the already mentioned 2-domain modularity: a variable N-terminal domain and a conserved C-terminal domain.

The N-terminal domain was composed of hydrophobic clusters with lengths and shapes typical of regular secondary structures found in globular domains (Lamiable et al. 2019). According to their N-terminal domain, we classified the 35 calcyanin homologs into four groups: W, X, Y, and Z (fig. 4b). The X- and Z-type N-terminal domains were distinct from known protein domains. The Y-type N-terminal domain consisted of a duplicated small domain, with no obvious similarity with any known domain (supplementary fig. 3). By contrast, significant sequence similarities were detected between the W-type N-terminal domain and three known domain families (fig. 5): 1) YAM domains, found in the cytosolic C-terminus of *Escherichia coli* Major Facilitator Superfamily transporter YajR (Jiang et al. 2013; Jiang et al. 2014); 2) Heavy-Metal Associated (HMA) domains (also called Metal Binding Domains) present in various proteins (e.g., P-type ATPases and metallochaperones), generally involved in metal transport and detoxification pathways (Bull and Cox 1994); and 3) integrated HMA (iHMA) domains detected in plant immune receptors, where they are involved in fungal effector recognition (De la Concepcion et al. 2018). Similarly to these three domains, the W-type N-terminal domain showed a repeated β – α – β motif corresponding to a ferredoxin-like fold, characteristic of the HMA superfamily (fig. 5). However, while most HMA domains possess two conserved cysteine residues directly involved in binding heavy metals, YAM, iHMA and W-type calcyanin N-terminal domains do not conserve these amino acids (fig. 5). Moreover, the W-type domain showed a specific signature consisting of several basic amino acids distributed in strands β1 and β2 and a histidine located upstream of strand β1, in a region appearing as a calcyanin-specific extension of the HMA core (strand β0 in fig. 5). Therefore, we named this novel domain family CoBaHMA, after *domain with <u>Co</u>nserved <u>Ba</u>sic residues in the <u>HMA</u> superfamily*. Although the calcyanin sequence of *Synechococcus* sp. Lanier also contained the specific signature of W-type domains with several basic amino acids, it clearly differed from the rest of the W-type N-terminal domains (fig. 5A), suggesting that calcyanin has deeply diverged in this species.

**Figure 5:**
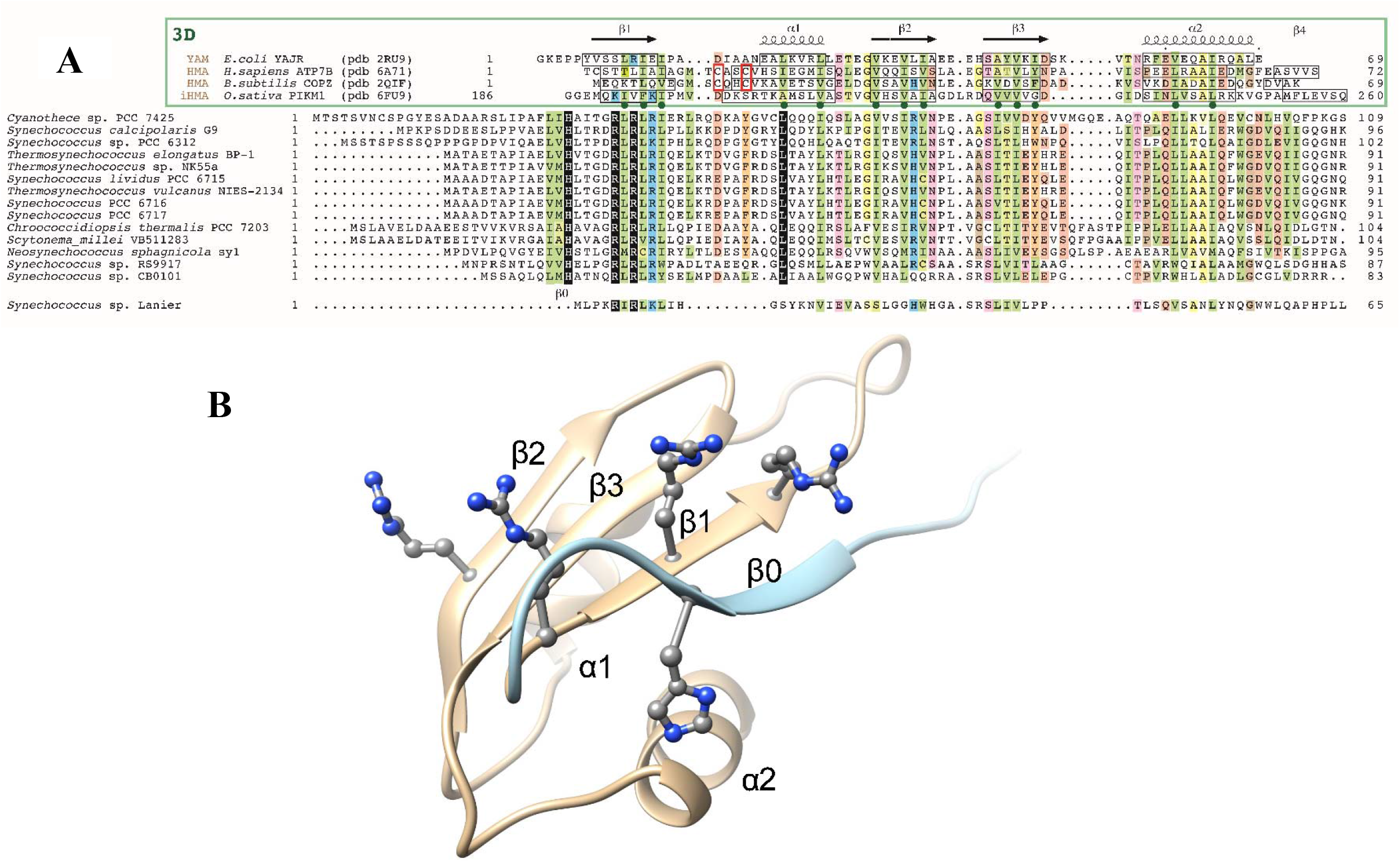
The CoBaHMA domain. **A**. Multiple sequence alignment of calcyanins and members of the HMA superfamily, with known 3D structures. Identical amino acids are shown white on a black background, similarities are colored according to amino acid properties: (i) green: hydrophobic (dark for strong hydrophobic amino acids (V, I, L, F, M, Y, W), light for other amino acids which can also occupy such positions (A, T, S, C)), (ii) yellow: small amino acids (G, A),(iii) orange: aromatic amino acids (W, F, Y), (iv): blue: basic amino acids (K,R,H), (v) pink: polar, non-basic amino acids (D, E, Q, N, S, T), (vii) brown: loop-forming amino acids (P, G, D, N, S), with positions mainly occupied by P highlighted darker. Sequences of proteins of the HMA superfamily, whose 3D structures are known and with which the CoBaHMA sequences can be aligned, are shown on top. PDB identifiers are provided. Observed 2D structures are boxed. The two cysteines of the CXXC motif specific of the HMA family are boxed in red. Green dots highlight the positions in which the hydrophobic character is strongly conserved, corresponding to amino acids participating in the hydrophobic core of the ferredoxin fold. An additional β-strand, named β0, is predicted in the CoBaHMA sequences, including a strictly conserved histidine. B. Model of the CoBaHMA 3D structure, illustrated here with the *Synechococcus* sp. RS9917 sequence.

The C-terminal domain of the different calcyanin types consisted in three repetitions of a ∼50 amino acid motif, which was largely apolar and displayed a constant periodicity in hydrophobic and small (glycine/alanine) amino acids (supplementary fig. 4). We called this motif “GlyZip” in reference to the name proposed by (Kim et al. 2005) to describe recurrent, short Gly-X(3)-Gly-X(3)-Gly motifs allowing tight packing of transmembrane helices (Senes et al. 2004).

However, the calcyanin GlyZip motifs, organized around a central, highly conserved Gly-Pro dipeptide, were much longer (2×6 basic Gly-X(3)-Gly units) than those already known at the 3D level, which generally contained no more than three such units (Leonov and Arkin 2005). Moreover, they did not share any obvious sequence similarity with known domains, suggesting that these repeated motifs form a novel architecture. The repeated presence of glycine and hydrophobic amino acids every four amino acid residues over a large sequence length, with an unusual persistence of this periodic motif along evolution (especially for the first repeat) suggest that it may form compact and highly constrained assemblages of helices compatible with a membrane-embedded structure. These assemblages might resemble homo-oligomeric structures formed by short subunits, such as the c-rings of sodium-translocating ATP synthases (Kuehlbrandt 2019), which share similar, albeit smaller, glycine zippers. Analysis of multiple sequence alignments (supplementary fig. 4) allowed discriminating each of the three GlyZip calcyanin motifs based on specific signatures, including the presence of aromatic and polar amino acids, outside the repeated patterns. In particular, a tryptophan and a glutamic acid were strictly conserved in the third GlyZip motif in all calcyanin sequences. The second GlyZip motif of several calcyanin sequences matched part of a family model called PdsO (sortase-associated OmpA-like protein), found in, e.g., *Shewanella oneidensis* (see CDD annotations in supplementary table 2). The matching region, located before the OpmA-like C terminal domain, shows the typical features of a GlyZip unit (supplementary fig. 5) and is present as a single copy in PdsO, suggesting that this base unit evolved within calcyanin by triplication and enrichment in polar amino acids (see below). Last, among Y-type calcyanins, only that of *Fischerella* sp. NIES-4106 possessed all three GlyZip motifs, while all others, including in iACC-forming strains, contained only the first and third GlyZip ones, suggesting that calcyanins with only two GlyZip motifs remain functional (supplementary fig. 4). Interestingly, although it did not match the characteristic GlyZip profile, the duplicated domain found in the N-terminal region of Y-type calcyanins was also largely apolar and rich in small amino acids so as in GlyZip motifs.

### Calcyanin is involved in Ca homeostasis

In *C. fritschii* PCC 9212 and PCC 6912, the genes located directly upstream and downstream of *ccyA* were annotated as encoding a Ca(2+)/H(+) antiporter and a Na(+)-dependent bicarbonate transporter BicA, respectively (supplementary table 4). This is particularly interesting when considering that bicarbonates and calcium are obvious crucial ingredients for the synthesis of CaCO_3_. By searching homologs of these two transporters in a dataset of 602 cyanobacterial genomes, we observed that their combined presence was significantly associated with that of *ccyA* (chi2 test, p-value=1.4e-08; supplementary table 5). Indeed, all 35 genomes harboring *ccyA* had at least one copy of both genes, except *Synechococcus* sp. Lanier, which lacked BicA. The latter strain was also deviant from other *ccyA*-harboring strains based on very atypical N-ter and C-ter *ccyA* sequences. Since this strain was not available for EM analysis, we could not test if it contained iACC or not. By contrast, among the 567 genomes lacking *ccyA*, only 293 contained both transporter genes. Interestingly, in *Fischerella* sp. NIES-4106 megaplasmid, *ccyA* was located downstream a calcium/proton exchanger, in a region containing several additional genes potentially involved in biomineralization, such as two cation-transporting ATPases and a carbonic anhydrase (fig. 6). Overall, the correlation and/or co-localization of *ccyA* and genes involved in Ca or HCO3^-^transport and homeostasis supports the hypothesis of a functional role of calcyanin in Ca-carbonate biomineralization.

**Figure 6:**
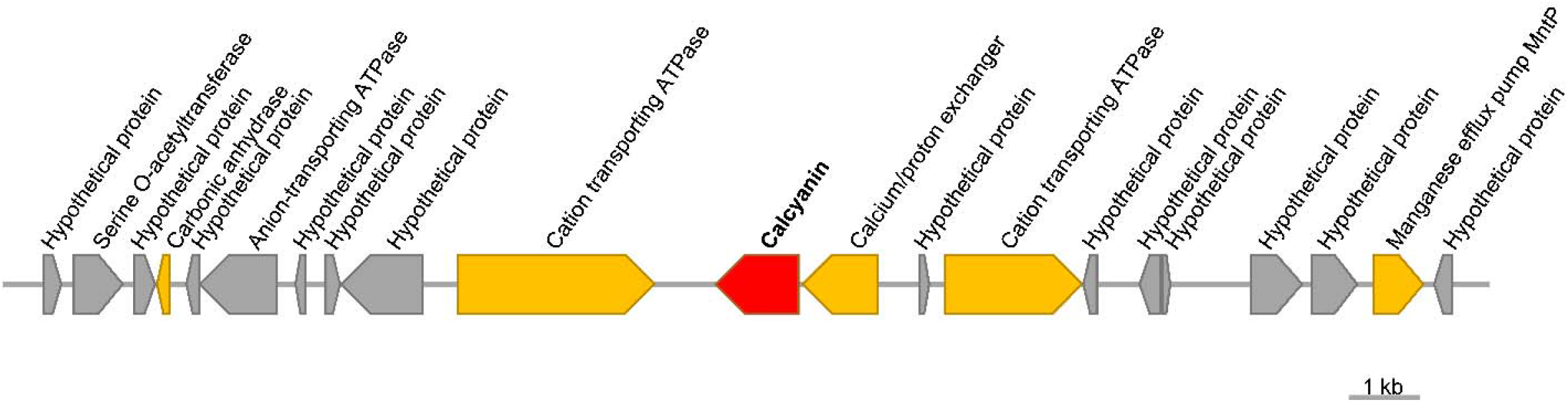
Genomic environment of *ccyA* in the megaplasmid of *Fischerella* sp. NIES-4106. Neighboring genes with functions potentially involved in biomineralization are indicated in orange. This megaplasmid sequence can be found in GenBank under accession number AP018299.1.

Attempts to obtain *ccyA* deletion mutants in the iACC-forming strains *Synechococcus* sp. PCC 6312 and *Cyanothece* sp. PCC 7425 were unsuccessful possibly because *ccyA* deletion was lethal in both strains, suggesting that this gene carries out an essential function in these cyanobacteria. Since this prevented a direct loss-of-function genetic analysis, we overexpressed the *ccyA* genes of the two evolutionary distant cyanobacteria *Synechococcus* sp. PCC 6312 and *G. lithophora* in the non-iACC-forming, but genetically manipulable host *Synechococcus elongatus* PCC 7942, which does not originally contain *ccyA* (supplementary table 6 and figure 6). Investigation of PCC 7942 cells overexpressing the *ccyA* gene of either *Synechococcus* sp. PCC 6312 or *G. lithophora* by EM-associated elemental chemical analyses did not show the presence of iACC but detected a significant Ca signal. In contrast, Ca was below detection limit in wild-type *Synechococcus elongatus* PCC 7942 cells (fig. 7). These results suggests that *ccyA* is functionally involved in Ca homeostasis, via a molecular process that remains to be fully elucidated.

**Figure 7.**
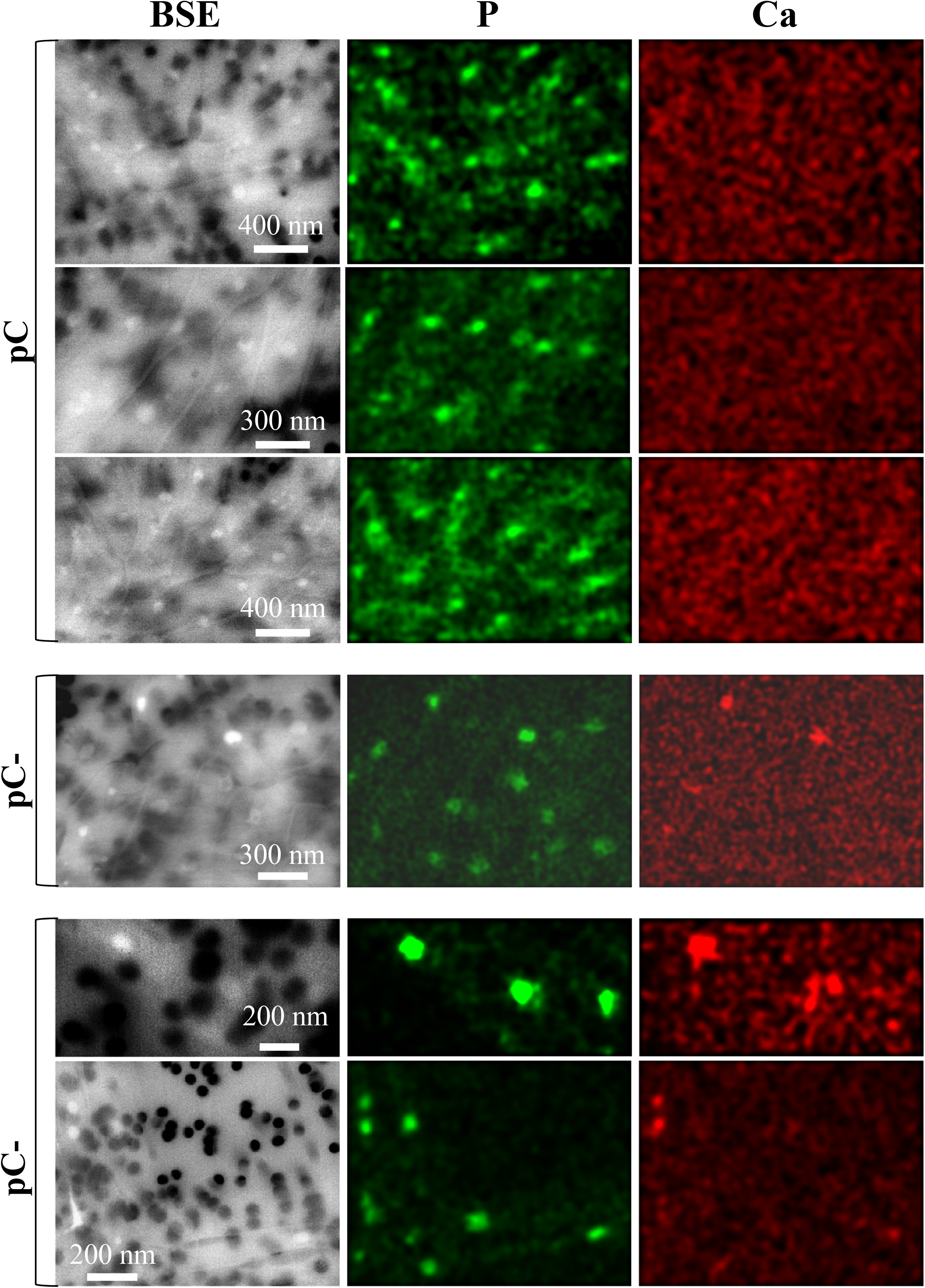
SEM analyses of mutants overexpressing ccyA. SEM-EDXS images (in backscattered electron (BSE) mode), P (green) and Ca (red) maps of *Synechochoccus elongatus* PCC 7942 mutants. The first three rows show cells of a *S. elongatus* PCC 7942 mutant harboring the empty pC plasmid. No Ca-rich inclusions are observed in these cells. In contrast, Ca-rich inclusions (polyphosphate) are observed in cells of *S. elongatus* PCC 7942 mutants harboring the plasmids pC-ccyA_Gloeo_ (fourth row) or pC-ccyA_S6312_ (fifth and sixth rows). See supplementary data 3 for details concerning the plasmid and strains.

### Phylogenetic distribution and evolution of calcyanin

Whatever the function of this diagnostic gene family, constructing its phylogeny allows to infer the possible evolutionary history of iACC biomineralization. We placed the species containing the *ccyA* gene on a general phylogeny of cyanobacteria constructed using 58 conserved proteins (supplementary table 7). The four calcyanin types were found in various lineages widely dispersed across this cyanobacterial tree (fig. 4a). Whereas the X, Y and Z types showed a distribution restricted to some particular clades (*Gloeomargarita, Fischerella* and closely related genera, and *Microcystis*, respectively), the CoBaHMA domain (i.e. W-type) was found in several distantly related branches (fig. 4a). Similarly, *ccyA* was detected in all the species of some clades (e.g., *Thermosynechococcus* clade), suggesting that it already existed in the genome of their last common ancestor, whereas it was missing in some species of other clades such as the *Chlorogloeopsis-Fischerella* one, suggesting several independent losses and/or horizontal gene transfer (HGT) events. To better characterize these evolutionary processes, we reconstructed the phylogeny of calcyanin using the conserved GlyZip domain sequences and compared it with the corresponding cyanobacterial species tree (fig. 4c). Despite a weaker resolution of the deep branches, reflecting the higher sequence variability of calcyanin, we retrieved the monophyly of most of the groups as found in the species tree, supporting the idea that *ccyA* was ancestral in these groups, and that the *ccyA-*lacking species, most likely lost it secondarily. In addition, we also identified *ccyA* HGT events between distant groups, the clearest case being that of the filamentous, heterocystous species *Scytonema millei*, which branched very close to the coccoidal, baeocystous species *Chroococcidiopsis thermalis* in the GlyZip tree, far from its expected relatives (*Fischerella* and *Mastigocladus*). Moreover, *S. millei* should possess a Y-type calcyanin because of its close phylogenetic relationship with *Fischerella*, instead of the CoBaHMA-type calcyanin that it actually harbors. Finally, the presence of *ccyA* in a megaplasmid of *Fischerella* sp. NIES-4106, also suggested that this gene could be mobilized between different species (fig. 6).

The overall congruence between the calcyanin tree and the species tree, both retrieving the monophyly of several large cyanobacterial clades (fig. 4c), supported a very ancient origin of *ccyA* in cyanobacteria, with independent losses in various lineages and some HGT cases, as found in *S. millei*. The alternative scenario of a recent origin of *ccyA* in one group followed by its transfer to the rest by HGT was unlikely given the extreme divergence of the N-terminal domains among the different types of calcyanin (fig. 4b). Because of its larger phylogenetic distribution, the CoBaHMA-type seemed to be the most ancient calcyanin version, while the Y-and Z-types have evolved in cyanobacterial groups that diverged more recently. The situation is less clear for the X-type due to its exclusive presence in *G. lithophora*, the so-far single representative species of the poorly known Gloeomargaritales. As mentioned above, the N-terminal domains of these four types of calcyanin did not share any apparent sequence similarity (fig. 4b). This could reflect either an extreme divergence from a common ancestral domain or the independent recruitment of non-homologous domains generating the different calcyanins by their fusion to the conserved GlyZip C-terminal domain.

## Conclusions

Here we show that the newly identified *ccyA* gene family, belonging to the genomic ‘dark matter’ of cyanobacteria, can be used as a diagnostic iACC biomineralization marker. The *ccyA*-encoded calcyanin protein has a unique architecture composed of highly divergent N-terminal domains fused with a novel, much more conserved GlyZip-containing C-terminal domain, which is likely to adopt an original, not yet described fold. Among the diverse N-terminal domains of calcyanin that we have identified here, the domain family CoBaHMA is found in the most widespread, and likely most ancient, calcyanin version. This domain family likely supports an as-yet undisclosed function within the HMA superfamily, associated with a patch of conserved basic amino acids. By tracking this gene in available genome databases, we uncovered a diversity of *ccyA*-bearing cyanobacteria capable of iACC biomineralization that is phylogenetically and environmentally much broader than previously thought, supporting a potential environmental significance. Moreover, the distribution and phylogeny of *ccyA* suggest that iACC biomineralization is ancient, with independent losses in various lineages and some HGT cases. Additional genes are likely involved in iACC formation but, unlike *ccyA*, they may not be specific to this function and/or they are not shared by all iACC-forming cyanobacteria. The specific distribution of *ccyA* in iACC-forming cyanobacteria, its correlated presence with bicarbonate and calcium transporters, and genetic analyses, all support a pivotal role of *ccyA* in iACC biomineralization. Further investigations are required to determine whether this function may involve the conserved glutamic acid residues of the C-ter domain, reminding Glu-rich proteins involved in ACC biomineralization (Aizenberg et al. 2002), or the basic amino acids in the N-ter domain, which may stabilize dense liquid phases of CaCO_3_ and delay the formation of ACC(Finney et al. 2020). Alternatively, calcyanin may have a more indirect role in iACC biomineralization serving as a cation transporter or a signaling molecule. In any case, iACC biomineralization clearly appears as an original case of controlled biomineralization in bacteria.

## Materials and Methods

### Identification of candidate iACC-specific orthologous groups

In a first step, the 56 genomic assemblies used to identify groups of orthologous genes specific to iACC-forming cyanobacteria (supplementary table 1) were retrieved from the NCBI database, except *Candidatus Synechococcus calcipolaris* PCC 11701 (D. Moreira, unpublished data). The 523 680 translated coding sequences derived from these genomes were processed using OrthoMCL with default settings (Li et al. 2003). This analysis included an all-vs-all blastp routine (E-value < 1e-05) and a clustering procedure into orthologous groups using the MCL algorithm.

### Iterative search for homologs of calcyanin in cyanobacterial genomes

Homologs of calcyanin were searched based on similarities of the conserved C-terminal domain in 594 available genomes of cyanobacteria using an iterative process. Our search dataset contained the protein sequences provided by the NCBI genome assembly records assigned to Cyanobacteria, published online before December 1st, 2017 (except the 6 identified in the first step, see above). For each genome assembly, we processed the first set of amino acid sequences available in the following ordered list (supplementary Data 2): (i) translated CDS or (ii) proteins in RefSeq annotation records, (iii) translated CDS or (iv) proteins in GenBank annotation records.

A multiple sequence alignment of the conserved C-terminal domain was built for the 6 calcyanin sequences identified in the first step (see above), using MAFFT (Katoh and Standley 2013). A HMM-profile was generated based on this alignment with the program hmmbuild from the HMMER package (version 3.3) (Eddy 2011). The options wblossum with wid 0.62 were used to downweight closely related sequences and upweight distantly related ones. To avoid biases towards glycine-rich unrelated proteins, we artificially reduced glycine weight by 20% in the profiles. The profile *vs* sequence similarity search was done with the program hmmsearch (E-value < 1.0e-70). The hits matching 100% of the profile length and corresponding to newly identified sequences were added to the new calcyanin dataset. The multiple sequence alignment and the HMM-profile of this dataset were then updated. These steps (alignment, building of HMM-profile, similarity search) were repeated until no new sequence was detected. In order to detect remote homologs of calcyanin, seven iterations of the entire process were done as described in supplementary table 8, with a progressive decrease of the stringency of the similarity search. In the beginning, we set a very low E-value and high cover to the profile. As the iterations proceeded, we increased the E-value and decreased the cover to the profile down to 70%. This cover threshold higher than 66% was designed to avoid (Gly)2 (instead of (Gly)3) to be matched. At the end of the whole process, we used the final HMM profile (supplementary Data 3) to search for similarity in the GenBank records of the processed genomic assemblies.

Last, *ccyA* was searched in the newly sequenced genomes of *Synechococcus* sp. PCC 6716 and PCC 6717 using tBLASTn with all previously identified *ccyA* sequences as queries. The CDS boundaries of the best BLAST hits were further assessed using Prodigal (Hyatt et al. 2010).

### Comparative genomics of C. fritschii PCC 6912 (no iACC observed) and C. fritschii PCC 9212 (with iACC)

The search of homologous genes shared by *C. fritschii* PCC 6912 and PCC 9212 genomes was achieved based on unidirectional BLASTp best hits as implemented in the PATRIC proteome comparison tool (Gillespie et al. 2011) (E-value < 1.0e-05, sequence coverage > 30%). For each genome assembly, we used the set of translated CDS as provided in the RefSeq annotation record. Gene functional categories were searched in COG database (v1) using CD-search (Marchler-Bauer and Bryant 2004) (E-value < 1.0e-05). The nucleotide sequences of the *ccyA*-containing contigs of *C. fritschii* PCC 6912 and PCC 9212 (NCBI accessions NZ_AJLN01000033.1 and NZ_AJLM01000017.1, 97,542 bp and 97,528 bp, respectively) were compared using BLASTn.

### Search for homologs of the Ca(2+)/H(+) antiporter and BicA Na(+)-dependent bicarbonate transporter in cyanobacterial genomes

Homologs of the Ca(2+)/H(+) antiporter and the Na(+)-dependent bicarbonate transporter BicA, encoded in *C. fritschii* PCC 6912 and PCC 9212 by the genes located upstream and downstream of *ccyA*, respectively, were searched using BLASTp (E-value < 1.0e-10) in a dataset of 602 cyanobacterial genomes. Owing to the incompleteness of the *ccyA*-upstream gene in the genomic sequence of these two strains, we used the most similar full-length sequence as Ca(2+)/H(+) antiporter query (96% identity; accession WP_016868870.1, *Fischerella muscicola* PCC 7414). BicA homologs were identified using the protein sequence from *C. fritschii* PCC 6912 and PCC 9212 as query (accession WP_016872894.1).

### Calcyanin functional annotation and structure prediction

The structural features of calcyanins were explored based on the information provided by amino acid sequences using Hydrophobic Cluster Analysis (HCA) (Callebaut et al. 1997; Bitard-Feildel et al. 2018). HCA provides a global view of the protein texture, with insights into the structural features of foldable regions (Bitard-Feildel et al. 2018). Similarities between domains composing calcyanin and known domains/3D structures were searched against different databases (NCBI nr sequence database, Conserved Domain Database (Yang et al. 2020) or the Protein Data Bank (PDB)) using tools for profile-sequence and profile-profile comparison such as PSI-BLAST(Altschul et al. 1997) and HH-PRED (Zimmermann et al. 2018), respectively. 3D structure modelling and visualization were performed using Modeller 9.23 (Webb and Sali 2016) and UCSF Chimera (Pettersen et al. 2004), respectively. Models were built using as templates the experimental 3D structures of the HMA, iHMA and YAM. The position of strand β0 was moreover putatively assigned with reference to the 3D structure of KipI (pdb 2KWA), based on the results of HH-PRED searches and subsequent superimposition of the 3D corresponding 3D structures (pdb 2RU9 and 2KWA, root mean square value of 2.1 Å on 55 Cα superimposed positions). Multiple sequence alignment handling and rendering were made using SeaView (Gouy et al. 2010) and EsPript (Robert and Gouet 2014), respectively.

### Molecular phylogenetic analyses

Phylogeny of cyanobacteria using different sets of species was reconstructed using 58 conserved proteins (Moreira et al. 2017). Each individual protein was aligned using MAFFT with the accurate L-INS-I option (Katoh and Standley 2013) and poorly aligned regions were removed with trimAl –automated1 (Capella-Gutiérrez et al. 2009). Trimmed alignments were concatenated to produce a supermatrix and maximum likelihood phylogenetic trees were reconstructed with the program IQ-Tree using the mixture model LG+C60+F+G (Nguyen et al. 2015). Statistical support was estimated using 1000 bootstrap replicates. The phylogeny of calcyanin was studied using the manually curated alignment of the conserved GlyZip C-terminal domain. A maximum likelihood tree was constructed with the program IQ-Tree using the mixture model LG+C20+F+G (Nguyen et al. 2015). Statistical support was estimated using 1000 bootstrap replicates.

### Electron microscopy analyses of iACC

Strains recovered from collections as well as mutants strains of *Synechococcus elongatus* PCC 7942 were analyzed by scanning transmission electron microscopy (STEM) for iACC search and/or Ca enrichment using a field emission gun JEOL-2100F microscope operating at 200 kV, equipped with a JEOL detector with an ultrathin window allowing detection of light elements. STEM allowed Z-contrast imaging in the HAADF mode. Compositional maps of Ca, P, and C were acquired by performing EDXS analysis in the STEM HAADF mode. For these analyses, a total of 0.5 mL of cultures was centrifuged at 8,000 × g for 10 min. Pellets were rinsed three times in Milli-Q (mQ) water (Millipore). After the final centrifugation, pellets were suspended in 200 μL of mQ water. A drop of 5 μL was deposited on a glow discharged carbon-coated 200-mesh copper grid and let dry at ambient temperature.

### Genetics

The pC-*ccyA*_Gloeo_ and pC-*ccyA*_S6312_ plasmids were derivatives of the RSF1010-derived pC vector (Veaudor et al. 2018) replicating in *E. coli* (supplementary table 6 and supplementary figure 6). They were transferred to *Synechococcus elongatus* PCC 7942 by trans-conjugation (Mermet-Bouvier and Chauvat 1994), using the improved triparental-mating protocol that follows. Overnight-grown cultures of the *E. coli* strains CM404, which propagates the self-transferable mobilization vector pRK2013, and TOP10, which propagates either pC, pC-*ccyA*_Gloeo_ or pC-*ccyA*_S6312_, were washed twice and resuspended in LB medium (1 × 10^9^ cells.mL^− 1^). Meanwhile, *S. elongatus* PCC 7942 mid-log phase cultures grown in mineral growth medium (MM) were centrifuged and concentrated five times (about 1× 10^8^ cells/mL) in fresh MM. Then, 100 μL of *S. elongatus* PCC 7942 cells were mixed with 30 μL of CM404 cells and 30 μL of TOP10 cells harboring either pC, pC-*ccyA*_Gloeo_ or pC-*ccyA*_S6312_. Thirty microliter aliquots of this mixture were spotted onto MM solidified with 1% agar (Difco), and incubated for 48 h under standard temperature (30°C) and light conditions (2500 lux, i.e. 31 μE.m^−2^.s^−1^). Then, cells were collected from each plate and suspended into 50 μL of liquid MM, prior to plating onto MM containing 5 μg.mL^-1^ of each the streptomycin (Sm) and spectinomycin (Sp) selective antibiotics. After about 10 days of incubation under standard light and temperature, Sm^R^Sp^R^ resistant conjugant clones were collected and re-streaked onto selective plates, prior to analyzing their plasmid content by PCR and DNA sequencing (Eurofins Genomics) using specific primers (supplementary Data 1).

## Acknowledgments

We thank Alexis De Wever and Marine Blondeau for helping acquiring some TEM data. We thank Mélanie Poinsot for helping in the preparation of some samples for transmission electron microscopy analyses. We thank Eva Jahodarova for shipping a culture of *Neosynehcococcus. sphagnicola*. This work was supported by the Agence Nationale de la Recherche (ANR Harley, ANR-19-CE44-0017-01), and the European Research Council under the European Union’s Seven Framework Program: ERC grants Calcyan (PI: K. Benzerara, Grant Agreement no. 307110) and PlastEvol (PI: D. Moreira, Grant Agreement no. 787904). Sigrid Görgen PhD grant was funded by the Sorbonne Université doctoral program Interfaces pour le Vivant.

## Author Contributions

KBE, EDU, CCC, FCH, DMO, PLG, ICA conceived and designed the work. KBE, EDU, TBF, GCA, CCC, MDE, IDI, GGA, MGU, SGO, FSP, DMO, ICA acquired, analysed and/or interpreted data. KBE, EDU, CCC, FCH, MGU, PLG, DMO, ICA drafted the work or substantively revised it.

## Data availability

Further information and requests for resources and reagents should be directed to and will be fulfilled by the lead contact, Karim Benzerara.

- Plasmids and mutant strains generated in this study are available upon request to the lead contact.
- Full datasets and codes will be uploaded to Zenodo upon acceptance for publication. DOIs are listed in the key resources table.
- The genomic assemblies of *Candidatus* Synechococcus calcipolaris PCC 11701 and *Synechococcus* sp. PCC 6716 and 6717 have been submitted on GenBank and are publicly available as of the date of publication. Accession numbers are listed in the key resources table.
- TEM-EDXS and SEM-EDXS data will be deposited on the Zenodo repository and will be publicly available as of the date of publication. DOIs will be listed in the key resources table.

## Supporting Information

### CONTENTS

#### 1. Supplementary Data (Separate files)

**Supplementary Data 1:** Nucleotide and aminoacid sequences for genetics

**Supplementary Data 2**: List of NCBI ftp server URL for downloading the 599 public proteomes of Cyanobacteria used in this study. These proteomes were published online before December 1st, 2017.

**Supplementary Data 3**: Final HMM profile generated at the end of the iterative search for homologs of calcyanin in cyanobacterial genomes.

#### 2. Tables (Separate files)

**Supplementary Table 1**: Strain and genomic features of the 56 organisms used to identify groups of orthologous genes specific to iACC-forming cyanobacteria.

**Supplementary Table 2**: Description of the 35 calcyanin sequences identified in this study. For each sequence: homology detection method, genomic and protein accessions, protein length, N-ter domain type, culture and environmental conditions. Strains with and without observed iACC are highlighted in green and red, respectively. Available functional annotations (RefSeq, CDD) are listed in the second tab.

**Supplementary Table 3**: Comparison of the genomic repertoires of *C. fritschii* PCC 6912 (no iACC observed) and *C. fritschii* PCC 9212 (with iACC) using the PATRIC proteome comparison tool. Each line corresponds to a pair of homologous genes (identified either as unidirectional or bidirectional blastp best hit) or to a singleton in one genome, i.e. without homolog in the other. The pair of ccyA genes is highlighted in yellow (line number 329).

**Supplementary Table 4** Protein markers used for the species tree reconstruction; accession numbers are from *Gloeomargarita lithophora*.

**Supplementary Table 5**: Genomic position and annotations (as provided by RefSeq and PATRIC) of the genes located directly upstream and downstream of ccyA, in the same strand, in *C. fritschii* PCC 6912 and PCC 9212.

**Supplementary Table 6**: Characteristics of the plasmids used in this study

**Supplementary Table 7**: Number of copies and sequence accession of Ca(2+)/H(+) antiporter and Na(+)-dependent bicarbonate transporter BicA found in the 602 cyanobacterial genomes of our complete dataset.

**Supplementary Table 8**: Step-dependent parameters of the iterative similarity search of the *ccyA* gene in cyanobacterial genomes, corresponding to a progressive decrease of the stringency.

#### 3. Supplementary Figures (below)

**Supplementary Figure 1:**
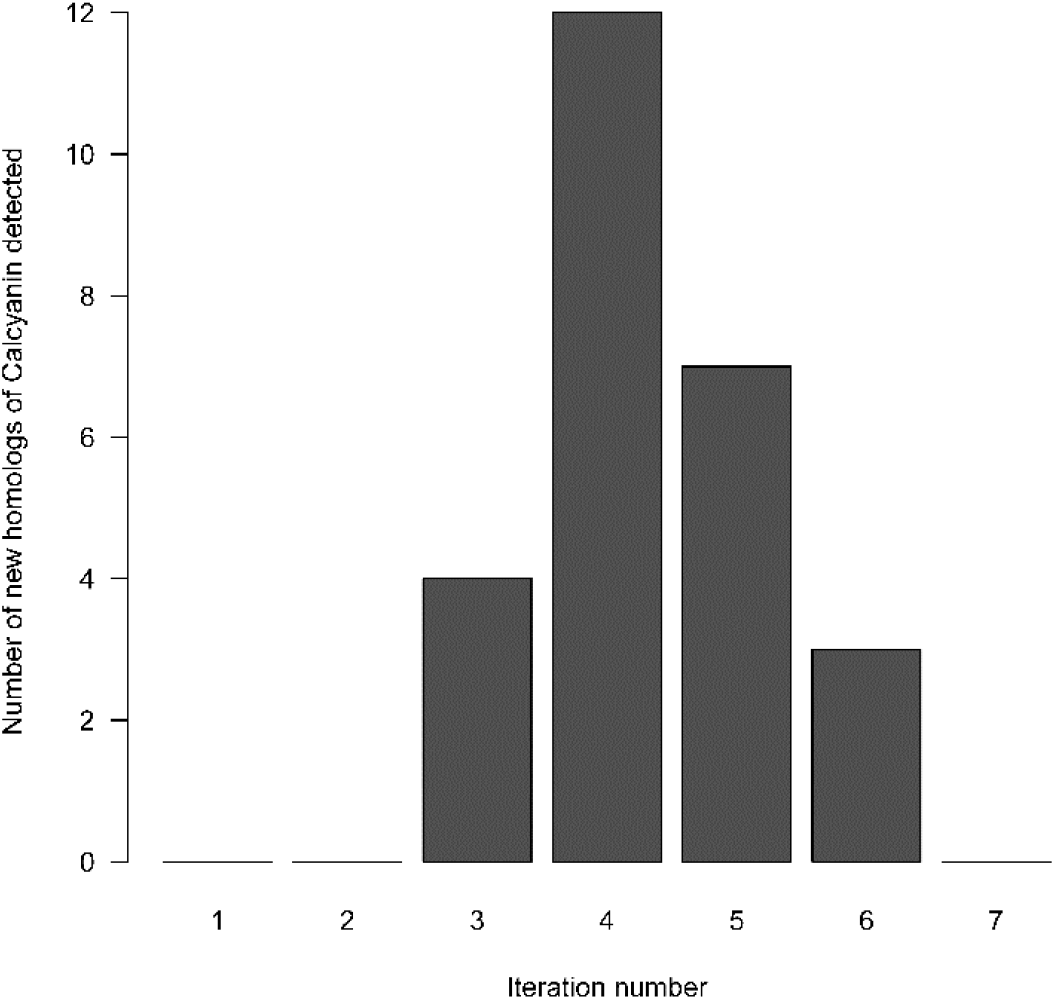
Number of calcyanin sequences detected at each step (or iteration) of the iterative search in cyanobacterial genomes.

**Supplementary Figure 2:**
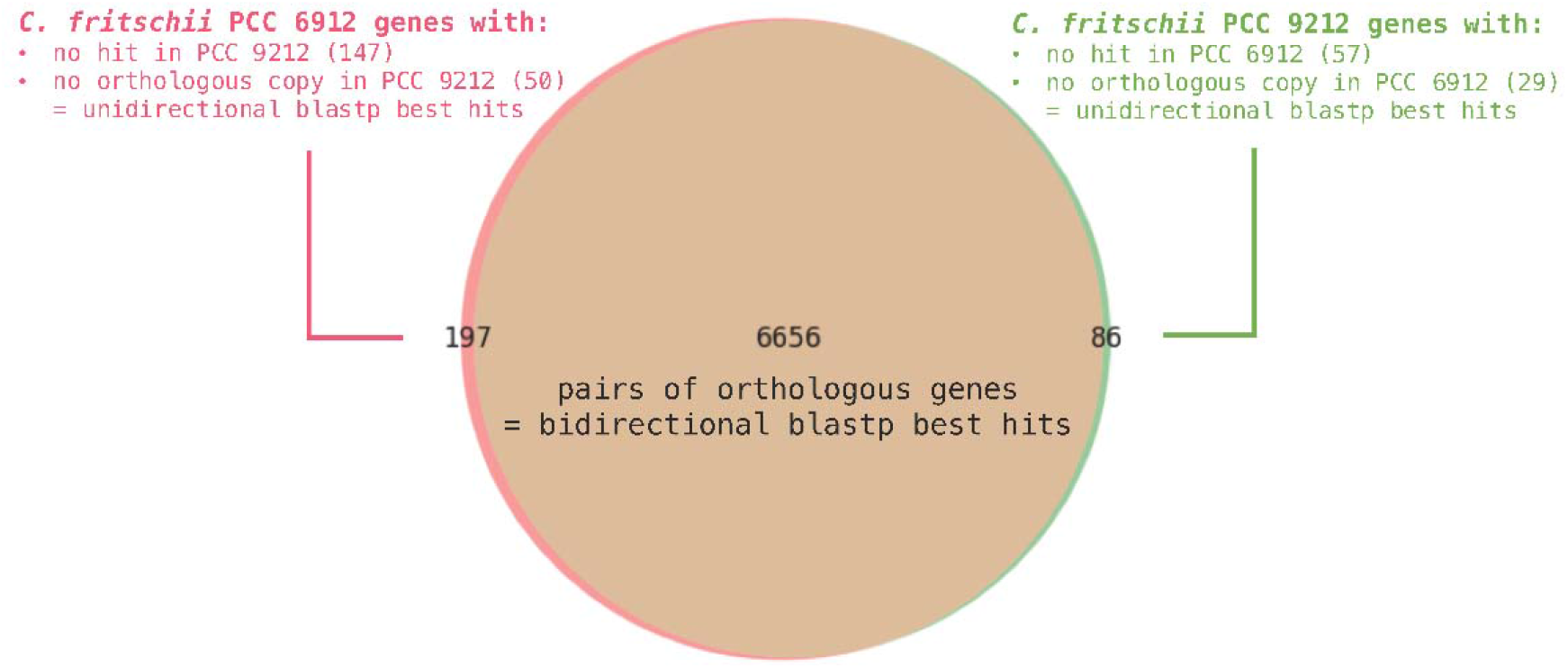
Venn diagram showing the overlap of the proteomes of *C. fritschii* PCC 6912 (no iACC observed) and *C. fritschii* PCC 9212 (with iACC), as processed using the PATRIC proteome comparison tool.

**Supplementary Figure 3:**
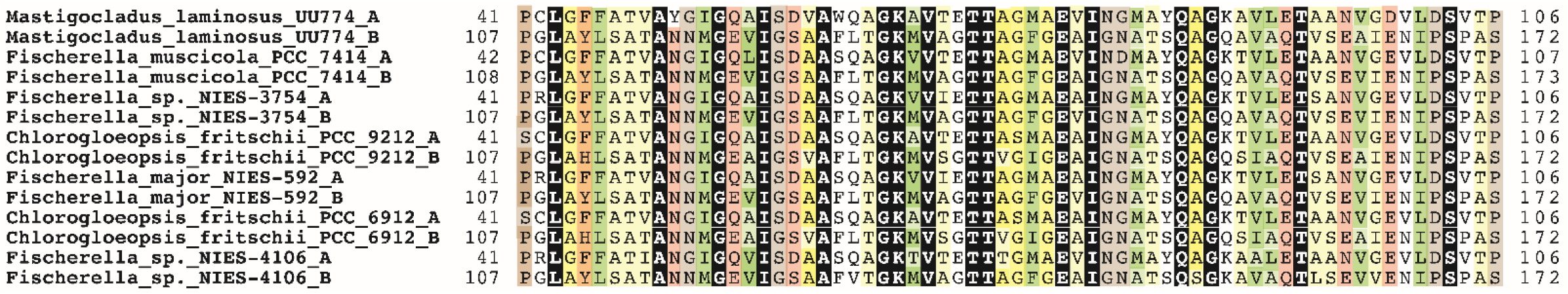
Sequence alignment of the duplicated domain found in the N-ter of Y-type calcyanins. The Y N-terminal domain consists of a twice-repeated sequence that is shown here. Identical amino acids are shown in white on a black background. Positions with conserved features are colored according to amino acid properties: green for hydrophobic (dark for strong hydrophobic amino acids such as V, I, L, F, M, Y, W; light for other amino acids, which can also occupy such positions such as A, T, S, C); orange for aromatic amino acids (F, Y, with possible substitutions in H); pink for polar, non-basic amino acids (D, E, Q, N, S, T); yellow for small amino acids, with positions mainly occupied by G or A (darker yellow); grey or brown for loop-forming amino acids (P, G, D, N, S), with positions mainly occupied by P highlighted darker. This repeated sequence is particularly rich in small amino acids, reminiscent of the global composition of the GlyZip motifs, although the periodicity of small amino acids every four residues is less pronounced.

**Supplementary Figure 4:**
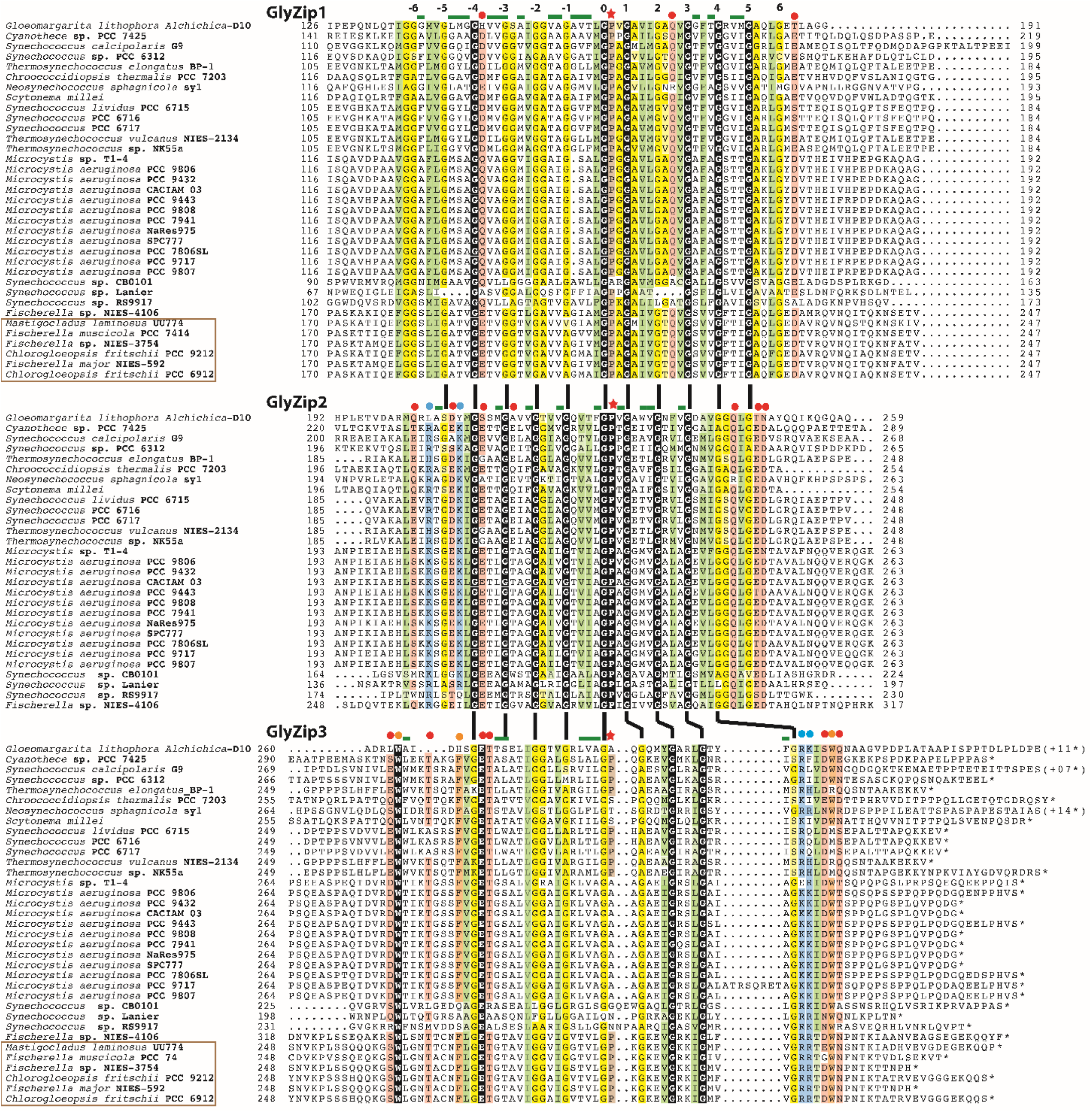
Sequence alignment of the calcyanin GlyZips. Identical amino acids are shown in white on a black background; similarities are colored according to amino acid properties: (i) green: hydrophobic (dark for strong hydrophobic amino acids (V, I, L, F, M, Y, W); light for other amino acids, which can also occupy such positions (A, T, S, C)); (ii) yellow: small amino acids, with positions mainly occupied by G or A highlighted darker; (iii) orange: aromatic amino acids (W, F, Y), (iv) pink: polar, non-basic amino acids (D, E, Q, N, S, T); brown: loop-forming amino acids (P, G, D, N, S), with positions mainly occupied by P highlighted darker. Positions for which a globally non polar character is observed are highlighted with green bars above the alignment, in addition to those for which a strong hydrophobic character is observed (boxed green). The names of the sequences from the Y family which do not possess the GlyZip2 motif are boxed. The periodic motif involved glycine (or a small amino acid) residues every four residue is highlighted with numbers positioned relative to the central conserved proline (0), and can be followed from one GlyZip motif to another one with vertical bars.

**Supplementary Figure 5:**
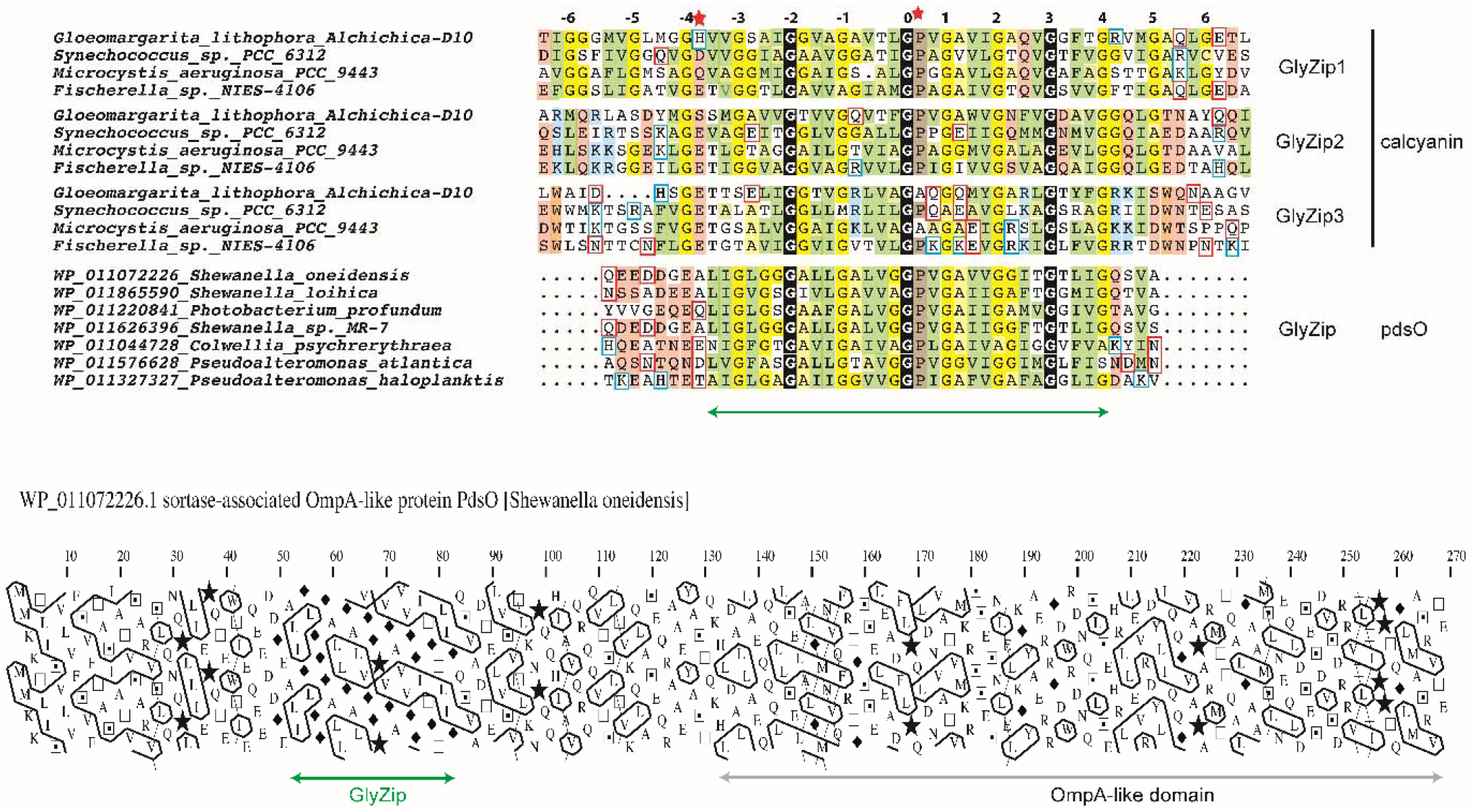
A completely apolar, single GlyZip in pdsO proteins. The GlyZip sequences of one member of each of the calcyanin groups are aligned with the conserved, apolar motif found in the N-terminal pdsO proteins (delineated with an arrow). See caption of Supplementary Fig. 6 for the color code. Polar amino acids are either colored (when conserved in a group of sequences) or boxed (T and S are considered as polar only in positions in which other polar amino acids are also found).

**Supplementary Figure 6:**
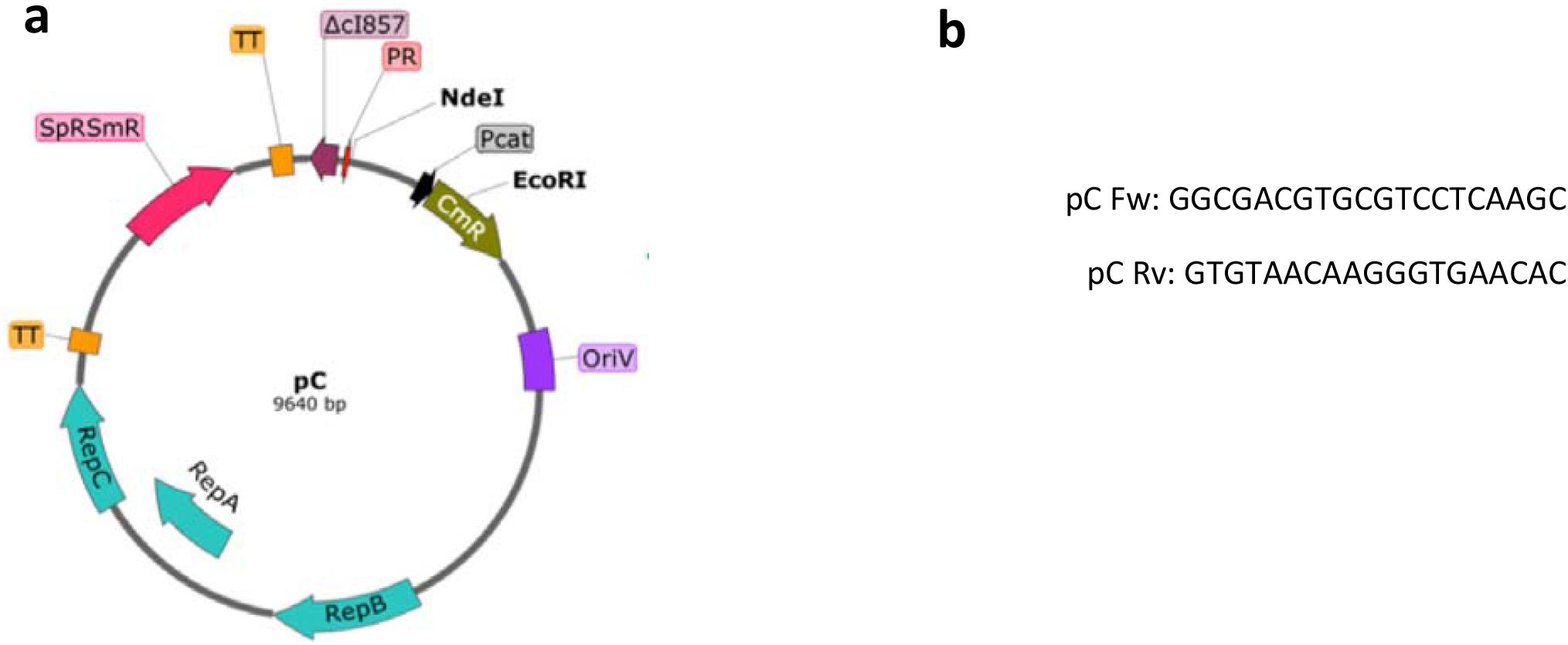
a) Structure of the pC plasmid, i.e. a RSF1010-derived plasmid vector (Sp^R^/Sm^R^,Cm^R^) harboring the strong constitutive *p*R promoter (Veaudor et al. 2018). B) Sequence of the primers used for sequencing *Synechococcus* sp. PCC 6312 and *Gloeomargarita lithophora ccyA* genes cloned into the pC-derived plasmids.

## Notes

### Competing Interest Statement

The authors have declared no competing interest.

## References

Aizenberg J, Lambert G, Weiner S, Addadi L. 2002. Factors involved in the formation of amorphous and crystalline calcium carbonate: a study of an ascidian skeleton. J Am Chem Soc 124:32–39.

Altermann W, Kazmierczak J, Oren A, Wright DT. 2006. Cyanobacterial calcification and its rock-building potential during 3.5 billion years of Earth history. Geobiology 4:147–166.

Altschul SF, Madden TL, Schäffer AA, Zhang J, Zhang Z, Miller W, Lipman DJ. 1997. Gapped BLAST and PSI-BLAST: a new generation of protein database search programs. Nucleic Acids Res 25:3389–3402.

Benzerara K, Skouri-Panet F, Li J, Férard C, Gugger M, Laurent T, Couradeau E, Ragon M, Cosmidis J, Menguy N, et al. 2014. Intracellular Ca-carbonate biomineralization is widespread in cyanobacteria. Proc Natl Acad Sci U S A 111:10933–10938.

Bitard-Feildel T, Lamiable A, Mornon J-P, Callebaut I. 2018. Order in disorder as observed by the “Hydrophobic Cluster Analysis” of protein sequences. Proteomics 18:e1800054.

Blondeau M, Benzerara K, Ferard C, Guigner J-M, Poinsot M, Coutaud M, Tharaud M, Cordier L, Skouri-Panet F. 2018. Impact of the cyanobacterium Gloeomargarita lithophora on the geochemical cycles of Sr and Ba. Chemical Geology 483:88–97.

Blondeau M, Sachse M, Boulogne C, Gillet C, Guigner J-M, Skouri-Panet F, Poinsot M, Ferard C, Miot J, Benzerara K. 2018. Amorphous Calcium Carbonate Granules Form Within an Intracellular Compartment in Calcifying Cyanobacteria. Front. Microbiol. 9:1768.

Blue CR, Giuffre A, Mergelsberg S, Han N, De Yoreo JJ, Dove PM. 2017. Chemical and physical controls on the transformation of amorphous calcium carbonate into crystalline CaCO3 polymorphs. Geochim. Cosmochim. Acta 196:179–196.

Bradley JA, Daille LK, Trivedi CB, Bojanowski CL, Stamps BW, Stevenson BS, Nunn HS, Johnson HA, Loyd SJ, Berelson WM, et al. 2017. Carbonate-rich dendrolitic cones: insights into a modern analog for incipient microbialite formation, Little Hot Creek, Long Valley Caldera, California. npj Biofilms Microbomes 3:32.

Bull PC, Cox DW. 1994. Wilson disease and Menkes disease: new handles on heavy-metal transport. Trends Genet 10:246–252.

Callebaut I, Labesse G, Durand P, Poupon A, Canard L, Chomilier J, Henrissat B, Mornon JP. 1997. Deciphering protein sequence information through hydrophobic cluster analysis (HCA): current status and perspectives. Cell Mol Life Sci 53:621–645.

Cam N, Benzerara K, Georgelin T, Jaber M, Lambert J-F, Poinsot M, Skouri-Panet F, Cordier L. 2016. Selective uptake of alkaline earth metals by cyanobacteria forming intracellular carbonates. Environ. Sci. Technol. 50:11654–11662.

Cam N, Benzerara K, Georgelin T, Jaber M, Lambert J-F, Poinsot M, Skouri-Panet F, Moreira D, López-García P, Raimbault E, et al. 2018. Cyanobacterial formation of intracellular Ca-carbonates in undersaturated solutions. Geobiology 16:49–61.

Capella-Gutiérrez S, Silla-Martínez JM, Gabaldón T. 2009. trimAl: a tool for automated alignment trimming in large-scale phylogenetic analyses. Bioinformatics 25:1972–1973.

Couradeau E, Benzerara K, Gerard E, Moreira D, Bernard S, Brown GE, Lopez-Garcia P. 2012. An Early-Branching Microbialite Cyanobacterium Forms Intracellular Carbonates. Science 336:459–462.

De la Concepcion JC, Franceschetti M, Maqbool A, Saitoh H, Terauchi R, Kamoun S, Banfield MJ. 2018. Polymorphic residues in rice NLRs expand binding and response to effectors of the blast pathogen. Nature Plants 4:576–585.

De Wever A, Benzerara K, Coutaud M, Caumes G, Poinsot M, Skouri-Panet F, Laurent T, Duprat E, Gugger M. 2019. Evidence of high Ca uptake by cyanobacteria forming intracellular CaCO3 and impact on their growth. Geobiology 17:676–690.

Eddy SR. 2011. Accelerated Profile HMM Searches. PLOS Computational Biology 7:e1002195.

Finney AR, Innocenti Malini R, Freeman CL, Harding JH. 2020. Amino acid and oligopeptide effects on calcium carbonate solutions. Crystal Growth & Design 20:3077–3092.

Frangeul L, Quillardet P, Castets A-M, Humbert J-F, Matthijs HCP, Cortez D, Tolonen A, Zhang C-C, Gribaldo S, Kehr J-C, et al. 2008. Highly plastic genome of Microcystis aeruginosa PCC 7806, a ubiquitous toxic freshwater cyanobacterium. BMC Genomics 9:274.

Gillespie JJ, Wattam AR, Cammer SA, Gabbard JL, Shukla MP, Dalay O, Driscoll T, Hix D, Mane SP, Mao C, et al. 2011. PATRIC: the comprehensive bacterial bioinformatics resource with a focus on human pathogenic species. Infect Immun 79:4286–4298.

Gouy M, Guindon S, Gascuel O. 2010. SeaView version 4: A multiplatform graphical user interface for sequence alignment and phylogenetic tree building. Mol Biol Evol 27:221–224.

Humbert J-F, Barbe V, Latifi A, Gugger M, Calteau A, Coursin T, Lajus A, Castelli V, Oztas S, Samson G, et al. 2013. A tribute to disorder in the genome of the bloom-forming freshwater cyanobacterium Microcystis aeruginosa. PLoS One 8:e70747.

Hyatt D, Chen G-L, LoCascio PF, Land ML, Larimer FW, Hauser LJ. 2010. Prodigal: prokaryotic gene recognition and translation initiation site identification. BMC Bioinformatics 11:119.

Jiang D, Zhao Y, Fan J, Liu X, Wu Y, Feng W, Zhang XC. 2014. Atomic resolution structure of the E. coli YajR transporter YAM domain. Biochem Biophys Res Commun 450:929–935.

Jiang D, Zhao Y, Wang X, Fan J, Heng J, Liu X, Feng W, Kang X, Huang B, Liu J, et al. 2013. Structure of the YajR transporter suggests a transport mechanism based on the conserved motif A. Proc Natl Acad Sci U S A 110:14664–14669.

Katoh K, Standley DM. 2013. MAFFT Multiple Sequence Alignment Software Version 7: Improvements in Performance and Usability. Mol Biol Evol 30:772–780.

Kim S, Jeon T-J, Oberai A, Yang D, Schmidt JJ, Bowie JU. 2005. Transmembrane glycine zippers: physiological and pathological roles in membrane proteins. Proc Natl Acad Sci U S A 102:14278–14283.

Komarek J, Johansen JR, Smarda J, Strunecky O. 2020. Phylogeny and taxonomy of Synechococcus-like cyanobacteria. Fottea 20:171–191.

Kuehlbrandt W. 2019. Structure and Mechanisms of F-Type ATP Synthases. In: Kornberg RD, editor. Annual Review of Biochemistry. Vol. 88. Palo Alto: Annual Reviews. p. 515–549.

Lamiable A, Bitard-Feildel T, Rebehmed J, Quintus F, Schoentgen F, Mornon J-P, Callebaut I. 2019. A topology-based investigation of protein interaction sites using Hydrophobic Cluster Analysis. Biochimie 167:68–80.

Latour D, Salençon M-J, Reyss J-L, Giraudet H. 2007. Sedimentary Imprint of Microcystis aeruginosa (cyanobacteria) Blooms in Grangent Reservoir (Loire, France). Journal of Phycology 43:417–425.

Lefevre CT, Bazylinski DA. 2013. Ecology, diversity, and evolution of magnetotactic bacteria. Microbiol. Mol. Biol. Rev. 77:497–526.

Leonov H, Arkin IT. 2005. A periodicity analysis of transmembrane helices. Bioinformatics 21:2604–2610.

Li L, Stoeckert CJ, Roos DS. 2003. OrthoMCL: Identification of Ortholog Groups for Eukaryotic Genomes. Genome Res. 13:2178–2189.

Marchler-Bauer A, Bryant SH. 2004. CD-Search: protein domain annotations on the fly. Nucleic Acids Res 32:W327–331.

Marron AO, Ratcliffe S, Wheeler GL, Goldstein RE, King N, Not F, de Vargas C, Richter DJ. 2016. The Evolution of Silicon Transport in Eukaryotes. Molecular Biology and Evolution 33:3226–3248.

Mehta N, Benzerara K, Kocar BD, Chapon V. 2019. Sequestration of radionuclides radium-226 and strontium-90 by cyanobacteria forming intracellular calcium carbonates. Environ. Sci. Technol. 53:12639–12647.

Mermet-Bouvier P, Chauvat F. 1994. A conditional expression vector for the cyanobacteria Synechocystis sp. strains PCC6803 and PCC6714 or Synechococcus sp. strains PCC7942 and PCC6301. Curr Microbiol 28:145–148.

Monteil CL, Benzerara K, Menguy N, Bidaud CC, Michot-Achdjian E, Bolzoni R, Mathon FP, Coutaud M, Alonso B, Garau C, et al. 2020. Intracellular amorphous Ca-carbonate and magnetite biomineralization by a magnetotactic bacterium affiliated to the Alphaproteobacteria. The ISME Journal:1–18.

Moreira D, Tavera R, Benzerara K, Skouri-Panet F, Couradeau E, Gerard E, Fonta CL, Novelo E, Zivanovic Y, Lopez-Garcia P. 2017. Description of Gloeomargarita lithophora gen. nov., sp nov., a thylakoid-bearing, basal-branching cyanobacterium with intracellular carbonates, and proposal for Gloeomargaritales ord. nov. Int. J. Syst. Evol. Microbiol. 67:653–658.

Nguyen L-T, Schmidt HA, von Haeseler A, Minh BQ. 2015. IQ-TREE: a fast and effective stochastic algorithm for estimating maximum-likelihood phylogenies. Mol Biol Evol 32:268–274.

Pettersen EF, Goddard TD, Huang CC, Couch GS, Greenblatt DM, Meng EC, Ferrin TE. 2004. UCSF Chimera-a visualization system for exploratory research and analysis. J Comput Chem 25:1605–1612.

Ragon M, Benzerara K, Moreira D, Tavera R, Lopez-Garcia P. 2014. 16S rDNA-based analysis reveals cosmopolitan occurrence but limited diversity of two cyanobacterial lineages with contrasted patterns of intracellular carbonate mineralization. Front. Microbiol. [Internet] 5. Available from: http://journal.frontiersin.org/article/10.3389/fmicb.2014.00331/abstract

Reynolds CS, Rogers DA. 1976. Seasonal variations in the vertical distribution and buoyancy of Microcystis aeruginosa Kütz. emend. Elenkin in Rostherne Mere, England. Hydrobiologia 48:17–23.

Riding R. 2012. A Hard Life for Cyanobacteria. Science 336:427–428.

Robert X, Gouet P. 2014. Deciphering key features in protein structures with the new ENDscript server. Nucleic Acids Res 42:W320–324.

Senes A, Engel DE, DeGrado WF. 2004. Folding of helical membrane proteins: the role of polar, GxxxG-like and proline motifs. Curr. Opin. Struct. Biol. 14:465–479.

Veaudor T, Ortega-Ramos M, Jittawuttipoka T, Bottin H, Cassier-Chauvat C, Chauvat F. 2018. Overproduction of the cyanobacterial hydrogenase and selection of a mutant thriving on urea, as a possible step towards the future production of hydrogen coupled with water treatment. PLoS One 13:e0198836.

Wang X, Zoccola D, Liew YJ, Tambutte E, Cui G, Allemand D, Tambutte S, Aranda M. 2021. The evolution of calcification in reef-building corals. Mol Biol Evol:msab103.

Webb B, Sali A. 2016. Comparative protein structure modeling using MODELLER. Curr Protoc Protein Sci 86:2.9.1-2.9.37.

Weiner S, Dove PM. 2003. An overview of biomineralization processes and the problem of the vital effect. In: Dove PM, DeYoreo JJ, Weiner S, editors. Biomineralization. Vol. 54. Chantilly: Mineralogical Soc Amer. p. 1–29.

Yang M, Derbyshire MK, Yamashita RA, Marchler-Bauer A. 2020. NCBI’s conserved domain database and tools for protein domain analysis. Curr Protoc Bioinformatics 69:e90.

Yarra T, Blaxter M, Clark MS. 2021. A bivalve biomineralization toolbox. Mol Biol Evol:msab153.

Zhao L, Song Y, Li L, Gan N, Brand JJ, Song L. 2018. The highly heterogeneous methylated genomes and diverse restriction-modification systems of bloom-forming Microcystis. Harmful Algae 75:87–93.

Zimmermann L, Stephens A, Nam S-Z, Rau D, Kübler J, Lozajic M, Gabler F, Söding J, Lupas AN, Alva V. 2018. A completely reimplemented MPI bioinformatics toolkit with a new HHpred server at its core. J Mol Biol 430:2237–2243.

