## Supplementary Data 1-3 for "A new gene family diagnostic for intracellular biomineralization of amorphous Ca-carbonates by cyanobacteria"

**Three sets of supplementary data (Sup data 1, sup data 2, sup data 3)**

**Supplementary Data 1: Nucleotide and aminoacid sequences for genetics**

**Data 1a. Nucleotide sequence of part of the pC-ccyA*_S6312_* plasmid used for the constitutive expression of the *Synechococcus* sp. PCC 6312 *ccyA* gene.** The lambda phage P*_R_* promoter sequence originating from the lambda phage *cro* gene is shown in red. It comprises the -35 box (5'-**TTGACT-3')**, the -10 box (5'-**GATAAT-3'**) and the transcription start site (**A**) as well as the Shine d'Algarno sequence (ribosome bind site; **aaggagg**). For further details, see (Mermet-Bouvier and Chauvat, 1994). The *ccyA* gene was recovered by PCR from *Synechococcus* PCC 6312 genome. For this purpose, we used the 6312_gene_Nde1_Fw and 6312_gene_EcoR1_Rv primers and cloned in the *Nde*I (**catATG**) and *EcoR*I (**gaattc**) restriction site of the pC plasmid (Veaudor et al. 2018) thereby truncating the chloramphenicol resistance encoding gene (Cm^S^). The coding sequence of *ccyA_S6312_* is highlighted in green.

**5’GGCGACGTGCGTCCTCAAGC**tgctcttgtgttaatggtttcttttttgtgctcatacgttaaatctatcaccgcaagggataaatatctaacaccgtgcgtg**TTGACT**attttacctctggcggt**GATAAT**ggttgc**A**tgtact**aaggagg**t**cat**ATGAGCAGCACATCCCCATCCTCTAGCCAGCCACCTCCAGGCCCCGATCCTGTTATCCAGGCTGAGCTCGTCCATCTCACTCCAGATCGGTTGCGGTTGAAAATCCCCCATCTCCGCCAAGACCCAGGCTACGGAACCTATCTCCAACAGCATCTCCAGGCCCAGACCGGGATCACCGAAGTTCGCCTCAACTCAACCGCCCAATCTTTAACCCTTCACTGGAACCCCCAAGTCATTTCCCTCCCGCAACTGTTAACCCAACTCCAGGCCATTGGGGACTTAGAGGTGATAGGTCAGGGAAATCATGGCATCACGAACCTCACCCGCGCCTTTGCCCTAGAACCAGAACAAGTCAGCGACAAAGCCCAGGATATTGGCAGTTTTATTGTTGGGGGACAAGTTGGAGATGTAGTCGGCGGGATAGCTGGGGCGGCAGTGGGTGGGGCCACAATCGGGCCAGCCGGGGTAGTTTTGGGGACACAGGTGGGGACGTTTGTAGGCGGTGTCATTGGGGCAAGAGTTTGTGTAGAGTCCATGCAAACCCTCAAGGAGCATGCCTTTGATCTGCAAACGCTTTGTTTAGATAAAACTAAAGAAAAGGTCACCCAAAGTTTAGAAATTCGCACCAGCAGTAAAGCCGGAGAGGTAGCTGGGGAAATCACGGGGGGCCTGGTGGGGGGAGCTTTGCTTGGGCCGCCCGGAGAAATTATTGGGCAAATGATGGGGAATATGGTCGGCGGACAAATTGCCGAAGACGCAGCCCGACAAGTCATCAGCCCTGATCCAAAACCTGACACAACAGCCCCGACATCGTCCTCGGTTAATATTGTTCTGGAATGGTGGATGAAAACGAGTCGCGCCTTTG AAGCAGGTAGTCGGGCCGGCCGAATCATTGATTGGAATACAGAGTCTGCTTCCTGTAAACAGCCCCAGTCAAACCAGGCTAAAACTGAGGAATTATAATCCAATAAGAACAGTTCAGGAATTTTGTTCGGAGTACTGAGACGATTGCG**gaattc**cgtatggcaatgaaagacggtgagctggtgatatgggatagtgttcacccttgttacac-3’

**Data 1b. Amino-acid sequence of the *Synechococcus* sp. PCC 6312 calcyanin encoded by the pC-ccyA*_S6312_* plasmid**

MSSTSPSSSQPPPGPDPVIQAELVHLTPDRLRLKIPHLRQDPGYGTYLQQHLQAQTGITEVRLNSTAQSLTLHWNPQVISLPQLLTQLQAIGDLEVIGQGNHGITNLTRAFALEPEQVSDKAQDIGSFIVGGQVGDVVGGIAGAAVGGATIGPAGVVLGTQVGTFVGGVIGARVCVESMQTLKEHAFDLQTLCLDKTKEKVTQSLEIRTSSKAGEVAGEITGGLVGGALLGPPGEIIGQMMGNMVGGQIAEDAARQVISPDPKPDTTAPTSSSVNIVLEWWMKTSRAFVGETALATLGGLLMRLILGPQAEAVGLKAGSRAGRIIDWNTESASCKQPQSNQAKTEEL

**Data 1c. Sequence of primers used for PCR amplification of the *Synechococcus* sp*.* PCC 6312 *ccyA* coding sequence by *Nde*I and *Eco*RI restriction sites (underlined) for cloning into the pC plasmid vector opened with the same enzymes.**

6312_gene_Nde1_Fw: CCTGGCAGTTGGCCCCTCCATATGAGCAGCACATCCCCATCCTCTA

6312gene_EcoR1_Rev:xATTACCGCTCTTTAGAGGGAATTCCGCAATCGTCTCAGTACTCCGAACAAA

**Data 1d. Sequence of primers used for sequencing the *ccyA*_S6312_ gene**

6312 Fw for sequencing of *ccyA*: ATGAGCAGCACATCCCCATCCTCT

6312 Rv for sequencing of *ccyA:* CGCAATCGTCTCAGTACTCCGAACAAA

**Data 1e.** **Nucleotide sequence of part of the pC-ccyA*_gloeo_* plasmid used for the constitutive expression of the *Gloeomargarita lithophora ccyA* gene. Sequence was adapted to the codon usage of the (non-iACC forming) model cyanobacterium *Synechococcus* *elongatus* PCC 7942. The lambda phage P*_R_* promoter, which comprises the -35 box (5'-TTGACT-3'), the -10 box (5'-GATAAT-3') and the transcription start site (A), and the ribosome binding sequence (5'-aaggagg-3') originating from the lambda phage *cro* gene is shown in red (Mermet-Bouvier and Chauvat, 1994). The *ccyA_Gloeo_* sequence flanked by the *Nde*I (catATG) and *Eco*RI (gaattc) restriction sites used for its cloning in the pC expression vector (see Table 1 above SX) is highlighted in green.**

**5’GGCGACGTGCGTCCTCAAGC**tgctcttgtgttaatggtttcttttttgtgctcatacgttaaatctatcaccgcaagggataaatatctaacaccgtgcgtg**TTGACT**attttacctctggcggt**GATAAT**ggttgc**A**tgtact**aaggagg**t**cat**ATGATGTTCCCCAAACCCGAGTTTCCCTGGTGGGAACAGACCCTGGAGGATCTGCAGAAGCTGGCTGAGCAGACGAAGTCTCAAATCACAGAGGCGATCACCAGCGTCCAGGATACGGCGACGCAAGCGCAGCATACCCTGGAGCAGACCGTGCAGCAGGTCGTGCCCCCGGAAGTGCAAGAACAGGTCCAAACCGCCGTCGCCACCAATCAAGTCCTGGTGAATCAGTGGGTGCACCGCGCCCAAGACACCGTGGCAAGTCAGATTATTGTCGTGGAAAAAATTTTCATCGAACTCGACCAGAGCCGCGTGGGCGTGACCCAACAGGTGACGGTGCGCCTGCAGGCCATCATCCCCGAACCCCAGAACCTGCAGACCATCGGCGGCGGCATGGTCGGTCTGATGGGCGGCCATGTCGTGGGCTCGGCCATCGGCGGCGTGGCGGGCGCCGTGACGCTGGGCCCGGTCGGCGCTGTGATCGGCGCCCAAGTGGGCGGCTTTACCGGCCGCGTCATGGGCGCCCAACTGGGTGAAACCCTGGCGGGCGGCCACCCCCTGGAGACGGTGGATGCCCGCATGCAACGGCTGGCCAGCGATTATATGGGCAGCAGCATGGGCGCGGTGGTGGGCACCGTAGTAGGTCAGGTGACATTTGGTCCGGTAGGAGCCTGGGTAGGTAACTTTGTAGGTGACGCCGTAGGGGGCCAATTAGGTACTAACGCATATCAGCAAATTAAACAGGGACAGGCCCAAGCTGATAGATTGTGGGCCATTGACCATAGCGGTGAAACGACGTCGGAGCTGATCGGCGGCACGGTCGGAAGACTAGTCGCTGGGGCTCAAGGGCAGATGTATGGCGCGCGACTGGGTACTTACTTCGGACGCAAAATTAGTTGGCAGAACGCTGCCGGCGTGCCGGACCCACTCGCCACCGCCGCCCCGATCAGTCCTCCCACTGATCTGCCACTAGATCCTGAGTTATGCTACCCTAAAGAGGAAGCCCCCCAGAATTAG**gaattc**cgtatggcaatgaaagacggtgagctggtgatatgggatagtgttcacccttgttacac-3’

**Data 1f. Amino-acid sequence of *Gloeomargarita lithophora* calcyanin encoded by the pC-ccyA*_Gloeo_* plasmid.**

MMFPKPEFPWWEQTLEDLQKLAEQTKSQITEAITSVQDTATQAQHTLEQTVQQVVPPEVQEQVQTAVATNQVLVNQWVHRAQDTVASQIIVVEKIFIELDQSRVGVTQQVTVRLQAIIPEPQNLQTIGGGMVGLMGGHVVGSAIGGVAGAVTLGPVGAVIGAQVGGFTGRVMGAQLGETLAGGHPLETVDARMQRLASDYMGSSMGAVVGTVVGQVTFGPVGAWVGNFVGDAVGGQLGTNAYQQIKQGQAQADRLWAIDHSGETTSELIGGTVGRLVAGAQGQMYGARLGTYFGRKISWQNAAGVPDPLATAAPISPPTDLPLDPELCYPKEEAPQN

**Supplementary Data 2: List of NCBI ftp server URL for downloading the 599 public proteomes of Cyanobacteria used in this study. These proteomes were published online before December 1st, 2017.**

'Nostoc_azollae'_0708,ftp://ftp.ncbi.nlm.nih.gov/genomes/all/GCF/000/196/515/GCF_000196515.1_ASM19651v1/GCF_000196515.1_ASM19651v1_translated_cds.faa

Acaryochloris_marina_MBIC11017,ftp://ftp.ncbi.nlm.nih.gov/genomes/all/GCF/000/018/105/GCF_000018105.1_ASM1810v1/GCF_000018105.1_ASM1810v1_translated_cds.faa

Acaryochloris_sp._CCMEE_5410,ftp://ftp.ncbi.nlm.nih.gov/genomes/all/GCF/000/238/775/GCF_000238775.1_ASM23877v2/GCF_000238775.1_ASM23877v2_translated_cds.faa

Aliterella_atlantica_CENA595,ftp://ftp.ncbi.nlm.nih.gov/genomes/all/GCF/000/952/155/GCF_000952155.1_asmbly001/GCF_000952155.1_asmbly001_translated_cds.faa

Alkalinema_sp._CACIAM_70d,ftp://ftp.ncbi.nlm.nih.gov/genomes/all/GCA/002/148/405/GCA_002148405.1_ASM214840v1/GCA_002148405.1_ASM214840v1_translated_cds.faa

Anabaena_cylindrica_PCC_7122,ftp://ftp.ncbi.nlm.nih.gov/genomes/all/GCF/002/367/955/GCF_002367955.1_ASM236795v1/GCF_002367955.1_ASM236795v1_translated_cds.faa

Anabaena_sp._4-3,ftp://ftp.ncbi.nlm.nih.gov/genomes/all/GCF/001/597/745/GCF_001597745.1_ASM159774v1/GCF_001597745.1_ASM159774v1_translated_cds.faa

Anabaena_sp._90,ftp://ftp.ncbi.nlm.nih.gov/genomes/all/GCF/000/312/705/GCF_000312705.1_ASM31270v1/GCF_000312705.1_ASM31270v1_translated_cds.faa

Anabaena_sp._AL09,ftp://ftp.ncbi.nlm.nih.gov/genomes/all/GCA/001/672/255/GCA_001672255.1_ASM167225v1/GCA_001672255.1_ASM167225v1_translated_cds.faa

Anabaena_sp._AL93,ftp://ftp.ncbi.nlm.nih.gov/genomes/all/GCA/001/672/085/GCA_001672085.1_ASM167208v1/GCA_001672085.1_ASM167208v1_translated_cds.faa

Anabaena_sp._CA_=_ATCC_33047,ftp://ftp.ncbi.nlm.nih.gov/genomes/all/GCF/001/597/855/GCF_001597855.1_ASM159785v1/GCF_001597855.1_ASM159785v1_translated_cds.faa

Anabaena_sp._CRKS33,ftp://ftp.ncbi.nlm.nih.gov/genomes/all/GCA/001/672/075/GCA_001672075.1_ASM167207v1/GCA_001672075.1_ASM167207v1_translated_cds.faa

Anabaena_sp._LE011-02,ftp://ftp.ncbi.nlm.nih.gov/genomes/all/GCA/001/672/225/GCA_001672225.1_ASM167222v1/GCA_001672225.1_ASM167222v1_translated_cds.faa

Anabaena_sp._MDT14b,ftp://ftp.ncbi.nlm.nih.gov/genomes/all/GCA/001/672/195/GCA_001672195.1_ASM167219v1/GCA_001672195.1_ASM167219v1_translated_cds.faa

Anabaena_sp._PCC_7108,ftp://ftp.ncbi.nlm.nih.gov/genomes/all/GCF/000/332/135/GCF_000332135.1_ASM33213v1/GCF_000332135.1_ASM33213v1_translated_cds.faa

Anabaena_sp._WA102,ftp://ftp.ncbi.nlm.nih.gov/genomes/all/GCF/001/277/295/GCF_001277295.1_ASM127729v1/GCF_001277295.1_ASM127729v1_translated_cds.faa

Anabaena_sp._WA113,ftp://ftp.ncbi.nlm.nih.gov/genomes/all/GCA/001/672/155/GCA_001672155.1_ASM167215v1/GCA_001672155.1_ASM167215v1_translated_cds.faa

Anabaenopsis_circularis_NIES-21,ftp://ftp.ncbi.nlm.nih.gov/genomes/all/GCF/002/367/975/GCF_002367975.1_ASM236797v1/GCF_002367975.1_ASM236797v1_translated_cds.faa

Aphanizomenon_flos-aquae_2012/KM1/D3,ftp://ftp.ncbi.nlm.nih.gov/genomes/all/GCF/000/789/435/GCF_000789435.1_ASM78943v1/GCF_000789435.1_ASM78943v1_translated_cds.faa

Aphanizomenon_flos-aquae_LD13,ftp://ftp.ncbi.nlm.nih.gov/genomes/all/GCA/001/672/165/GCA_001672165.1_ASM167216v1/GCA_001672165.1_ASM167216v1_translated_cds.faa

Aphanizomenon_flos-aquae_MDT14a,ftp://ftp.ncbi.nlm.nih.gov/genomes/all/GCA/001/672/095/GCA_001672095.1_ASM167209v1/GCA_001672095.1_ASM167209v1_translated_cds.faa

Aphanizomenon_flos-aquae_NIES-81,ftp://ftp.ncbi.nlm.nih.gov/genomes/all/GCF/000/521/175/GCF_000521175.1_Aflos454contigs199p/GCF_000521175.1_Aflos454contigs199p_translated_cds.faa

Aphanizomenon_flos-aquae_WA102,ftp://ftp.ncbi.nlm.nih.gov/genomes/all/GCA/001/672/105/GCA_001672105.1_ASM167210v1/GCA_001672105.1_ASM167210v1_translated_cds.faa

Aphanocapsa_montana_BDHKU210001,ftp://ftp.ncbi.nlm.nih.gov/genomes/all/GCF/000/817/745/GCF_000817745.1_ASM81774v1/GCF_000817745.1_ASM81774v1_protein.faa

Arthrospira_maxima_CS-328,ftp://ftp.ncbi.nlm.nih.gov/genomes/all/GCF/000/173/555/GCF_000173555.1_ASM17355v1/GCF_000173555.1_ASM17355v1_translated_cds.faa

Arthrospira_platensis_C1,ftp://ftp.ncbi.nlm.nih.gov/genomes/all/GCF/000/307/915/GCF_000307915.1_ASM30791v1/GCF_000307915.1_ASM30791v1_translated_cds.faa

Arthrospira_platensis_NIES-39,ftp://ftp.ncbi.nlm.nih.gov/genomes/all/GCF/000/210/375/GCF_000210375.1_ASM21037v1/GCF_000210375.1_ASM21037v1_translated_cds.faa

Arthrospira_platensis_YZ,ftp://ftp.ncbi.nlm.nih.gov/genomes/all/GCF/001/611/905/GCF_001611905.1_ASM161190v1/GCF_001611905.1_ASM161190v1_translated_cds.faa

Arthrospira_platensis_str._Paraca,ftp://ftp.ncbi.nlm.nih.gov/genomes/all/GCF/000/175/415/GCF_000175415.3_ASM17541v3/GCF_000175415.3_ASM17541v3_translated_cds.faa

Arthrospira_sp._PCC_8005,ftp://ftp.ncbi.nlm.nih.gov/genomes/all/GCF/000/176/895/GCF_000176895.2_ASM17689v2/GCF_000176895.2_ASM17689v2_translated_cds.faa

Arthrospira_sp._TJSD091,ftp://ftp.ncbi.nlm.nih.gov/genomes/all/GCF/000/974/245/GCF_000974245.1_ASM97424v1/GCF_000974245.1_ASM97424v1_translated_cds.faa

Aulosira_laxa_NIES-50,ftp://ftp.ncbi.nlm.nih.gov/genomes/all/GCF/002/368/055/GCF_002368055.1_ASM236805v1/GCF_002368055.1_ASM236805v1_translated_cds.faa

Calothrix_brevissima_NIES-22,ftp://ftp.ncbi.nlm.nih.gov/genomes/all/GCF/002/367/995/GCF_002367995.1_ASM236799v1/GCF_002367995.1_ASM236799v1_translated_cds.faa

Calothrix_elsteri_CCALA_953,ftp://ftp.ncbi.nlm.nih.gov/genomes/all/GCF/002/289/455/GCF_002289455.1_ASM228945v1/GCF_002289455.1_ASM228945v1_translated_cds.faa

Calothrix_parasitica_NIES-267,ftp://ftp.ncbi.nlm.nih.gov/genomes/all/GCF/002/368/095/GCF_002368095.1_ASM236809v1/GCF_002368095.1_ASM236809v1_translated_cds.faa

Calothrix_rhizosoleniae_SC01,ftp://ftp.ncbi.nlm.nih.gov/genomes/all/GCF/900/185/595/GCF_900185595.1_CalSC01_2013/GCF_900185595.1_CalSC01_2013_translated_cds.faa

Calothrix_sp._336/3,ftp://ftp.ncbi.nlm.nih.gov/genomes/all/GCF/000/734/895/GCF_000734895.2_ASM73489v2/GCF_000734895.2_ASM73489v2_translated_cds.faa

Calothrix_sp._HK-06,ftp://ftp.ncbi.nlm.nih.gov/genomes/all/GCF/001/904/745/GCF_001904745.1_ASM190474v1/GCF_001904745.1_ASM190474v1_translated_cds.faa

Calothrix_sp._NIES-2098,ftp://ftp.ncbi.nlm.nih.gov/genomes/all/GCF/002/368/175/GCF_002368175.1_ASM236817v1/GCF_002368175.1_ASM236817v1_translated_cds.faa

Calothrix_sp._NIES-2100,ftp://ftp.ncbi.nlm.nih.gov/genomes/all/GCF/002/368/195/GCF_002368195.1_ASM236819v1/GCF_002368195.1_ASM236819v1_translated_cds.faa

Calothrix_sp._NIES-3974,ftp://ftp.ncbi.nlm.nih.gov/genomes/all/GCF/002/368/395/GCF_002368395.1_ASM236839v1/GCF_002368395.1_ASM236839v1_translated_cds.faa

Calothrix_sp._NIES-4071,ftp://ftp.ncbi.nlm.nih.gov/genomes/all/GCF/002/368/455/GCF_002368455.1_ASM236845v1/GCF_002368455.1_ASM236845v1_translated_cds.faa

Calothrix_sp._NIES-4101,ftp://ftp.ncbi.nlm.nih.gov/genomes/all/GCF/002/368/375/GCF_002368375.1_ASM236837v1/GCF_002368375.1_ASM236837v1_translated_cds.faa

Calothrix_sp._NIES-4105,ftp://ftp.ncbi.nlm.nih.gov/genomes/all/GCF/002/368/415/GCF_002368415.1_ASM236841v1/GCF_002368415.1_ASM236841v1_translated_cds.faa

Calothrix_sp._PCC_6303,ftp://ftp.ncbi.nlm.nih.gov/genomes/all/GCF/000/317/435/GCF_000317435.1_ASM31743v1/GCF_000317435.1_ASM31743v1_translated_cds.faa

Calothrix_sp._PCC_7103,ftp://ftp.ncbi.nlm.nih.gov/genomes/all/GCF/000/331/305/GCF_000331305.1_ASM33130v1/GCF_000331305.1_ASM33130v1_translated_cds.faa

Calothrix_sp._PCC_7507,ftp://ftp.ncbi.nlm.nih.gov/genomes/all/GCF/000/316/575/GCF_000316575.1_ASM31657v1/GCF_000316575.1_ASM31657v1_translated_cds.faa

Candidatus_Atelocyanobacterium_thalassa_isolate_ALOHA,ftp://ftp.ncbi.nlm.nih.gov/genomes/all/GCF/000/025/125/GCF_000025125.1_ASM2512v1/GCF_000025125.1_ASM2512v1_translated_cds.faa

Candidatus_Atelocyanobacterium_thalassa_isolate_SIO64986,ftp://ftp.ncbi.nlm.nih.gov/genomes/all/GCF/000/737/945/GCF_000737945.1_ASM73794v1/GCF_000737945.1_ASM73794v1_protein.faa

Candidatus_Synechococcus_spongiarum,ftp://ftp.ncbi.nlm.nih.gov/genomes/all/GCF/900/047/545/GCF_900047545.1_m9/GCF_900047545.1_m9_translated_cds.faa

Candidatus_Synechococcus_spongiarum_142,ftp://ftp.ncbi.nlm.nih.gov/genomes/all/GCA/001/007/625/GCA_001007625.1_ASM100762v1/GCA_001007625.1_ASM100762v1_translated_cds.faa

Candidatus_Synechococcus_spongiarum_15L,ftp://ftp.ncbi.nlm.nih.gov/genomes/all/GCF/001/007/635/GCF_001007635.1_ASM100763v1/GCF_001007635.1_ASM100763v1_protein.faa

Candidatus_Synechococcus_spongiarum_LMB_bulk10D,ftp://ftp.ncbi.nlm.nih.gov/genomes/all/GCA/002/017/915/GCA_002017915.1_ASM201791v1/GCA_002017915.1_ASM201791v1_translated_cds.faa

Candidatus_Synechococcus_spongiarum_LMB_bulk10E,ftp://ftp.ncbi.nlm.nih.gov/genomes/all/GCA/002/017/955/GCA_002017955.1_ASM201795v1/GCA_002017955.1_ASM201795v1_translated_cds.faa

Candidatus_Synechococcus_spongiarum_LMB_bulk15M,ftp://ftp.ncbi.nlm.nih.gov/genomes/all/GCA/002/018/045/GCA_002018045.1_ASM201804v1/GCA_002018045.1_ASM201804v1_translated_cds.faa

Candidatus_Synechococcus_spongiarum_LMB_bulk15N,ftp://ftp.ncbi.nlm.nih.gov/genomes/all/GCF/002/018/015/GCF_002018015.1_ASM201801v1/GCF_002018015.1_ASM201801v1_translated_cds.faa

Candidatus_Synechococcus_spongiarum_SH4,ftp://ftp.ncbi.nlm.nih.gov/genomes/all/GCF/000/586/015/GCF_000586015.1_SynSpo.0/GCF_000586015.1_SynSpo.0_translated_cds.faa

Candidatus_Synechococcus_spongiarum_SP3,ftp://ftp.ncbi.nlm.nih.gov/genomes/all/GCF/001/007/665/GCF_001007665.1_ASM100766v1/GCF_001007665.1_ASM100766v1_protein.faa

Chamaesiphon_minutus_PCC_6605,ftp://ftp.ncbi.nlm.nih.gov/genomes/all/GCF/000/317/145/GCF_000317145.1_ASM31714v1/GCF_000317145.1_ASM31714v1_translated_cds.faa

Chlorogloeopsis_fritschii_PCC_6912,ftp://ftp.ncbi.nlm.nih.gov/genomes/all/GCF/000/317/285/GCF_000317285.1_ChlPCC6912_1.0/GCF_000317285.1_ChlPCC6912_1.0_translated_cds.faa

Chlorogloeopsis_fritschii_PCC_9212,ftp://ftp.ncbi.nlm.nih.gov/genomes/all/GCF/000/317/265/GCF_000317265.1_ChlPCC9212_1.0/GCF_000317265.1_ChlPCC9212_1.0_translated_cds.faa

Chondrocystis_sp._NIES-4102,ftp://ftp.ncbi.nlm.nih.gov/genomes/all/GCF/002/368/355/GCF_002368355.1_ASM236835v1/GCF_002368355.1_ASM236835v1_translated_cds.faa

Chroococcales_cyanobacterium_IPPAS_B-1203,ftp://ftp.ncbi.nlm.nih.gov/genomes/all/GCF/002/749/975/GCF_002749975.1_ASM274997v1/GCF_002749975.1_ASM274997v1_translated_cds.faa

Chroococcidiopsis_thermalis_PCC_7203,ftp://ftp.ncbi.nlm.nih.gov/genomes/all/GCF/000/317/125/GCF_000317125.1_ASM31712v1/GCF_000317125.1_ASM31712v1_translated_cds.faa

Chroogloeocystis_siderophila_5.2_s.c.1,ftp://ftp.ncbi.nlm.nih.gov/genomes/all/GCF/001/904/655/GCF_001904655.1_ASM190465v1/GCF_001904655.1_ASM190465v1_translated_cds.faa

Chrysosporum_ovalisporum,ftp://ftp.ncbi.nlm.nih.gov/genomes/all/GCA/001/458/455/GCA_001458455.1_Assembly_of_Aphanizomenon_ovalisporum/GCA_001458455.1_Assembly_of_Aphanizomenon_ovalisporum_translated_cds.faa

Coleofasciculus_chthonoplastes_PCC_7420,ftp://ftp.ncbi.nlm.nih.gov/genomes/all/GCF/000/155/555/GCF_000155555.1_ASM15555v1/GCF_000155555.1_ASM15555v1_translated_cds.faa

Crinalium_epipsammum_PCC_9333,ftp://ftp.ncbi.nlm.nih.gov/genomes/all/GCF/000/317/495/GCF_000317495.1_ASM31749v1/GCF_000317495.1_ASM31749v1_translated_cds.faa

Crocosphaera_watsonii_WH_0003,ftp://ftp.ncbi.nlm.nih.gov/genomes/all/GCF/000/235/665/GCF_000235665.1_ASM23566v2/GCF_000235665.1_ASM23566v2_translated_cds.faa

Crocosphaera_watsonii_WH_0005,ftp://ftp.ncbi.nlm.nih.gov/genomes/all/GCF/001/050/835/GCF_001050835.1_ASM105083v1/GCF_001050835.1_ASM105083v1_translated_cds.faa

Crocosphaera_watsonii_WH_0401,ftp://ftp.ncbi.nlm.nih.gov/genomes/all/GCF/001/039/615/GCF_001039615.1_WH0401_v1/GCF_001039615.1_WH0401_v1_translated_cds.faa

Crocosphaera_watsonii_WH_0402,ftp://ftp.ncbi.nlm.nih.gov/genomes/all/GCF/001/039/635/GCF_001039635.1_WH0402_v1/GCF_001039635.1_WH0402_v1_translated_cds.faa

Crocosphaera_watsonii_WH_8501,ftp://ftp.ncbi.nlm.nih.gov/genomes/all/GCF/000/167/195/GCF_000167195.1_ASM16719v1/GCF_000167195.1_ASM16719v1_translated_cds.faa

Crocosphaera_watsonii_WH_8502,ftp://ftp.ncbi.nlm.nih.gov/genomes/all/GCF/001/039/555/GCF_001039555.1_WH8502_v1/GCF_001039555.1_WH8502_v1_translated_cds.faa

Cyanobacteria_bacterium_13_1_20CM_4_61_6,ftp://ftp.ncbi.nlm.nih.gov/genomes/all/GCA/001/919/945/GCA_001919945.1_ASM191994v1/GCA_001919945.1_ASM191994v1_translated_cds.faa

Cyanobacteria_bacterium_13_1_40CM_2_61_4,ftp://ftp.ncbi.nlm.nih.gov/genomes/all/GCA/001/919/235/GCA_001919235.1_ASM191923v1/GCA_001919235.1_ASM191923v1_translated_cds.faa

Cyanobacteria_bacterium_TMED177,ftp://ftp.ncbi.nlm.nih.gov/genomes/all/GCA/002/170/825/GCA_002170825.1_ASM217082v1/GCA_002170825.1_ASM217082v1_translated_cds.faa

Cyanobacteria_bacterium_TMED188,ftp://ftp.ncbi.nlm.nih.gov/genomes/all/GCA/002/172/015/GCA_002172015.1_ASM217201v1/GCA_002172015.1_ASM217201v1_translated_cds.faa

Cyanobacteria_bacterium_TMED229,ftp://ftp.ncbi.nlm.nih.gov/genomes/all/GCA/002/169/615/GCA_002169615.1_ASM216961v1/GCA_002169615.1_ASM216961v1_translated_cds.faa

Cyanobacterium_aponinum_IPPAS_B-1201,ftp://ftp.ncbi.nlm.nih.gov/genomes/all/GCF/002/736/005/GCF_002736005.1_ASM273600v1/GCF_002736005.1_ASM273600v1_translated_cds.faa

Cyanobacterium_aponinum_PCC_10605,ftp://ftp.ncbi.nlm.nih.gov/genomes/all/GCF/000/317/675/GCF_000317675.1_ASM31767v1/GCF_000317675.1_ASM31767v1_translated_cds.faa

Cyanobacterium_sp._IPPAS_B-1200,ftp://ftp.ncbi.nlm.nih.gov/genomes/all/GCF/001/747/005/GCF_001747005.1_ASM174700v1/GCF_001747005.1_ASM174700v1_translated_cds.faa

Cyanobacterium_stanieri_PCC_7202,ftp://ftp.ncbi.nlm.nih.gov/genomes/all/GCF/000/317/655/GCF_000317655.1_ASM31765v1/GCF_000317655.1_ASM31765v1_protein.faa

Cyanobium_gracile_PCC_6307,ftp://ftp.ncbi.nlm.nih.gov/genomes/all/GCF/000/316/515/GCF_000316515.1_ASM31651v1/GCF_000316515.1_ASM31651v1_translated_cds.faa

Cyanobium_sp._ARS6,ftp://ftp.ncbi.nlm.nih.gov/genomes/all/GCA/002/687/115/GCA_002687115.1_ASM268711v1/GCA_002687115.1_ASM268711v1_translated_cds.faa

Cyanobium_sp._Baikal-G2,ftp://ftp.ncbi.nlm.nih.gov/genomes/all/GCA/002/737/505/GCA_002737505.1_ASM273750v1/GCA_002737505.1_ASM273750v1_translated_cds.faa

Cyanobium_sp._CACIAM_14,ftp://ftp.ncbi.nlm.nih.gov/genomes/all/GCF/000/708/525/GCF_000708525.1_ASM70852v1/GCF_000708525.1_ASM70852v1_protein.faa

Cyanobium_sp._MED195,ftp://ftp.ncbi.nlm.nih.gov/genomes/all/GCA/002/691/945/GCA_002691945.1_ASM269194v1/GCA_002691945.1_ASM269194v1_translated_cds.faa

Cyanobium_sp._MED843,ftp://ftp.ncbi.nlm.nih.gov/genomes/all/GCA/002/700/895/GCA_002700895.1_ASM270089v1/GCA_002700895.1_ASM270089v1_translated_cds.faa

Cyanobium_sp._NAT70,ftp://ftp.ncbi.nlm.nih.gov/genomes/all/GCA/002/701/375/GCA_002701375.1_ASM270137v1/GCA_002701375.1_ASM270137v1_translated_cds.faa

Cyanobium_sp._NIES-981,ftp://ftp.ncbi.nlm.nih.gov/genomes/all/GCF/900/088/535/GCF_900088535.1_ASM90008853v1/GCF_900088535.1_ASM90008853v1_translated_cds.faa

Cyanobium_sp._PCC_7001,ftp://ftp.ncbi.nlm.nih.gov/genomes/all/GCF/000/155/635/GCF_000155635.1_ASM15563v1/GCF_000155635.1_ASM15563v1_translated_cds.faa

Cyanobium_sp._RS427,ftp://ftp.ncbi.nlm.nih.gov/genomes/all/GCA/002/728/955/GCA_002728955.1_ASM272895v1/GCA_002728955.1_ASM272895v1_translated_cds.faa

Cyanobium_sp._SAT1300,ftp://ftp.ncbi.nlm.nih.gov/genomes/all/GCA/002/714/405/GCA_002714405.1_ASM271440v1/GCA_002714405.1_ASM271440v1_translated_cds.faa

Cyanothece_sp._ATCC_51142,ftp://ftp.ncbi.nlm.nih.gov/genomes/all/GCF/000/017/845/GCF_000017845.1_ASM1784v1/GCF_000017845.1_ASM1784v1_translated_cds.faa

Cyanothece_sp._ATCC_51472,ftp://ftp.ncbi.nlm.nih.gov/genomes/all/GCF/000/231/425/GCF_000231425.2_ASM23142v3/GCF_000231425.2_ASM23142v3_translated_cds.faa

Cyanothece_sp._CCY0110,ftp://ftp.ncbi.nlm.nih.gov/genomes/all/GCF/000/169/335/GCF_000169335.1_ASM16933v1/GCF_000169335.1_ASM16933v1_translated_cds.faa

Cyanothece_sp._PCC_7424,ftp://ftp.ncbi.nlm.nih.gov/genomes/all/GCF/000/021/825/GCF_000021825.1_ASM2182v1/GCF_000021825.1_ASM2182v1_translated_cds.faa

Cyanothece_sp._PCC_7425,ftp://ftp.ncbi.nlm.nih.gov/genomes/all/GCF/000/022/045/GCF_000022045.1_ASM2204v1/GCF_000022045.1_ASM2204v1_translated_cds.faa

Cyanothece_sp._PCC_7822,ftp://ftp.ncbi.nlm.nih.gov/genomes/all/GCF/000/147/335/GCF_000147335.1_ASM14733v1/GCF_000147335.1_ASM14733v1_translated_cds.faa

Cyanothece_sp._PCC_8801,ftp://ftp.ncbi.nlm.nih.gov/genomes/all/GCF/000/021/805/GCF_000021805.1_ASM2180v1/GCF_000021805.1_ASM2180v1_translated_cds.faa

Cyanothece_sp._PCC_8802,ftp://ftp.ncbi.nlm.nih.gov/genomes/all/GCF/000/024/045/GCF_000024045.1_ASM2404v1/GCF_000024045.1_ASM2404v1_translated_cds.faa

Cylindrospermopsis_raciborskii_CENA302,ftp://ftp.ncbi.nlm.nih.gov/genomes/all/GCF/002/027/345/GCF_002027345.1_ASM202734v1/GCF_002027345.1_ASM202734v1_translated_cds.faa

Cylindrospermopsis_raciborskii_CENA303,ftp://ftp.ncbi.nlm.nih.gov/genomes/all/GCF/002/114/155/GCF_002114155.1_ASM211415v1/GCF_002114155.1_ASM211415v1_translated_cds.faa

Cylindrospermopsis_raciborskii_CS-505,ftp://ftp.ncbi.nlm.nih.gov/genomes/all/GCF/000/175/835/GCF_000175835.1_ASM17583v1/GCF_000175835.1_ASM17583v1_translated_cds.faa

Cylindrospermopsis_raciborskii_CS-508,ftp://ftp.ncbi.nlm.nih.gov/genomes/all/GCF/001/858/115/GCF_001858115.1_ASM185811v1/GCF_001858115.1_ASM185811v1_translated_cds.faa

Cylindrospermopsis_raciborskii_CYLP,ftp://ftp.ncbi.nlm.nih.gov/genomes/all/GCF/002/321/935/GCF_002321935.1_ASM232193v1/GCF_002321935.1_ASM232193v1_translated_cds.faa

Cylindrospermopsis_raciborskii_CYRF,ftp://ftp.ncbi.nlm.nih.gov/genomes/all/GCF/002/321/945/GCF_002321945.1_ASM232194v1/GCF_002321945.1_ASM232194v1_translated_cds.faa

Cylindrospermopsis_raciborskii_ITEP-A1,ftp://ftp.ncbi.nlm.nih.gov/genomes/all/GCF/001/586/755/GCF_001586755.1_ASM158675v1/GCF_001586755.1_ASM158675v1_translated_cds.faa

Cylindrospermopsis_raciborskii_MVCC14,ftp://ftp.ncbi.nlm.nih.gov/genomes/all/GCF/001/858/125/GCF_001858125.1_ASM185812v1/GCF_001858125.1_ASM185812v1_translated_cds.faa

Cylindrospermopsis_sp._CR12,ftp://ftp.ncbi.nlm.nih.gov/genomes/all/GCF/001/432/185/GCF_001432185.1_ASM143218v1/GCF_001432185.1_ASM143218v1_translated_cds.faa

Cylindrospermum_stagnale_PCC_7417,ftp://ftp.ncbi.nlm.nih.gov/genomes/all/GCF/000/317/535/GCF_000317535.1_ASM31753v1/GCF_000317535.1_ASM31753v1_translated_cds.faa

Dactylococcopsis_salina_PCC_8305,ftp://ftp.ncbi.nlm.nih.gov/genomes/all/GCF/000/317/615/GCF_000317615.1_ASM31761v1/GCF_000317615.1_ASM31761v1_translated_cds.faa

Desertifilum_sp._IPPAS_B-1220,ftp://ftp.ncbi.nlm.nih.gov/genomes/all/GCF/001/746/915/GCF_001746915.1_ASM174691v1/GCF_001746915.1_ASM174691v1_translated_cds.faa

Dolichospermum_circinale_AWQC131C,ftp://ftp.ncbi.nlm.nih.gov/genomes/all/GCF/000/426/905/GCF_000426905.1_Dcir131C/GCF_000426905.1_Dcir131C_translated_cds.faa

Dolichospermum_circinale_AWQC310F,ftp://ftp.ncbi.nlm.nih.gov/genomes/all/GCF/000/426/925/GCF_000426925.1_Acir310F/GCF_000426925.1_Acir310F_translated_cds.faa

Dolichospermum_compactum_NIES-806,ftp://ftp.ncbi.nlm.nih.gov/genomes/all/GCF/002/368/115/GCF_002368115.1_ASM236811v1/GCF_002368115.1_ASM236811v1_translated_cds.faa

Fischerella_major_NIES-592,ftp://ftp.ncbi.nlm.nih.gov/genomes/all/GCF/001/904/645/GCF_001904645.1_ASM190464v1/GCF_001904645.1_ASM190464v1_translated_cds.faa

Fischerella_muscicola_PCC_7414,ftp://ftp.ncbi.nlm.nih.gov/genomes/all/GCF/000/317/205/GCF_000317205.1_FisPCC7414_1.0/GCF_000317205.1_FisPCC7414_1.0_translated_cds.faa

Fischerella_muscicola_SAG_1427-1_=_PCC_73103,ftp://ftp.ncbi.nlm.nih.gov/genomes/all/GCF/000/317/245/GCF_000317245.1_FisPCC73103_1.0/GCF_000317245.1_FisPCC73103_1.0_translated_cds.faa

Fischerella_sp._NIES-3754,ftp://ftp.ncbi.nlm.nih.gov/genomes/all/GCF/001/548/455/GCF_001548455.1_ASM154845v1/GCF_001548455.1_ASM154845v1_translated_cds.faa

Fischerella_sp._NIES-4106,ftp://ftp.ncbi.nlm.nih.gov/genomes/all/GCF/002/368/315/GCF_002368315.1_ASM236831v1/GCF_002368315.1_ASM236831v1_translated_cds.faa

Fischerella_sp._PCC_9339,ftp://ftp.ncbi.nlm.nih.gov/genomes/all/GCF/000/315/585/GCF_000315585.1_ASM31558v1/GCF_000315585.1_ASM31558v1_translated_cds.faa

Fischerella_sp._PCC_9431,ftp://ftp.ncbi.nlm.nih.gov/genomes/all/GCF/000/447/295/GCF_000447295.1_ASM44729v1/GCF_000447295.1_ASM44729v1_translated_cds.faa

Fischerella_sp._PCC_9605,ftp://ftp.ncbi.nlm.nih.gov/genomes/all/GCF/000/517/105/GCF_000517105.1_ASM51710v1/GCF_000517105.1_ASM51710v1_translated_cds.faa

Fischerella_thermalis,ftp://ftp.ncbi.nlm.nih.gov/genomes/all/GCF/000/231/365/GCF_000231365.1_ASM23136v1/GCF_000231365.1_ASM23136v1_translated_cds.faa

Fischerella_thermalis_PCC_7521,ftp://ftp.ncbi.nlm.nih.gov/genomes/all/GCF/000/317/225/GCF_000317225.1_FisPCC7521_1.0/GCF_000317225.1_FisPCC7521_1.0_translated_cds.faa

Fortiea_contorta_PCC_7126,ftp://ftp.ncbi.nlm.nih.gov/genomes/all/GCF/000/332/295/GCF_000332295.1_ASM33229v1/GCF_000332295.1_ASM33229v1_translated_cds.faa

Fremyella_diplosiphon_NIES-3275,ftp://ftp.ncbi.nlm.nih.gov/genomes/all/GCF/002/368/275/GCF_002368275.1_ASM236827v1/GCF_002368275.1_ASM236827v1_translated_cds.faa

Geitlerinema_sp._FC_II,ftp://ftp.ncbi.nlm.nih.gov/genomes/all/GCA/002/286/845/GCA_002286845.1_ASM228684v1/GCA_002286845.1_ASM228684v1_translated_cds.faa

Geitlerinema_sp._PCC_7105,ftp://ftp.ncbi.nlm.nih.gov/genomes/all/GCF/000/332/355/GCF_000332355.1_ASM33235v1/GCF_000332355.1_ASM33235v1_translated_cds.faa

Geitlerinema_sp._PCC_7407,ftp://ftp.ncbi.nlm.nih.gov/genomes/all/GCF/000/317/045/GCF_000317045.1_ASM31704v1/GCF_000317045.1_ASM31704v1_translated_cds.faa

Geitlerinema_sp._PCC_9228,ftp://ftp.ncbi.nlm.nih.gov/genomes/all/GCF/001/870/905/GCF_001870905.1_ASM187090v1/GCF_001870905.1_ASM187090v1_translated_cds.faa

Geminocystis_herdmanii_PCC_6308,ftp://ftp.ncbi.nlm.nih.gov/genomes/all/GCF/000/332/235/GCF_000332235.1_ASM33223v1/GCF_000332235.1_ASM33223v1_translated_cds.faa

Geminocystis_sp._NIES-3708,ftp://ftp.ncbi.nlm.nih.gov/genomes/all/GCF/001/548/095/GCF_001548095.1_Gm3709_assembly_1.0/GCF_001548095.1_Gm3709_assembly_1.0_translated_cds.faa

Geminocystis_sp._NIES-3709,ftp://ftp.ncbi.nlm.nih.gov/genomes/all/GCF/001/548/115/GCF_001548115.1_Gm3709_assembly_1.0/GCF_001548115.1_Gm3709_assembly_1.0_translated_cds.faa

Gloeobacter_kilaueensis_JS1,ftp://ftp.ncbi.nlm.nih.gov/genomes/all/GCF/000/484/535/GCF_000484535.1_ASM48453v1/GCF_000484535.1_ASM48453v1_translated_cds.faa

Gloeobacter_violaceus_PCC_7421,ftp://ftp.ncbi.nlm.nih.gov/genomes/all/GCF/000/011/385/GCF_000011385.1_ASM1138v1/GCF_000011385.1_ASM1138v1_translated_cds.faa

Gloeocapsa_sp._PCC_73106,ftp://ftp.ncbi.nlm.nih.gov/genomes/all/GCF/000/332/035/GCF_000332035.1_ASM33203v1/GCF_000332035.1_ASM33203v1_translated_cds.faa

Gloeocapsa_sp._PCC_7428,ftp://ftp.ncbi.nlm.nih.gov/genomes/all/GCF/000/317/555/GCF_000317555.1_ASM31755v1/GCF_000317555.1_ASM31755v1_translated_cds.faa

Gloeomargarita_lithophora_Alchichica-D10,ftp://ftp.ncbi.nlm.nih.gov/genomes/all/GCF/001/870/225/GCF_001870225.1_ASM187022v1/GCF_001870225.1_ASM187022v1_translated_cds.faa

Halomicronema_hongdechloris_C2206,ftp://ftp.ncbi.nlm.nih.gov/genomes/all/GCF/002/075/285/GCF_002075285.2_ASM207528v2/GCF_002075285.2_ASM207528v2_translated_cds.faa

Halothece_sp._PCC_7418,ftp://ftp.ncbi.nlm.nih.gov/genomes/all/GCF/000/317/635/GCF_000317635.1_ASM31763v1/GCF_000317635.1_ASM31763v1_translated_cds.faa

Hapalosiphon_sp._MRB220,ftp://ftp.ncbi.nlm.nih.gov/genomes/all/GCF/001/275/395/GCF_001275395.1_ASM127539v1/GCF_001275395.1_ASM127539v1_translated_cds.faa

Hassallia_byssoidea_VB512170,ftp://ftp.ncbi.nlm.nih.gov/genomes/all/GCF/000/817/785/GCF_000817785.1_V1.0/GCF_000817785.1_V1.0_protein.faa

Hydrococcus_rivularis_NIES-593,ftp://ftp.ncbi.nlm.nih.gov/genomes/all/GCF/001/904/635/GCF_001904635.1_ASM190463v1/GCF_001904635.1_ASM190463v1_translated_cds.faa

Hydrocoleum_sp._CS-953,ftp://ftp.ncbi.nlm.nih.gov/genomes/all/GCF/002/260/545/GCF_002260545.1_ASM226054v1/GCF_002260545.1_ASM226054v1_translated_cds.faa

Kamptonema_formosum_PCC_6407,ftp://ftp.ncbi.nlm.nih.gov/genomes/all/GCF/000/332/155/GCF_000332155.1_ASM33215v1/GCF_000332155.1_ASM33215v1_translated_cds.faa

Leptolyngbya_boryana_IAM_M-101,ftp://ftp.ncbi.nlm.nih.gov/genomes/all/GCF/002/142/475/GCF_002142475.1_ASM214247v1/GCF_002142475.1_ASM214247v1_translated_cds.faa

Leptolyngbya_boryana_NIES-2135,ftp://ftp.ncbi.nlm.nih.gov/genomes/all/GCF/002/368/255/GCF_002368255.1_ASM236825v1/GCF_002368255.1_ASM236825v1_translated_cds.faa

Leptolyngbya_boryana_PCC_6306,ftp://ftp.ncbi.nlm.nih.gov/genomes/all/GCF/000/353/285/GCF_000353285.1_ASM35328v1/GCF_000353285.1_ASM35328v1_translated_cds.faa

Leptolyngbya_boryana_dg5,ftp://ftp.ncbi.nlm.nih.gov/genomes/all/GCF/002/142/495/GCF_002142495.1_ASM214249v1/GCF_002142495.1_ASM214249v1_translated_cds.faa

Leptolyngbya_ohadii_IS1,ftp://ftp.ncbi.nlm.nih.gov/genomes/all/GCF/002/215/035/GCF_002215035.1_ASM221503v1/GCF_002215035.1_ASM221503v1_translated_cds.faa

Leptolyngbya_sp._'hensonii',ftp://ftp.ncbi.nlm.nih.gov/genomes/all/GCF/001/939/115/GCF_001939115.1_ASM193911v1/GCF_001939115.1_ASM193911v1_translated_cds.faa

Leptolyngbya_sp._BC1307,ftp://ftp.ncbi.nlm.nih.gov/genomes/all/GCF/002/286/735/GCF_002286735.1_ASM228673v1/GCF_002286735.1_ASM228673v1_translated_cds.faa

Leptolyngbya_sp._Heron_Island_J,ftp://ftp.ncbi.nlm.nih.gov/genomes/all/GCF/000/482/245/GCF_000482245.1_LepHIscaffolds/GCF_000482245.1_LepHIscaffolds_translated_cds.faa

Leptolyngbya_sp._KIOST-1,ftp://ftp.ncbi.nlm.nih.gov/genomes/all/GCF/000/763/385/GCF_000763385.1_ASM76338v1/GCF_000763385.1_ASM76338v1_translated_cds.faa

Leptolyngbya_sp._NIES-2104,ftp://ftp.ncbi.nlm.nih.gov/genomes/all/GCF/001/485/215/GCF_001485215.1_ASM148521v1/GCF_001485215.1_ASM148521v1_translated_cds.faa

Leptolyngbya_sp._NIES-3755,ftp://ftp.ncbi.nlm.nih.gov/genomes/all/GCF/001/548/435/GCF_001548435.1_ASM154843v1/GCF_001548435.1_ASM154843v1_translated_cds.faa

Leptolyngbya_sp._O-77,ftp://ftp.ncbi.nlm.nih.gov/genomes/all/GCF/001/548/395/GCF_001548395.1_ASM154839v1/GCF_001548395.1_ASM154839v1_translated_cds.faa

Leptolyngbya_sp._PCC_6406,ftp://ftp.ncbi.nlm.nih.gov/genomes/all/GCF/000/332/095/GCF_000332095.2_ASM33209v2/GCF_000332095.2_ASM33209v2_translated_cds.faa

Leptolyngbya_sp._PCC_7375,ftp://ftp.ncbi.nlm.nih.gov/genomes/all/GCF/000/316/115/GCF_000316115.1_ASM31611v1/GCF_000316115.1_ASM31611v1_translated_cds.faa

Leptolyngbya_sp._PCC_7376,ftp://ftp.ncbi.nlm.nih.gov/genomes/all/GCF/000/316/605/GCF_000316605.1_ASM31660v1/GCF_000316605.1_ASM31660v1_translated_cds.faa

Leptolyngbya_valderiana_BDU_20041,ftp://ftp.ncbi.nlm.nih.gov/genomes/all/GCF/001/637/395/GCF_001637395.1_ASM163739v1/GCF_001637395.1_ASM163739v1_translated_cds.faa

Limnoraphis_robusta_CS-951,ftp://ftp.ncbi.nlm.nih.gov/genomes/all/GCF/000/972/705/GCF_000972705.2_ASM97270v2/GCF_000972705.2_ASM97270v2_translated_cds.faa

Limnothrix_rosea_IAM_M-220,ftp://ftp.ncbi.nlm.nih.gov/genomes/all/GCF/001/904/615/GCF_001904615.1_ASM190461v1/GCF_001904615.1_ASM190461v1_translated_cds.faa

Limnothrix_sp._CACIAM_69d,ftp://ftp.ncbi.nlm.nih.gov/genomes/all/GCA/001/913/845/GCA_001913845.2_ASM191384v2/GCA_001913845.2_ASM191384v2_translated_cds.faa

Limnothrix_sp._P13C2,ftp://ftp.ncbi.nlm.nih.gov/genomes/all/GCA/001/698/445/GCA_001698445.1_ASM169844v1/GCA_001698445.1_ASM169844v1_translated_cds.faa

Limnothrix_sp._PR1529,ftp://ftp.ncbi.nlm.nih.gov/genomes/all/GCF/002/742/025/GCF_002742025.1_ASM274202v1/GCF_002742025.1_ASM274202v1_translated_cds.faa

Lyngbya_aestuarii_BL_J,ftp://ftp.ncbi.nlm.nih.gov/genomes/all/GCF/000/478/195/GCF_000478195.2_ASM47819v2/GCF_000478195.2_ASM47819v2_translated_cds.faa

Lyngbya_confervoides_BDU141951,ftp://ftp.ncbi.nlm.nih.gov/genomes/all/GCF/000/817/775/GCF_000817775.1_ASM81777v1/GCF_000817775.1_ASM81777v1_translated_cds.faa

Lyngbya_sp._PCC_8106,ftp://ftp.ncbi.nlm.nih.gov/genomes/all/GCF/000/169/095/GCF_000169095.1_ASM16909v1/GCF_000169095.1_ASM16909v1_translated_cds.faa

Mastigocladopsis_repens_PCC_10914,ftp://ftp.ncbi.nlm.nih.gov/genomes/all/GCF/000/315/565/GCF_000315565.1_ASM31556v1/GCF_000315565.1_ASM31556v1_translated_cds.faa

Mastigocladus_laminosus_UU774,ftp://ftp.ncbi.nlm.nih.gov/genomes/all/GCF/001/990/805/GCF_001990805.1_ASM199080v1/GCF_001990805.1_ASM199080v1_translated_cds.faa

Mastigocoleus_testarum_BC008,ftp://ftp.ncbi.nlm.nih.gov/genomes/all/GCF/000/472/885/GCF_000472885.1_ASM47288v1/GCF_000472885.1_ASM47288v1_protein.faa

Microcoleus_sp._PCC_7113,ftp://ftp.ncbi.nlm.nih.gov/genomes/all/GCF/000/317/515/GCF_000317515.1_ASM31751v1/GCF_000317515.1_ASM31751v1_translated_cds.faa

Microcoleus_vaginatus_FGP-2,ftp://ftp.ncbi.nlm.nih.gov/genomes/all/GCF/000/214/075/GCF_000214075.1_ASM21407v1/GCF_000214075.1_ASM21407v1_translated_cds.faa

Microcystis_aeruginosa_CACIAM_03,ftp://ftp.ncbi.nlm.nih.gov/genomes/all/GCA/001/706/385/GCA_001706385.1_ASM170638v1/GCA_001706385.1_ASM170638v1_translated_cds.faa

Microcystis_aeruginosa_CHAOHU_1326,ftp://ftp.ncbi.nlm.nih.gov/genomes/all/GCF/001/895/325/GCF_001895325.1_ASM189532v1/GCF_001895325.1_ASM189532v1_translated_cds.faa

Microcystis_aeruginosa_DIANCHI905,ftp://ftp.ncbi.nlm.nih.gov/genomes/all/GCF/000/332/585/GCF_000332585.1_MicAerD1.0/GCF_000332585.1_MicAerD1.0_translated_cds.faa

Microcystis_aeruginosa_KW,ftp://ftp.ncbi.nlm.nih.gov/genomes/all/GCF/002/025/445/GCF_002025445.1_KWv02/GCF_002025445.1_KWv02_translated_cds.faa

Microcystis_aeruginosa_NIES-2481,ftp://ftp.ncbi.nlm.nih.gov/genomes/all/GCF/001/704/955/GCF_001704955.2_ASM170495v2/GCF_001704955.2_ASM170495v2_translated_cds.faa

Microcystis_aeruginosa_NIES-2549,ftp://ftp.ncbi.nlm.nih.gov/genomes/all/GCF/000/981/785/GCF_000981785.2_ASM98178v2/GCF_000981785.2_ASM98178v2_translated_cds.faa

Microcystis_aeruginosa_NIES-44,ftp://ftp.ncbi.nlm.nih.gov/genomes/all/GCF/000/787/675/GCF_000787675.1_ASM78767v1/GCF_000787675.1_ASM78767v1_translated_cds.faa

Microcystis_aeruginosa_NIES-843,ftp://ftp.ncbi.nlm.nih.gov/genomes/all/GCF/000/010/625/GCF_000010625.1_ASM1062v1/GCF_000010625.1_ASM1062v1_translated_cds.faa

Microcystis_aeruginosa_NIES-88,ftp://ftp.ncbi.nlm.nih.gov/genomes/all/GCF/001/578/075/GCF_001578075.1_ASM157807v1/GCF_001578075.1_ASM157807v1_translated_cds.faa

Microcystis_aeruginosa_NIES-98,ftp://ftp.ncbi.nlm.nih.gov/genomes/all/GCF/001/725/075/GCF_001725075.1_ASM172507v1/GCF_001725075.1_ASM172507v1_translated_cds.faa

Microcystis_aeruginosa_NaRes975,ftp://ftp.ncbi.nlm.nih.gov/genomes/all/GCF/001/885/655/GCF_001885655.1_ASM188565v1/GCF_001885655.1_ASM188565v1_translated_cds.faa

Microcystis_aeruginosa_PCC_7005,ftp://ftp.ncbi.nlm.nih.gov/genomes/all/GCF/000/599/945/GCF_000599945.1_Mic70051.0/GCF_000599945.1_Mic70051.0_translated_cds.faa

Microcystis_aeruginosa_PCC_7806SL,ftp://ftp.ncbi.nlm.nih.gov/genomes/all/GCF/002/095/975/GCF_002095975.1_ASM209597v1/GCF_002095975.1_ASM209597v1_translated_cds.faa

Microcystis_aeruginosa_PCC_7941,ftp://ftp.ncbi.nlm.nih.gov/genomes/all/GCF/000/312/205/GCF_000312205.1_ASM31220v1/GCF_000312205.1_ASM31220v1_translated_cds.faa

Microcystis_aeruginosa_PCC_9432,ftp://ftp.ncbi.nlm.nih.gov/genomes/all/GCF/000/307/995/GCF_000307995.1_ASM30799v2/GCF_000307995.1_ASM30799v2_translated_cds.faa

Microcystis_aeruginosa_PCC_9443,ftp://ftp.ncbi.nlm.nih.gov/genomes/all/GCF/000/312/185/GCF_000312185.1_ASM31218v1/GCF_000312185.1_ASM31218v1_translated_cds.faa

Microcystis_aeruginosa_PCC_9701,ftp://ftp.ncbi.nlm.nih.gov/genomes/all/GCF/000/312/285/GCF_000312285.1_ASM31228v1/GCF_000312285.1_ASM31228v1_translated_cds.faa

Microcystis_aeruginosa_PCC_9717,ftp://ftp.ncbi.nlm.nih.gov/genomes/all/GCF/000/312/165/GCF_000312165.1_ASM31216v1/GCF_000312165.1_ASM31216v1_translated_cds.faa

Microcystis_aeruginosa_PCC_9806,ftp://ftp.ncbi.nlm.nih.gov/genomes/all/GCF/000/312/725/GCF_000312725.1_ASM31272v1/GCF_000312725.1_ASM31272v1_translated_cds.faa

Microcystis_aeruginosa_PCC_9807,ftp://ftp.ncbi.nlm.nih.gov/genomes/all/GCF/000/312/225/GCF_000312225.1_ASM31222v1/GCF_000312225.1_ASM31222v1_translated_cds.faa

Microcystis_aeruginosa_PCC_9808,ftp://ftp.ncbi.nlm.nih.gov/genomes/all/GCF/000/312/245/GCF_000312245.1_ASM31224v1/GCF_000312245.1_ASM31224v1_translated_cds.faa

Microcystis_aeruginosa_PCC_9809,ftp://ftp.ncbi.nlm.nih.gov/genomes/all/GCF/000/312/265/GCF_000312265.1_ASM31226v1/GCF_000312265.1_ASM31226v1_translated_cds.faa

Microcystis_aeruginosa_SPC777,ftp://ftp.ncbi.nlm.nih.gov/genomes/all/GCF/000/412/595/GCF_000412595.1_spc777-v1/GCF_000412595.1_spc777-v1_translated_cds.faa

Microcystis_aeruginosa_TAIHU98,ftp://ftp.ncbi.nlm.nih.gov/genomes/all/GCF/000/330/925/GCF_000330925.1_MicAerT1.0/GCF_000330925.1_MicAerT1.0_translated_cds.faa

Microcystis_panniformis_FACHB-1757,ftp://ftp.ncbi.nlm.nih.gov/genomes/all/GCF/001/264/245/GCF_001264245.1_ASM126424v1/GCF_001264245.1_ASM126424v1_translated_cds.faa

Microcystis_sp._T1-4,ftp://ftp.ncbi.nlm.nih.gov/genomes/all/GCF/000/297/435/GCF_000297435.1_ASM29743v1/GCF_000297435.1_ASM29743v1_translated_cds.faa

Moorea_bouillonii_PNG,ftp://ftp.ncbi.nlm.nih.gov/genomes/all/GCF/001/942/495/GCF_001942495.1_ASM194249v1/GCF_001942495.1_ASM194249v1_translated_cds.faa

Moorea_producens_3L,ftp://ftp.ncbi.nlm.nih.gov/genomes/all/GCF/000/211/815/GCF_000211815.1_ASM21181v1/GCF_000211815.1_ASM21181v1_translated_cds.faa

Moorea_producens_JHB,ftp://ftp.ncbi.nlm.nih.gov/genomes/all/GCF/001/854/205/GCF_001854205.1_ASM185420v1/GCF_001854205.1_ASM185420v1_translated_cds.faa

Moorea_producens_PAL,ftp://ftp.ncbi.nlm.nih.gov/genomes/all/GCF/001/942/475/GCF_001942475.1_ASM194247v1/GCF_001942475.1_ASM194247v1_translated_cds.faa

Moorea_producens_PAL-8-15-08-1,ftp://ftp.ncbi.nlm.nih.gov/genomes/all/GCF/001/767/235/GCF_001767235.1_ASM176723v1/GCF_001767235.1_ASM176723v1_translated_cds.faa

Myxosarcina_sp._GI1,ftp://ftp.ncbi.nlm.nih.gov/genomes/all/GCF/000/756/305/GCF_000756305.1_ASM75630v1/GCF_000756305.1_ASM75630v1_translated_cds.faa

Neosynechococcus_sphagnicola_sy1,ftp://ftp.ncbi.nlm.nih.gov/genomes/all/GCF/000/775/285/GCF_000775285.1_ASM77528v1/GCF_000775285.1_ASM77528v1_translated_cds.faa

Nodosilinea_nodulosa_PCC_7104,ftp://ftp.ncbi.nlm.nih.gov/genomes/all/GCF/000/309/385/GCF_000309385.1_ASM30938v1/GCF_000309385.1_ASM30938v1_translated_cds.faa

Nodularia_sp._NIES-3585,ftp://ftp.ncbi.nlm.nih.gov/genomes/all/GCF/002/218/065/GCF_002218065.1_ASM221806v1/GCF_002218065.1_ASM221806v1_translated_cds.faa

Nodularia_spumigena_CCY9414,ftp://ftp.ncbi.nlm.nih.gov/genomes/all/GCF/000/340/565/GCF_000340565.2_ASM34056v3/GCF_000340565.2_ASM34056v3_translated_cds.faa

Nodularia_spumigena_CENA596,ftp://ftp.ncbi.nlm.nih.gov/genomes/all/GCF/001/623/485/GCF_001623485.1_ASM162348v1/GCF_001623485.1_ASM162348v1_translated_cds.faa

Nostoc_calcicola_FACHB-389,ftp://ftp.ncbi.nlm.nih.gov/genomes/all/GCF/001/904/715/GCF_001904715.1_ASM190471v1/GCF_001904715.1_ASM190471v1_translated_cds.faa

Nostoc_carneum_NIES-2107,ftp://ftp.ncbi.nlm.nih.gov/genomes/all/GCF/002/368/155/GCF_002368155.1_ASM236815v1/GCF_002368155.1_ASM236815v1_translated_cds.faa

Nostoc_linckia_NIES-25,ftp://ftp.ncbi.nlm.nih.gov/genomes/all/GCF/002/368/035/GCF_002368035.1_ASM236803v1/GCF_002368035.1_ASM236803v1_translated_cds.faa

Nostoc_linckia_z1,ftp://ftp.ncbi.nlm.nih.gov/genomes/all/GCF/002/607/965/GCF_002607965.1_ASM260796v1/GCF_002607965.1_ASM260796v1_translated_cds.faa

Nostoc_linckia_z13,ftp://ftp.ncbi.nlm.nih.gov/genomes/all/GCF/002/608/275/GCF_002608275.1_ASM260827v1/GCF_002608275.1_ASM260827v1_translated_cds.faa

Nostoc_linckia_z14,ftp://ftp.ncbi.nlm.nih.gov/genomes/all/GCF/002/608/345/GCF_002608345.1_ASM260834v1/GCF_002608345.1_ASM260834v1_translated_cds.faa

Nostoc_linckia_z15,ftp://ftp.ncbi.nlm.nih.gov/genomes/all/GCF/002/608/385/GCF_002608385.1_ASM260838v1/GCF_002608385.1_ASM260838v1_translated_cds.faa

Nostoc_linckia_z16,ftp://ftp.ncbi.nlm.nih.gov/genomes/all/GCF/002/608/325/GCF_002608325.1_ASM260832v1/GCF_002608325.1_ASM260832v1_translated_cds.faa

Nostoc_linckia_z18,ftp://ftp.ncbi.nlm.nih.gov/genomes/all/GCF/002/607/925/GCF_002607925.1_ASM260792v1/GCF_002607925.1_ASM260792v1_translated_cds.faa

Nostoc_linckia_z2,ftp://ftp.ncbi.nlm.nih.gov/genomes/all/GCF/002/607/955/GCF_002607955.1_ASM260795v1/GCF_002607955.1_ASM260795v1_translated_cds.faa

Nostoc_linckia_z3,ftp://ftp.ncbi.nlm.nih.gov/genomes/all/GCF/002/608/015/GCF_002608015.1_ASM260801v1/GCF_002608015.1_ASM260801v1_translated_cds.faa

Nostoc_linckia_z4,ftp://ftp.ncbi.nlm.nih.gov/genomes/all/GCF/002/608/075/GCF_002608075.1_ASM260807v1/GCF_002608075.1_ASM260807v1_translated_cds.faa

Nostoc_linckia_z6,ftp://ftp.ncbi.nlm.nih.gov/genomes/all/GCF/002/608/105/GCF_002608105.1_ASM260810v1/GCF_002608105.1_ASM260810v1_translated_cds.faa

Nostoc_linckia_z7,ftp://ftp.ncbi.nlm.nih.gov/genomes/all/GCF/002/608/135/GCF_002608135.1_ASM260813v1/GCF_002608135.1_ASM260813v1_translated_cds.faa

Nostoc_linckia_z8,ftp://ftp.ncbi.nlm.nih.gov/genomes/all/GCF/002/608/145/GCF_002608145.1_ASM260814v1/GCF_002608145.1_ASM260814v1_translated_cds.faa

Nostoc_linckia_z9,ftp://ftp.ncbi.nlm.nih.gov/genomes/all/GCF/002/608/225/GCF_002608225.1_ASM260822v1/GCF_002608225.1_ASM260822v1_translated_cds.faa

Nostoc_piscinale_CENA21,ftp://ftp.ncbi.nlm.nih.gov/genomes/all/GCF/001/298/445/GCF_001298445.1_ASM129844v1/GCF_001298445.1_ASM129844v1_translated_cds.faa

Nostoc_punctiforme_PCC_73102,ftp://ftp.ncbi.nlm.nih.gov/genomes/all/GCF/000/020/025/GCF_000020025.1_ASM2002v1/GCF_000020025.1_ASM2002v1_translated_cds.faa

Nostoc_sp._'Peltigera_malacea_cyanobiont'_DB3992,ftp://ftp.ncbi.nlm.nih.gov/genomes/all/GCF/002/631/755/GCF_002631755.1_ASM263175v1/GCF_002631755.1_ASM263175v1_translated_cds.faa

Nostoc_sp._'Peltigera_membranacea_cyanobiont'_210A,ftp://ftp.ncbi.nlm.nih.gov/genomes/all/GCF/002/246/015/GCF_002246015.1_ASM224601v1/GCF_002246015.1_ASM224601v1_translated_cds.faa

Nostoc_sp._'Peltigera_membranacea_cyanobiont'_213,ftp://ftp.ncbi.nlm.nih.gov/genomes/all/GCF/002/245/975/GCF_002245975.1_ASM224597v1/GCF_002245975.1_ASM224597v1_translated_cds.faa

Nostoc_sp._'Peltigera_membranacea_cyanobiont'_232,ftp://ftp.ncbi.nlm.nih.gov/genomes/all/GCF/002/245/985/GCF_002245985.1_ASM224598v1/GCF_002245985.1_ASM224598v1_translated_cds.faa

Nostoc_sp._106C,ftp://ftp.ncbi.nlm.nih.gov/genomes/all/GCF/002/154/725/GCF_002154725.1_ASM215472v1/GCF_002154725.1_ASM215472v1_translated_cds.faa

Nostoc_sp._KVJ20,ftp://ftp.ncbi.nlm.nih.gov/genomes/all/GCF/001/712/795/GCF_001712795.1_ASM171279v1/GCF_001712795.1_ASM171279v1_translated_cds.faa

Nostoc_sp._MBR_210,ftp://ftp.ncbi.nlm.nih.gov/genomes/all/GCA/001/698/435/GCA_001698435.1_ASM169843v1/GCA_001698435.1_ASM169843v1_translated_cds.faa

Nostoc_sp._NIES-2111,ftp://ftp.ncbi.nlm.nih.gov/genomes/all/GCF/002/368/215/GCF_002368215.1_ASM236821v1/GCF_002368215.1_ASM236821v1_translated_cds.faa

Nostoc_sp._NIES-3756,ftp://ftp.ncbi.nlm.nih.gov/genomes/all/GCF/001/548/375/GCF_001548375.1_ASM154837v1/GCF_001548375.1_ASM154837v1_translated_cds.faa

Nostoc_sp._NIES-4103,ftp://ftp.ncbi.nlm.nih.gov/genomes/all/GCF/002/368/335/GCF_002368335.1_ASM236833v1/GCF_002368335.1_ASM236833v1_translated_cds.faa

Nostoc_sp._PCC_7107,ftp://ftp.ncbi.nlm.nih.gov/genomes/all/GCF/000/316/625/GCF_000316625.1_ASM31662v1/GCF_000316625.1_ASM31662v1_translated_cds.faa

Nostoc_sp._PCC_7120,ftp://ftp.ncbi.nlm.nih.gov/genomes/all/GCF/000/009/705/GCF_000009705.1_ASM970v1/GCF_000009705.1_ASM970v1_translated_cds.faa

Nostoc_sp._PCC_7524,ftp://ftp.ncbi.nlm.nih.gov/genomes/all/GCF/000/316/645/GCF_000316645.1_ASM31664v1/GCF_000316645.1_ASM31664v1_translated_cds.faa

Nostoc_sp._RF31Y,ftp://ftp.ncbi.nlm.nih.gov/genomes/all/GCF/002/155/185/GCF_002155185.1_ASM215518v1/GCF_002155185.1_ASM215518v1_translated_cds.faa

Nostoc_sp._T09,ftp://ftp.ncbi.nlm.nih.gov/genomes/all/GCF/002/154/695/GCF_002154695.1_ASM215469v1/GCF_002154695.1_ASM215469v1_translated_cds.faa

Nostocales_cyanobacterium_HT-58-2,ftp://ftp.ncbi.nlm.nih.gov/genomes/all/GCF/002/163/975/GCF_002163975.1_ASM216397v1/GCF_002163975.1_ASM216397v1_translated_cds.faa

Oscillatoria_acuminata_PCC_6304,ftp://ftp.ncbi.nlm.nih.gov/genomes/all/GCF/000/317/105/GCF_000317105.1_ASM31710v1/GCF_000317105.1_ASM31710v1_translated_cds.faa

Oscillatoria_nigro-viridis_PCC_7112,ftp://ftp.ncbi.nlm.nih.gov/genomes/all/GCF/000/317/475/GCF_000317475.1_ASM31747v1/GCF_000317475.1_ASM31747v1_translated_cds.faa

Oscillatoria_sp._PCC_10802,ftp://ftp.ncbi.nlm.nih.gov/genomes/all/GCF/000/332/335/GCF_000332335.1_ASM33233v1/GCF_000332335.1_ASM33233v1_translated_cds.faa

Oscillatoriales_cyanobacterium_CG2_30_40_61,ftp://ftp.ncbi.nlm.nih.gov/genomes/all/GCA/001/873/365/GCA_001873365.1_ASM187336v1/GCA_001873365.1_ASM187336v1_translated_cds.faa

Oscillatoriales_cyanobacterium_CG2_30_44_21,ftp://ftp.ncbi.nlm.nih.gov/genomes/all/GCA/001/873/375/GCA_001873375.1_ASM187337v1/GCA_001873375.1_ASM187337v1_translated_cds.faa

Oscillatoriales_cyanobacterium_JSC-12,ftp://ftp.ncbi.nlm.nih.gov/genomes/all/GCF/000/309/945/GCF_000309945.1_ASM30994v1/GCF_000309945.1_ASM30994v1_translated_cds.faa

Oscillatoriales_cyanobacterium_MTP1,ftp://ftp.ncbi.nlm.nih.gov/genomes/all/GCF/001/482/745/GCF_001482745.2_ASM148274v2/GCF_001482745.2_ASM148274v2_translated_cds.faa

Oscillatoriales_cyanobacterium_USR001,ftp://ftp.ncbi.nlm.nih.gov/genomes/all/GCA/001/698/425/GCA_001698425.1_ASM169842v1/GCA_001698425.1_ASM169842v1_translated_cds.faa

Phormidesmis_priestleyi_Ana,ftp://ftp.ncbi.nlm.nih.gov/genomes/all/GCA/001/314/865/GCA_001314865.1_ASM131486v1/GCA_001314865.1_ASM131486v1_translated_cds.faa

Phormidesmis_priestleyi_BC1401,ftp://ftp.ncbi.nlm.nih.gov/genomes/all/GCF/001/650/195/GCF_001650195.1_ASM165019v1/GCF_001650195.1_ASM165019v1_translated_cds.faa

Phormidesmis_priestleyi_ULC007,ftp://ftp.ncbi.nlm.nih.gov/genomes/all/GCF/001/895/925/GCF_001895925.1_ASM189592v1/GCF_001895925.1_ASM189592v1_translated_cds.faa

Phormidium_ambiguum_IAM_M-71,ftp://ftp.ncbi.nlm.nih.gov/genomes/all/GCF/001/904/725/GCF_001904725.1_ASM190472v1/GCF_001904725.1_ASM190472v1_translated_cds.faa

Phormidium_sp._HE10JO,ftp://ftp.ncbi.nlm.nih.gov/genomes/all/GCF/900/149/785/GCF_900149785.1_Phormidium_lacuna/GCF_900149785.1_Phormidium_lacuna_translated_cds.faa

Phormidium_sp._OSCR,ftp://ftp.ncbi.nlm.nih.gov/genomes/all/GCA/001/314/905/GCA_001314905.1_ASM131490v1/GCA_001314905.1_ASM131490v1_translated_cds.faa

Phormidium_tenue_NIES-30,ftp://ftp.ncbi.nlm.nih.gov/genomes/all/GCF/001/904/775/GCF_001904775.1_ASM190477v1/GCF_001904775.1_ASM190477v1_translated_cds.faa

Phormidium_willei_BDU_130791,ftp://ftp.ncbi.nlm.nih.gov/genomes/all/GCF/001/637/315/GCF_001637315.1_ASM163731v1/GCF_001637315.1_ASM163731v1_translated_cds.faa

Planktothricoides_sp._SR001,ftp://ftp.ncbi.nlm.nih.gov/genomes/all/GCF/001/276/715/GCF_001276715.1_ASM127671v1/GCF_001276715.1_ASM127671v1_translated_cds.faa

Planktothrix_agardhii_NIVA-CYA_126/8,ftp://ftp.ncbi.nlm.nih.gov/genomes/all/GCF/000/710/505/GCF_000710505.1_Plagard2.0/GCF_000710505.1_Plagard2.0_translated_cds.faa

Planktothrix_agardhii_NIVA-CYA_15,ftp://ftp.ncbi.nlm.nih.gov/genomes/all/GCF/000/464/665/GCF_000464665.1_NC15_1/GCF_000464665.1_NC15_1_translated_cds.faa

Planktothrix_agardhii_NIVA-CYA_34,ftp://ftp.ncbi.nlm.nih.gov/genomes/all/GCF/000/464/725/GCF_000464725.1_NC34_1/GCF_000464725.1_NC34_1_translated_cds.faa

Planktothrix_agardhii_NIVA-CYA_56/3,ftp://ftp.ncbi.nlm.nih.gov/genomes/all/GCF/000/464/825/GCF_000464825.1_NC56_3_1/GCF_000464825.1_NC56_3_1_translated_cds.faa

Planktothrix_mougeotii_NIVA-CYA_405,ftp://ftp.ncbi.nlm.nih.gov/genomes/all/GCF/000/464/745/GCF_000464745.1_NC405_1/GCF_000464745.1_NC405_1_translated_cds.faa

Planktothrix_paucivesiculata_PCC_9631,ftp://ftp.ncbi.nlm.nih.gov/genomes/all/GCF/900/009/265/GCF_900009265.1_BBR_PRJEB10994/GCF_900009265.1_BBR_PRJEB10994_translated_cds.faa

Planktothrix_prolifica_NIVA-CYA_406,ftp://ftp.ncbi.nlm.nih.gov/genomes/all/GCF/000/464/765/GCF_000464765.1_NC406_1/GCF_000464765.1_NC406_1_translated_cds.faa

Planktothrix_prolifica_NIVA-CYA_540,ftp://ftp.ncbi.nlm.nih.gov/genomes/all/GCF/000/464/805/GCF_000464805.1_NC540_1/GCF_000464805.1_NC540_1_translated_cds.faa

Planktothrix_prolifica_NIVA-CYA_98,ftp://ftp.ncbi.nlm.nih.gov/genomes/all/GCF/000/464/845/GCF_000464845.1_NC98_2/GCF_000464845.1_NC98_2_translated_cds.faa

Planktothrix_rubescens,ftp://ftp.ncbi.nlm.nih.gov/genomes/all/GCF/900/009/275/GCF_900009275.1_AAI_PRJEB10990/GCF_900009275.1_AAI_PRJEB10990_translated_cds.faa

Planktothrix_rubescens_NIVA-CYA_407,ftp://ftp.ncbi.nlm.nih.gov/genomes/all/GCF/000/464/785/GCF_000464785.1_NC407_1/GCF_000464785.1_NC407_1_translated_cds.faa

Planktothrix_serta_PCC_8927,ftp://ftp.ncbi.nlm.nih.gov/genomes/all/GCF/900/010/725/GCF_900010725.1_BBR_PRJEB10992/GCF_900010725.1_BBR_PRJEB10992_translated_cds.faa

Planktothrix_sp._PCC_11201,ftp://ftp.ncbi.nlm.nih.gov/genomes/all/GCF/900/009/135/GCF_900009135.1_BBR_PRJEB10991_v2/GCF_900009135.1_BBR_PRJEB10991_v2_translated_cds.faa

Planktothrix_tepida_PCC_9214,ftp://ftp.ncbi.nlm.nih.gov/genomes/all/GCF/900/009/145/GCF_900009145.1_BBR_PRJEB10993/GCF_900009145.1_BBR_PRJEB10993_translated_cds.faa

Pleurocapsa_sp._PCC_7319,ftp://ftp.ncbi.nlm.nih.gov/genomes/all/GCF/000/332/195/GCF_000332195.1_ASM33219v1/GCF_000332195.1_ASM33219v1_translated_cds.faa

Pleurocapsa_sp._PCC_7327,ftp://ftp.ncbi.nlm.nih.gov/genomes/all/GCF/000/317/025/GCF_000317025.1_ASM31702v1/GCF_000317025.1_ASM31702v1_translated_cds.faa

Prochlorococcus_marinus,ftp://ftp.ncbi.nlm.nih.gov/genomes/all/GCF/001/180/245/GCF_001180245.1_CLC_assembled_contigs/GCF_001180245.1_CLC_assembled_contigs_translated_cds.faa

Prochlorococcus_marinus_bv._HNLC1,ftp://ftp.ncbi.nlm.nih.gov/genomes/all/GCF/000/218/705/GCF_000218705.1_HNLC1/GCF_000218705.1_HNLC1_protein.faa

Prochlorococcus_marinus_bv._HNLC2,ftp://ftp.ncbi.nlm.nih.gov/genomes/all/GCF/000/218/745/GCF_000218745.1_HNLC2/GCF_000218745.1_HNLC2_protein.faa

Prochlorococcus_marinus_str._AS9601,ftp://ftp.ncbi.nlm.nih.gov/genomes/all/GCF/000/015/645/GCF_000015645.1_ASM1564v1/GCF_000015645.1_ASM1564v1_translated_cds.faa

Prochlorococcus_marinus_str._EQPAC1,ftp://ftp.ncbi.nlm.nih.gov/genomes/all/GCF/000/759/875/GCF_000759875.1_ASM75987v1/GCF_000759875.1_ASM75987v1_translated_cds.faa

Prochlorococcus_marinus_str._GP2,ftp://ftp.ncbi.nlm.nih.gov/genomes/all/GCF/000/759/885/GCF_000759885.1_ASM75988v1/GCF_000759885.1_ASM75988v1_translated_cds.faa

Prochlorococcus_marinus_str._LG,ftp://ftp.ncbi.nlm.nih.gov/genomes/all/GCF/000/760/155/GCF_000760155.1_ASM76015v1/GCF_000760155.1_ASM76015v1_translated_cds.faa

Prochlorococcus_marinus_str._MIT_1312,ftp://ftp.ncbi.nlm.nih.gov/genomes/all/GCF/001/632/005/GCF_001632005.1_ASM163200v1/GCF_001632005.1_ASM163200v1_protein.faa

Prochlorococcus_marinus_str._MIT_1313,ftp://ftp.ncbi.nlm.nih.gov/genomes/all/GCF/001/632/065/GCF_001632065.1_ASM163206v1/GCF_001632065.1_ASM163206v1_protein.faa

Prochlorococcus_marinus_str._MIT_1318,ftp://ftp.ncbi.nlm.nih.gov/genomes/all/GCF/001/632/045/GCF_001632045.1_ASM163204v1/GCF_001632045.1_ASM163204v1_protein.faa

Prochlorococcus_marinus_str._MIT_1320,ftp://ftp.ncbi.nlm.nih.gov/genomes/all/GCF/001/632/075/GCF_001632075.1_ASM163207v1/GCF_001632075.1_ASM163207v1_translated_cds.faa

Prochlorococcus_marinus_str._MIT_1323,ftp://ftp.ncbi.nlm.nih.gov/genomes/all/GCF/001/632/025/GCF_001632025.1_ASM163202v1/GCF_001632025.1_ASM163202v1_translated_cds.faa

Prochlorococcus_marinus_str._MIT_1327,ftp://ftp.ncbi.nlm.nih.gov/genomes/all/GCF/001/632/125/GCF_001632125.1_ASM163212v1/GCF_001632125.1_ASM163212v1_protein.faa

Prochlorococcus_marinus_str._MIT_1342,ftp://ftp.ncbi.nlm.nih.gov/genomes/all/GCF/001/632/145/GCF_001632145.1_ASM163214v1/GCF_001632145.1_ASM163214v1_translated_cds.faa

Prochlorococcus_marinus_str._MIT_9107,ftp://ftp.ncbi.nlm.nih.gov/genomes/all/GCF/000/759/855/GCF_000759855.1_ASM75985v1/GCF_000759855.1_ASM75985v1_translated_cds.faa

Prochlorococcus_marinus_str._MIT_9116,ftp://ftp.ncbi.nlm.nih.gov/genomes/all/GCF/000/759/865/GCF_000759865.1_ASM75986v1/GCF_000759865.1_ASM75986v1_translated_cds.faa

Prochlorococcus_marinus_str._MIT_9123,ftp://ftp.ncbi.nlm.nih.gov/genomes/all/GCF/000/759/935/GCF_000759935.1_ASM75993v1/GCF_000759935.1_ASM75993v1_translated_cds.faa

Prochlorococcus_marinus_str._MIT_9201,ftp://ftp.ncbi.nlm.nih.gov/genomes/all/GCF/000/759/955/GCF_000759955.1_ASM75995v1/GCF_000759955.1_ASM75995v1_translated_cds.faa

Prochlorococcus_marinus_str._MIT_9202,ftp://ftp.ncbi.nlm.nih.gov/genomes/all/GCF/000/158/595/GCF_000158595.1_ASM15859v1/GCF_000158595.1_ASM15859v1_translated_cds.faa

Prochlorococcus_marinus_str._MIT_9211,ftp://ftp.ncbi.nlm.nih.gov/genomes/all/GCF/000/018/585/GCF_000018585.1_ASM1858v1/GCF_000018585.1_ASM1858v1_translated_cds.faa

Prochlorococcus_marinus_str._MIT_9215,ftp://ftp.ncbi.nlm.nih.gov/genomes/all/GCF/000/018/065/GCF_000018065.1_ASM1806v1/GCF_000018065.1_ASM1806v1_translated_cds.faa

Prochlorococcus_marinus_str._MIT_9301,ftp://ftp.ncbi.nlm.nih.gov/genomes/all/GCF/000/015/965/GCF_000015965.1_ASM1596v1/GCF_000015965.1_ASM1596v1_translated_cds.faa

Prochlorococcus_marinus_str._MIT_9302,ftp://ftp.ncbi.nlm.nih.gov/genomes/all/GCF/000/759/975/GCF_000759975.1_ASM75997v1/GCF_000759975.1_ASM75997v1_translated_cds.faa

Prochlorococcus_marinus_str._MIT_9303,ftp://ftp.ncbi.nlm.nih.gov/genomes/all/GCF/000/015/705/GCF_000015705.1_ASM1570v1/GCF_000015705.1_ASM1570v1_protein.faa

Prochlorococcus_marinus_str._MIT_9311,ftp://ftp.ncbi.nlm.nih.gov/genomes/all/GCF/000/760/015/GCF_000760015.1_ASM76001v1/GCF_000760015.1_ASM76001v1_translated_cds.faa

Prochlorococcus_marinus_str._MIT_9312,ftp://ftp.ncbi.nlm.nih.gov/genomes/all/GCF/000/012/645/GCF_000012645.1_ASM1264v1/GCF_000012645.1_ASM1264v1_translated_cds.faa

Prochlorococcus_marinus_str._MIT_9313,ftp://ftp.ncbi.nlm.nih.gov/genomes/all/GCF/000/011/485/GCF_000011485.1_ASM1148v1/GCF_000011485.1_ASM1148v1_translated_cds.faa

Prochlorococcus_marinus_str._MIT_9314,ftp://ftp.ncbi.nlm.nih.gov/genomes/all/GCF/000/760/035/GCF_000760035.1_ASM76003v1/GCF_000760035.1_ASM76003v1_translated_cds.faa

Prochlorococcus_marinus_str._MIT_9321,ftp://ftp.ncbi.nlm.nih.gov/genomes/all/GCF/000/760/055/GCF_000760055.1_ASM76005v1/GCF_000760055.1_ASM76005v1_translated_cds.faa

Prochlorococcus_marinus_str._MIT_9322,ftp://ftp.ncbi.nlm.nih.gov/genomes/all/GCF/000/760/075/GCF_000760075.1_ASM76007v1/GCF_000760075.1_ASM76007v1_translated_cds.faa

Prochlorococcus_marinus_str._MIT_9401,ftp://ftp.ncbi.nlm.nih.gov/genomes/all/GCF/000/760/095/GCF_000760095.1_ASM76009v1/GCF_000760095.1_ASM76009v1_translated_cds.faa

Prochlorococcus_marinus_str._MIT_9515,ftp://ftp.ncbi.nlm.nih.gov/genomes/all/GCF/000/015/665/GCF_000015665.1_ASM1566v1/GCF_000015665.1_ASM1566v1_translated_cds.faa

Prochlorococcus_marinus_str._NATL1A,ftp://ftp.ncbi.nlm.nih.gov/genomes/all/GCF/000/015/685/GCF_000015685.1_ASM1568v1/GCF_000015685.1_ASM1568v1_translated_cds.faa

Prochlorococcus_marinus_str._NATL2A,ftp://ftp.ncbi.nlm.nih.gov/genomes/all/GCF/000/012/465/GCF_000012465.1_ASM1246v1/GCF_000012465.1_ASM1246v1_translated_cds.faa

Prochlorococcus_marinus_str._PAC1,ftp://ftp.ncbi.nlm.nih.gov/genomes/all/GCF/000/760/235/GCF_000760235.1_ASM76023v1/GCF_000760235.1_ASM76023v1_translated_cds.faa

Prochlorococcus_marinus_str._SB,ftp://ftp.ncbi.nlm.nih.gov/genomes/all/GCF/000/760/115/GCF_000760115.1_ASM76011v1/GCF_000760115.1_ASM76011v1_translated_cds.faa

Prochlorococcus_marinus_str._SS2,ftp://ftp.ncbi.nlm.nih.gov/genomes/all/GCF/000/760/255/GCF_000760255.1_ASM76025v1/GCF_000760255.1_ASM76025v1_translated_cds.faa

Prochlorococcus_marinus_str._SS35,ftp://ftp.ncbi.nlm.nih.gov/genomes/all/GCF/000/760/275/GCF_000760275.1_ASM76027v1/GCF_000760275.1_ASM76027v1_translated_cds.faa

Prochlorococcus_marinus_str._SS51,ftp://ftp.ncbi.nlm.nih.gov/genomes/all/GCF/000/760/355/GCF_000760355.1_ASM76035v1/GCF_000760355.1_ASM76035v1_translated_cds.faa

Prochlorococcus_marinus_str._XMU1401,ftp://ftp.ncbi.nlm.nih.gov/genomes/all/GCF/002/812/945/GCF_002812945.1_ASM281294v1/GCF_002812945.1_ASM281294v1_translated_cds.faa

Prochlorococcus_marinus_subsp._marinus_str._CCMP1375,ftp://ftp.ncbi.nlm.nih.gov/genomes/all/GCF/000/007/925/GCF_000007925.1_ASM792v1/GCF_000007925.1_ASM792v1_translated_cds.faa

Prochlorococcus_marinus_subsp._pastoris_str._CCMP1986,ftp://ftp.ncbi.nlm.nih.gov/genomes/all/GCF/000/011/465/GCF_000011465.1_ASM1146v1/GCF_000011465.1_ASM1146v1_translated_cds.faa

Prochlorococcus_sp._HOT208_60m_805A16,ftp://ftp.ncbi.nlm.nih.gov/genomes/all/GCF/002/026/045/GCF_002026045.1_ASM202604v1/GCF_002026045.1_ASM202604v1_translated_cds.faa

Prochlorococcus_sp._HOT208_60m_808G21,ftp://ftp.ncbi.nlm.nih.gov/genomes/all/GCF/002/026/015/GCF_002026015.1_ASM202601v1/GCF_002026015.1_ASM202601v1_translated_cds.faa

Prochlorococcus_sp._HOT208_60m_808M21,ftp://ftp.ncbi.nlm.nih.gov/genomes/all/GCF/002/026/005/GCF_002026005.1_ASM202600v1/GCF_002026005.1_ASM202600v1_translated_cds.faa

Prochlorococcus_sp._HOT208_60m_810B23,ftp://ftp.ncbi.nlm.nih.gov/genomes/all/GCF/002/026/205/GCF_002026205.1_ASM202620v1/GCF_002026205.1_ASM202620v1_translated_cds.faa

Prochlorococcus_sp._HOT208_60m_810P02,ftp://ftp.ncbi.nlm.nih.gov/genomes/all/GCF/002/025/905/GCF_002025905.1_ASM202590v1/GCF_002025905.1_ASM202590v1_translated_cds.faa

Prochlorococcus_sp._HOT208_60m_813B04,ftp://ftp.ncbi.nlm.nih.gov/genomes/all/GCF/002/025/975/GCF_002025975.1_ASM202597v1/GCF_002025975.1_ASM202597v1_translated_cds.faa

Prochlorococcus_sp._HOT208_60m_813E23,ftp://ftp.ncbi.nlm.nih.gov/genomes/all/GCF/002/026/175/GCF_002026175.1_ASM202617v1/GCF_002026175.1_ASM202617v1_translated_cds.faa

Prochlorococcus_sp._HOT208_60m_813G15,ftp://ftp.ncbi.nlm.nih.gov/genomes/all/GCF/002/026/165/GCF_002026165.1_ASM202616v1/GCF_002026165.1_ASM202616v1_translated_cds.faa

Prochlorococcus_sp._HOT208_60m_813I02,ftp://ftp.ncbi.nlm.nih.gov/genomes/all/GCF/002/025/865/GCF_002025865.1_ASM202586v1/GCF_002025865.1_ASM202586v1_translated_cds.faa

Prochlorococcus_sp._HOT208_60m_813L03,ftp://ftp.ncbi.nlm.nih.gov/genomes/all/GCF/002/026/145/GCF_002026145.1_ASM202614v1/GCF_002026145.1_ASM202614v1_translated_cds.faa

Prochlorococcus_sp._HOT208_60m_813O14,ftp://ftp.ncbi.nlm.nih.gov/genomes/all/GCF/002/025/945/GCF_002025945.1_ASM202594v1/GCF_002025945.1_ASM202594v1_translated_cds.faa

Prochlorococcus_sp._HOT212_60m_824C06,ftp://ftp.ncbi.nlm.nih.gov/genomes/all/GCF/002/026/105/GCF_002026105.1_ASM202610v1/GCF_002026105.1_ASM202610v1_translated_cds.faa

Prochlorococcus_sp._HOT212_60m_824E10,ftp://ftp.ncbi.nlm.nih.gov/genomes/all/GCF/002/026/185/GCF_002026185.1_ASM202618v1/GCF_002026185.1_ASM202618v1_translated_cds.faa

Prochlorococcus_sp._HOT212_60m_826P21,ftp://ftp.ncbi.nlm.nih.gov/genomes/all/GCF/002/026/065/GCF_002026065.1_ASM202606v1/GCF_002026065.1_ASM202606v1_translated_cds.faa

Prochlorococcus_sp._HOT_208_60,ftp://ftp.ncbi.nlm.nih.gov/genomes/all/GCA/002/156/565/GCA_002156565.1_ASM215656v1/GCA_002156565.1_ASM215656v1_translated_cds.faa

Prochlorococcus_sp._MED105,ftp://ftp.ncbi.nlm.nih.gov/genomes/all/GCA/002/692/385/GCA_002692385.1_ASM269238v1/GCA_002692385.1_ASM269238v1_translated_cds.faa

Prochlorococcus_sp._MED630,ftp://ftp.ncbi.nlm.nih.gov/genomes/all/GCA/002/689/785/GCA_002689785.1_ASM268978v1/GCA_002689785.1_ASM268978v1_translated_cds.faa

Prochlorococcus_sp._MIT_0601,ftp://ftp.ncbi.nlm.nih.gov/genomes/all/GCF/000/760/175/GCF_000760175.1_ASM76017v1/GCF_000760175.1_ASM76017v1_translated_cds.faa

Prochlorococcus_sp._MIT_0602,ftp://ftp.ncbi.nlm.nih.gov/genomes/all/GCF/000/760/195/GCF_000760195.1_ASM76019v1/GCF_000760195.1_ASM76019v1_translated_cds.faa

Prochlorococcus_sp._MIT_0603,ftp://ftp.ncbi.nlm.nih.gov/genomes/all/GCF/000/760/215/GCF_000760215.1_ASM76021v1/GCF_000760215.1_ASM76021v1_translated_cds.faa

Prochlorococcus_sp._MIT_0604,ftp://ftp.ncbi.nlm.nih.gov/genomes/all/GCF/000/757/845/GCF_000757845.1_ASM75784v1/GCF_000757845.1_ASM75784v1_translated_cds.faa

Prochlorococcus_sp._MIT_0701,ftp://ftp.ncbi.nlm.nih.gov/genomes/all/GCF/000/760/295/GCF_000760295.1_ASM76029v1/GCF_000760295.1_ASM76029v1_translated_cds.faa

Prochlorococcus_sp._MIT_0702,ftp://ftp.ncbi.nlm.nih.gov/genomes/all/GCF/000/760/315/GCF_000760315.1_ASM76031v1/GCF_000760315.1_ASM76031v1_translated_cds.faa

Prochlorococcus_sp._MIT_0703,ftp://ftp.ncbi.nlm.nih.gov/genomes/all/GCF/000/760/335/GCF_000760335.1_ASM76033v1/GCF_000760335.1_ASM76033v1_translated_cds.faa

Prochlorococcus_sp._MIT_0801,ftp://ftp.ncbi.nlm.nih.gov/genomes/all/GCF/000/757/865/GCF_000757865.1_ASM75786v1/GCF_000757865.1_ASM75786v1_translated_cds.faa

Prochlorococcus_sp._MIT_1303,ftp://ftp.ncbi.nlm.nih.gov/genomes/all/GCF/001/631/965/GCF_001631965.1_ASM163196v1/GCF_001631965.1_ASM163196v1_translated_cds.faa

Prochlorococcus_sp._MIT_1306,ftp://ftp.ncbi.nlm.nih.gov/genomes/all/GCF/001/631/985/GCF_001631985.1_ASM163198v1/GCF_001631985.1_ASM163198v1_translated_cds.faa

Prochlorococcus_sp._RS01,ftp://ftp.ncbi.nlm.nih.gov/genomes/all/GCF/001/989/435/GCF_001989435.1_ASM198943v1/GCF_001989435.1_ASM198943v1_translated_cds.faa

Prochlorococcus_sp._RS04,ftp://ftp.ncbi.nlm.nih.gov/genomes/all/GCF/001/989/455/GCF_001989455.1_ASM198945v1/GCF_001989455.1_ASM198945v1_translated_cds.faa

Prochlorococcus_sp._RS50,ftp://ftp.ncbi.nlm.nih.gov/genomes/all/GCF/001/989/415/GCF_001989415.1_ASM198941v1/GCF_001989415.1_ASM198941v1_translated_cds.faa

Prochlorococcus_sp._SP3034,ftp://ftp.ncbi.nlm.nih.gov/genomes/all/GCA/002/720/205/GCA_002720205.1_ASM272020v1/GCA_002720205.1_ASM272020v1_translated_cds.faa

Prochlorococcus_sp._SS52,ftp://ftp.ncbi.nlm.nih.gov/genomes/all/GCF/000/760/375/GCF_000760375.1_ASM76037v1/GCF_000760375.1_ASM76037v1_translated_cds.faa

Prochlorococcus_sp._TMED223,ftp://ftp.ncbi.nlm.nih.gov/genomes/all/GCA/002/169/575/GCA_002169575.2_ASM216957v2/GCA_002169575.2_ASM216957v2_translated_cds.faa

Prochlorococcus_sp._W10,ftp://ftp.ncbi.nlm.nih.gov/genomes/all/GCF/000/291/845/GCF_000291845.1_ProW10_v1.0/GCF_000291845.1_ProW10_v1.0_protein.faa

Prochlorococcus_sp._W11,ftp://ftp.ncbi.nlm.nih.gov/genomes/all/GCF/000/291/945/GCF_000291945.1_ProW11_v1.0/GCF_000291945.1_ProW11_v1.0_protein.faa

Prochlorococcus_sp._W12,ftp://ftp.ncbi.nlm.nih.gov/genomes/all/GCF/000/291/965/GCF_000291965.1_ProW12_v1.0/GCF_000291965.1_ProW12_v1.0_protein.faa

Prochlorococcus_sp._W2,ftp://ftp.ncbi.nlm.nih.gov/genomes/all/GCF/000/291/885/GCF_000291885.1_ProW2_v1.0/GCF_000291885.1_ProW2_v1.0_protein.faa

Prochlorococcus_sp._W3,ftp://ftp.ncbi.nlm.nih.gov/genomes/all/GCF/000/291/905/GCF_000291905.1_ProW3_v1.0/GCF_000291905.1_ProW3_v1.0_protein.faa

Prochlorococcus_sp._W4,ftp://ftp.ncbi.nlm.nih.gov/genomes/all/GCF/000/291/785/GCF_000291785.1_ProW4_v1.0/GCF_000291785.1_ProW4_v1.0_protein.faa

Prochlorococcus_sp._W7,ftp://ftp.ncbi.nlm.nih.gov/genomes/all/GCF/000/291/805/GCF_000291805.1_ProW7_v1.0/GCF_000291805.1_ProW7_v1.0_protein.faa

Prochlorococcus_sp._W8,ftp://ftp.ncbi.nlm.nih.gov/genomes/all/GCF/000/291/825/GCF_000291825.1_ProW8_v1.0/GCF_000291825.1_ProW8_v1.0_protein.faa

Prochlorococcus_sp._W9,ftp://ftp.ncbi.nlm.nih.gov/genomes/all/GCF/000/291/925/GCF_000291925.1_ProW9_v1.0/GCF_000291925.1_ProW9_v1.0_protein.faa

Prochlorococcus_sp._scB241_526B17,ftp://ftp.ncbi.nlm.nih.gov/genomes/all/GCF/000/633/975/GCF_000633975.1_De_novo_assembly_of_single-cell_genomes/GCF_000633975.1_De_novo_assembly_of_single-cell_genomes_protein.faa

Prochlorococcus_sp._scB241_526B19,ftp://ftp.ncbi.nlm.nih.gov/genomes/all/GCF/000/634/615/GCF_000634615.1_De_novo_assembly_of_single-cell_genomes/GCF_000634615.1_De_novo_assembly_of_single-cell_genomes_protein.faa

Prochlorococcus_sp._scB241_526B22,ftp://ftp.ncbi.nlm.nih.gov/genomes/all/GCF/000/634/635/GCF_000634635.1_De_novo_assembly_of_single-cell_genomes/GCF_000634635.1_De_novo_assembly_of_single-cell_genomes_protein.faa

Prochlorococcus_sp._scB241_526D20,ftp://ftp.ncbi.nlm.nih.gov/genomes/all/GCF/000/635/275/GCF_000635275.1_De_novo_assembly_of_single-cell_genomes/GCF_000635275.1_De_novo_assembly_of_single-cell_genomes_protein.faa

Prochlorococcus_sp._scB241_526K3,ftp://ftp.ncbi.nlm.nih.gov/genomes/all/GCF/000/633/995/GCF_000633995.1_De_novo_assembly_of_single-cell_genomes/GCF_000633995.1_De_novo_assembly_of_single-cell_genomes_protein.faa

Prochlorococcus_sp._scB241_526N5,ftp://ftp.ncbi.nlm.nih.gov/genomes/all/GCF/000/634/655/GCF_000634655.1_De_novo_assembly_of_single-cell_genomes/GCF_000634655.1_De_novo_assembly_of_single-cell_genomes_protein.faa

Prochlorococcus_sp._scB241_526N9,ftp://ftp.ncbi.nlm.nih.gov/genomes/all/GCF/000/634/015/GCF_000634015.1_De_novo_assembly_of_single-cell_genomes/GCF_000634015.1_De_novo_assembly_of_single-cell_genomes_protein.faa

Prochlorococcus_sp._scB241_527E14,ftp://ftp.ncbi.nlm.nih.gov/genomes/all/GCF/000/634/035/GCF_000634035.1_De_novo_assembly_of_single-cell_genomes/GCF_000634035.1_De_novo_assembly_of_single-cell_genomes_protein.faa

Prochlorococcus_sp._scB241_527E15,ftp://ftp.ncbi.nlm.nih.gov/genomes/all/GCF/000/634/675/GCF_000634675.1_De_novo_assembly_of_single-cell_genomes/GCF_000634675.1_De_novo_assembly_of_single-cell_genomes_protein.faa

Prochlorococcus_sp._scB241_527G5,ftp://ftp.ncbi.nlm.nih.gov/genomes/all/GCF/000/635/295/GCF_000635295.1_De_novo_assembly_of_single-cell_genomes/GCF_000635295.1_De_novo_assembly_of_single-cell_genomes_protein.faa

Prochlorococcus_sp._scB241_527I9,ftp://ftp.ncbi.nlm.nih.gov/genomes/all/GCF/000/635/315/GCF_000635315.1_De_novo_assembly_of_single-cell_genomes/GCF_000635315.1_De_novo_assembly_of_single-cell_genomes_protein.faa

Prochlorococcus_sp._scB241_527L15,ftp://ftp.ncbi.nlm.nih.gov/genomes/all/GCF/000/634/695/GCF_000634695.1_De_novo_assembly_of_single-cell_genomes/GCF_000634695.1_De_novo_assembly_of_single-cell_genomes_protein.faa

Prochlorococcus_sp._scB241_527L16,ftp://ftp.ncbi.nlm.nih.gov/genomes/all/GCF/000/635/335/GCF_000635335.1_De_novo_assembly_of_single-cell_genomes/GCF_000635335.1_De_novo_assembly_of_single-cell_genomes_protein.faa

Prochlorococcus_sp._scB241_527L22,ftp://ftp.ncbi.nlm.nih.gov/genomes/all/GCF/000/635/355/GCF_000635355.1_De_novo_assembly_of_single-cell_genomes/GCF_000635355.1_De_novo_assembly_of_single-cell_genomes_protein.faa

Prochlorococcus_sp._scB241_527N11,ftp://ftp.ncbi.nlm.nih.gov/genomes/all/GCF/000/634/055/GCF_000634055.1_De_novo_assembly_of_single-cell_genomes/GCF_000634055.1_De_novo_assembly_of_single-cell_genomes_protein.faa

Prochlorococcus_sp._scB241_527P5,ftp://ftp.ncbi.nlm.nih.gov/genomes/all/GCF/000/635/375/GCF_000635375.1_De_novo_assembly_of_single-cell_genomes/GCF_000635375.1_De_novo_assembly_of_single-cell_genomes_protein.faa

Prochlorococcus_sp._scB241_528J14,ftp://ftp.ncbi.nlm.nih.gov/genomes/all/GCF/000/634/075/GCF_000634075.1_De_novo_assembly_of_single-cell_genomes/GCF_000634075.1_De_novo_assembly_of_single-cell_genomes_protein.faa

Prochlorococcus_sp._scB241_528J8,ftp://ftp.ncbi.nlm.nih.gov/genomes/all/GCF/000/634/095/GCF_000634095.1_De_novo_assembly_of_single-cell_genomes/GCF_000634095.1_De_novo_assembly_of_single-cell_genomes_protein.faa

Prochlorococcus_sp._scB241_528K19,ftp://ftp.ncbi.nlm.nih.gov/genomes/all/GCF/000/634/715/GCF_000634715.1_De_novo_assembly_of_single-cell_genomes/GCF_000634715.1_De_novo_assembly_of_single-cell_genomes_protein.faa

Prochlorococcus_sp._scB241_528N17,ftp://ftp.ncbi.nlm.nih.gov/genomes/all/GCF/000/634/735/GCF_000634735.1_De_novo_assembly_of_single-cell_genomes/GCF_000634735.1_De_novo_assembly_of_single-cell_genomes_protein.faa

Prochlorococcus_sp._scB241_528N20,ftp://ftp.ncbi.nlm.nih.gov/genomes/all/GCF/000/634/755/GCF_000634755.1_De_novo_assembly_of_single-cell_genomes/GCF_000634755.1_De_novo_assembly_of_single-cell_genomes_protein.faa

Prochlorococcus_sp._scB241_528N8,ftp://ftp.ncbi.nlm.nih.gov/genomes/all/GCF/000/634/775/GCF_000634775.1_De_novo_assembly_of_single-cell_genomes/GCF_000634775.1_De_novo_assembly_of_single-cell_genomes_protein.faa

Prochlorococcus_sp._scB241_528O2,ftp://ftp.ncbi.nlm.nih.gov/genomes/all/GCF/000/634/115/GCF_000634115.1_De_novo_assembly_of_single-cell_genomes/GCF_000634115.1_De_novo_assembly_of_single-cell_genomes_protein.faa

Prochlorococcus_sp._scB241_528P14,ftp://ftp.ncbi.nlm.nih.gov/genomes/all/GCF/000/634/135/GCF_000634135.1_De_novo_assembly_of_single-cell_genomes/GCF_000634135.1_De_novo_assembly_of_single-cell_genomes_protein.faa

Prochlorococcus_sp._scB241_528P18,ftp://ftp.ncbi.nlm.nih.gov/genomes/all/GCF/000/635/395/GCF_000635395.1_De_novo_assembly_of_single-cell_genomes/GCF_000635395.1_De_novo_assembly_of_single-cell_genomes_protein.faa

Prochlorococcus_sp._scB241_529B19,ftp://ftp.ncbi.nlm.nih.gov/genomes/all/GCF/000/635/415/GCF_000635415.1_De_novo_assembly_of_single-cell_genomes/GCF_000635415.1_De_novo_assembly_of_single-cell_genomes_protein.faa

Prochlorococcus_sp._scB241_529C4,ftp://ftp.ncbi.nlm.nih.gov/genomes/all/GCF/000/634/155/GCF_000634155.1_De_novo_assembly_of_single-cell_genomes/GCF_000634155.1_De_novo_assembly_of_single-cell_genomes_protein.faa

Prochlorococcus_sp._scB241_529D18,ftp://ftp.ncbi.nlm.nih.gov/genomes/all/GCF/000/634/175/GCF_000634175.1_De_novo_assembly_of_single-cell_genomes/GCF_000634175.1_De_novo_assembly_of_single-cell_genomes_protein.faa

Prochlorococcus_sp._scB241_529J11,ftp://ftp.ncbi.nlm.nih.gov/genomes/all/GCF/000/635/435/GCF_000635435.1_De_novo_assembly_of_single-cell_genomes/GCF_000635435.1_De_novo_assembly_of_single-cell_genomes_protein.faa

Prochlorococcus_sp._scB241_529J15,ftp://ftp.ncbi.nlm.nih.gov/genomes/all/GCF/000/634/795/GCF_000634795.1_De_novo_assembly_of_single-cell_genomes/GCF_000634795.1_De_novo_assembly_of_single-cell_genomes_protein.faa

Prochlorococcus_sp._scB241_529J16,ftp://ftp.ncbi.nlm.nih.gov/genomes/all/GCF/000/635/455/GCF_000635455.1_De_novo_assembly_of_single-cell_genomes/GCF_000635455.1_De_novo_assembly_of_single-cell_genomes_protein.faa

Prochlorococcus_sp._scB241_529O19,ftp://ftp.ncbi.nlm.nih.gov/genomes/all/GCF/000/635/475/GCF_000635475.1_De_novo_assembly_of_single-cell_genomes/GCF_000635475.1_De_novo_assembly_of_single-cell_genomes_protein.faa

Prochlorococcus_sp._scB243_495D8,ftp://ftp.ncbi.nlm.nih.gov/genomes/all/GCF/000/634/195/GCF_000634195.1_De_novo_assembly_of_single-cell_genomes/GCF_000634195.1_De_novo_assembly_of_single-cell_genomes_protein.faa

Prochlorococcus_sp._scB243_495G23,ftp://ftp.ncbi.nlm.nih.gov/genomes/all/GCF/000/634/815/GCF_000634815.1_De_novo_assembly_of_single-cell_genomes/GCF_000634815.1_De_novo_assembly_of_single-cell_genomes_protein.faa

Prochlorococcus_sp._scB243_495I8,ftp://ftp.ncbi.nlm.nih.gov/genomes/all/GCF/000/634/835/GCF_000634835.1_De_novo_assembly_of_single-cell_genomes/GCF_000634835.1_De_novo_assembly_of_single-cell_genomes_protein.faa

Prochlorococcus_sp._scB243_495K23,ftp://ftp.ncbi.nlm.nih.gov/genomes/all/GCF/000/635/495/GCF_000635495.1_De_novo_assembly_of_single-cell_genomes/GCF_000635495.1_De_novo_assembly_of_single-cell_genomes_protein.faa

Prochlorococcus_sp._scB243_495L20,ftp://ftp.ncbi.nlm.nih.gov/genomes/all/GCF/000/634/215/GCF_000634215.1_De_novo_assembly_of_single-cell_genomes/GCF_000634215.1_De_novo_assembly_of_single-cell_genomes_protein.faa

Prochlorococcus_sp._scB243_495N16,ftp://ftp.ncbi.nlm.nih.gov/genomes/all/GCF/000/634/235/GCF_000634235.1_De_novo_assembly_of_single-cell_genomes/GCF_000634235.1_De_novo_assembly_of_single-cell_genomes_protein.faa

Prochlorococcus_sp._scB243_495N3,ftp://ftp.ncbi.nlm.nih.gov/genomes/all/GCF/000/635/515/GCF_000635515.1_De_novo_assembly_of_single-cell_genomes/GCF_000635515.1_De_novo_assembly_of_single-cell_genomes_protein.faa

Prochlorococcus_sp._scB243_495N4,ftp://ftp.ncbi.nlm.nih.gov/genomes/all/GCF/000/634/855/GCF_000634855.1_De_novo_assembly_of_single-cell_genomes/GCF_000634855.1_De_novo_assembly_of_single-cell_genomes_protein.faa

Prochlorococcus_sp._scB243_495P20,ftp://ftp.ncbi.nlm.nih.gov/genomes/all/GCF/000/634/875/GCF_000634875.1_De_novo_assembly_of_single-cell_genomes/GCF_000634875.1_De_novo_assembly_of_single-cell_genomes_protein.faa

Prochlorococcus_sp._scB243_496A2,ftp://ftp.ncbi.nlm.nih.gov/genomes/all/GCF/000/634/255/GCF_000634255.1_De_novo_assembly_of_single-cell_genomes/GCF_000634255.1_De_novo_assembly_of_single-cell_genomes_protein.faa

Prochlorococcus_sp._scB243_496E10,ftp://ftp.ncbi.nlm.nih.gov/genomes/all/GCF/000/634/895/GCF_000634895.1_De_novo_assembly_of_single-cell_genomes/GCF_000634895.1_De_novo_assembly_of_single-cell_genomes_protein.faa

Prochlorococcus_sp._scB243_496G15,ftp://ftp.ncbi.nlm.nih.gov/genomes/all/GCF/000/634/915/GCF_000634915.1_De_novo_assembly_of_single-cell_genomes/GCF_000634915.1_De_novo_assembly_of_single-cell_genomes_protein.faa

Prochlorococcus_sp._scB243_496M6,ftp://ftp.ncbi.nlm.nih.gov/genomes/all/GCF/000/635/535/GCF_000635535.1_De_novo_assembly_of_single-cell_genomes/GCF_000635535.1_De_novo_assembly_of_single-cell_genomes_protein.faa

Prochlorococcus_sp._scB243_496N4,ftp://ftp.ncbi.nlm.nih.gov/genomes/all/GCF/000/635/555/GCF_000635555.1_De_novo_assembly_of_single-cell_genomes/GCF_000635555.1_De_novo_assembly_of_single-cell_genomes_protein.faa

Prochlorococcus_sp._scB243_497E17,ftp://ftp.ncbi.nlm.nih.gov/genomes/all/GCF/000/635/575/GCF_000635575.1_De_novo_assembly_of_single-cell_genomes/GCF_000635575.1_De_novo_assembly_of_single-cell_genomes_protein.faa

Prochlorococcus_sp._scB243_497I20,ftp://ftp.ncbi.nlm.nih.gov/genomes/all/GCF/000/634/935/GCF_000634935.1_De_novo_assembly_of_single-cell_genomes/GCF_000634935.1_De_novo_assembly_of_single-cell_genomes_protein.faa

Prochlorococcus_sp._scB243_497J18,ftp://ftp.ncbi.nlm.nih.gov/genomes/all/GCF/000/634/275/GCF_000634275.1_De_novo_assembly_of_single-cell_genomes/GCF_000634275.1_De_novo_assembly_of_single-cell_genomes_protein.faa

Prochlorococcus_sp._scB243_497N18,ftp://ftp.ncbi.nlm.nih.gov/genomes/all/GCF/000/634/955/GCF_000634955.1_De_novo_assembly_of_single-cell_genomes/GCF_000634955.1_De_novo_assembly_of_single-cell_genomes_protein.faa

Prochlorococcus_sp._scB243_498A3,ftp://ftp.ncbi.nlm.nih.gov/genomes/all/GCF/000/634/295/GCF_000634295.1_De_novo_assembly_of_single-cell_genomes/GCF_000634295.1_De_novo_assembly_of_single-cell_genomes_protein.faa

Prochlorococcus_sp._scB243_498B22,ftp://ftp.ncbi.nlm.nih.gov/genomes/all/GCF/000/634/975/GCF_000634975.1_De_novo_assembly_of_single-cell_genomes/GCF_000634975.1_De_novo_assembly_of_single-cell_genomes_protein.faa

Prochlorococcus_sp._scB243_498B23,ftp://ftp.ncbi.nlm.nih.gov/genomes/all/GCF/000/635/595/GCF_000635595.1_De_novo_assembly_of_single-cell_genomes/GCF_000635595.1_De_novo_assembly_of_single-cell_genomes_protein.faa

Prochlorococcus_sp._scB243_498C16,ftp://ftp.ncbi.nlm.nih.gov/genomes/all/GCF/000/634/995/GCF_000634995.1_De_novo_assembly_of_single-cell_genomes/GCF_000634995.1_De_novo_assembly_of_single-cell_genomes_protein.faa

Prochlorococcus_sp._scB243_498F21,ftp://ftp.ncbi.nlm.nih.gov/genomes/all/GCF/000/635/615/GCF_000635615.1_De_novo_assembly_of_single-cell_genomes/GCF_000635615.1_De_novo_assembly_of_single-cell_genomes_protein.faa

Prochlorococcus_sp._scB243_498G3,ftp://ftp.ncbi.nlm.nih.gov/genomes/all/GCF/000/635/635/GCF_000635635.1_De_novo_assembly_of_single-cell_genomes/GCF_000635635.1_De_novo_assembly_of_single-cell_genomes_protein.faa

Prochlorococcus_sp._scB243_498I20,ftp://ftp.ncbi.nlm.nih.gov/genomes/all/GCF/000/635/015/GCF_000635015.1_De_novo_assembly_of_single-cell_genomes/GCF_000635015.1_De_novo_assembly_of_single-cell_genomes_protein.faa

Prochlorococcus_sp._scB243_498J20,ftp://ftp.ncbi.nlm.nih.gov/genomes/all/GCF/000/634/315/GCF_000634315.1_De_novo_assembly_of_single-cell_genomes/GCF_000634315.1_De_novo_assembly_of_single-cell_genomes_protein.faa

Prochlorococcus_sp._scB243_498L10,ftp://ftp.ncbi.nlm.nih.gov/genomes/all/GCF/000/635/035/GCF_000635035.1_De_novo_assembly_of_single-cell_genomes/GCF_000635035.1_De_novo_assembly_of_single-cell_genomes_protein.faa

Prochlorococcus_sp._scB243_498M14,ftp://ftp.ncbi.nlm.nih.gov/genomes/all/GCF/000/635/655/GCF_000635655.1_De_novo_assembly_of_single-cell_genomes/GCF_000635655.1_De_novo_assembly_of_single-cell_genomes_protein.faa

Prochlorococcus_sp._scB243_498N4,ftp://ftp.ncbi.nlm.nih.gov/genomes/all/GCF/000/634/335/GCF_000634335.1_De_novo_assembly_of_single-cell_genomes/GCF_000634335.1_De_novo_assembly_of_single-cell_genomes_protein.faa

Prochlorococcus_sp._scB243_498N8,ftp://ftp.ncbi.nlm.nih.gov/genomes/all/GCF/000/634/355/GCF_000634355.1_De_novo_assembly_of_single-cell_genomes/GCF_000634355.1_De_novo_assembly_of_single-cell_genomes_protein.faa

Prochlorococcus_sp._scB243_498P15,ftp://ftp.ncbi.nlm.nih.gov/genomes/all/GCF/000/635/675/GCF_000635675.1_De_novo_assembly_of_single-cell_genomes/GCF_000635675.1_De_novo_assembly_of_single-cell_genomes_protein.faa

Prochlorococcus_sp._scB243_498P3,ftp://ftp.ncbi.nlm.nih.gov/genomes/all/GCF/000/635/695/GCF_000635695.1_De_novo_assembly_of_single-cell_genomes/GCF_000635695.1_De_novo_assembly_of_single-cell_genomes_protein.faa

Prochlorococcus_sp._scB245a_518A17,ftp://ftp.ncbi.nlm.nih.gov/genomes/all/GCF/000/634/375/GCF_000634375.1_De_novo_assembly_of_single-cell_genomes/GCF_000634375.1_De_novo_assembly_of_single-cell_genomes_protein.faa

Prochlorococcus_sp._scB245a_518A6,ftp://ftp.ncbi.nlm.nih.gov/genomes/all/GCF/000/635/055/GCF_000635055.1_De_novo_assembly_of_single-cell_genomes/GCF_000635055.1_De_novo_assembly_of_single-cell_genomes_protein.faa

Prochlorococcus_sp._scB245a_518E10,ftp://ftp.ncbi.nlm.nih.gov/genomes/all/GCF/000/635/075/GCF_000635075.1_De_novo_assembly_of_single-cell_genomes/GCF_000635075.1_De_novo_assembly_of_single-cell_genomes_protein.faa

Prochlorococcus_sp._scB245a_518I6,ftp://ftp.ncbi.nlm.nih.gov/genomes/all/GCF/000/635/715/GCF_000635715.1_De_novo_assembly_of_single-cell_genomes/GCF_000635715.1_De_novo_assembly_of_single-cell_genomes_protein.faa

Prochlorococcus_sp._scB245a_518J7,ftp://ftp.ncbi.nlm.nih.gov/genomes/all/GCF/000/635/095/GCF_000635095.1_De_novo_assembly_of_single-cell_genomes/GCF_000635095.1_De_novo_assembly_of_single-cell_genomes_protein.faa

Prochlorococcus_sp._scB245a_518K17,ftp://ftp.ncbi.nlm.nih.gov/genomes/all/GCF/000/635/115/GCF_000635115.1_De_novo_assembly_of_single-cell_genomes/GCF_000635115.1_De_novo_assembly_of_single-cell_genomes_protein.faa

Prochlorococcus_sp._scB245a_518O7,ftp://ftp.ncbi.nlm.nih.gov/genomes/all/GCF/000/635/135/GCF_000635135.1_De_novo_assembly_of_single-cell_genomes/GCF_000635135.1_De_novo_assembly_of_single-cell_genomes_protein.faa

Prochlorococcus_sp._scB245a_519A13,ftp://ftp.ncbi.nlm.nih.gov/genomes/all/GCF/000/634/415/GCF_000634415.1_De_novo_assembly_of_single-cell_genomes/GCF_000634415.1_De_novo_assembly_of_single-cell_genomes_protein.faa

Prochlorococcus_sp._scB245a_519B7,ftp://ftp.ncbi.nlm.nih.gov/genomes/all/GCF/000/634/435/GCF_000634435.1_De_novo_assembly_of_single-cell_genomes/GCF_000634435.1_De_novo_assembly_of_single-cell_genomes_protein.faa

Prochlorococcus_sp._scB245a_519C7,ftp://ftp.ncbi.nlm.nih.gov/genomes/all/GCF/000/635/155/GCF_000635155.1_De_novo_assembly_of_single-cell_genomes/GCF_000635155.1_De_novo_assembly_of_single-cell_genomes_protein.faa

Prochlorococcus_sp._scB245a_519D13,ftp://ftp.ncbi.nlm.nih.gov/genomes/all/GCF/000/635/735/GCF_000635735.1_De_novo_assembly_of_single-cell_genomes/GCF_000635735.1_De_novo_assembly_of_single-cell_genomes_protein.faa

Prochlorococcus_sp._scB245a_519E23,ftp://ftp.ncbi.nlm.nih.gov/genomes/all/GCF/000/635/175/GCF_000635175.1_De_novo_assembly_of_single-cell_genomes/GCF_000635175.1_De_novo_assembly_of_single-cell_genomes_protein.faa

Prochlorococcus_sp._scB245a_519G16,ftp://ftp.ncbi.nlm.nih.gov/genomes/all/GCF/000/635/755/GCF_000635755.1_De_novo_assembly_of_single-cell_genomes/GCF_000635755.1_De_novo_assembly_of_single-cell_genomes_protein.faa

Prochlorococcus_sp._scB245a_519L21,ftp://ftp.ncbi.nlm.nih.gov/genomes/all/GCF/000/635/195/GCF_000635195.1_De_novo_assembly_of_single-cell_genomes/GCF_000635195.1_De_novo_assembly_of_single-cell_genomes_protein.faa

Prochlorococcus_sp._scB245a_519O11,ftp://ftp.ncbi.nlm.nih.gov/genomes/all/GCF/000/635/775/GCF_000635775.1_De_novo_assembly_of_single-cell_genomes/GCF_000635775.1_De_novo_assembly_of_single-cell_genomes_protein.faa

Prochlorococcus_sp._scB245a_519O21,ftp://ftp.ncbi.nlm.nih.gov/genomes/all/GCF/000/634/455/GCF_000634455.1_De_novo_assembly_of_single-cell_genomes/GCF_000634455.1_De_novo_assembly_of_single-cell_genomes_protein.faa

Prochlorococcus_sp._scB245a_520B18,ftp://ftp.ncbi.nlm.nih.gov/genomes/all/GCF/000/634/475/GCF_000634475.1_De_novo_assembly_of_single-cell_genomes/GCF_000634475.1_De_novo_assembly_of_single-cell_genomes_protein.faa

Prochlorococcus_sp._scB245a_520D2,ftp://ftp.ncbi.nlm.nih.gov/genomes/all/GCF/000/634/495/GCF_000634495.1_De_novo_assembly_of_single-cell_genomes/GCF_000634495.1_De_novo_assembly_of_single-cell_genomes_protein.faa

Prochlorococcus_sp._scB245a_520E22,ftp://ftp.ncbi.nlm.nih.gov/genomes/all/GCF/000/635/795/GCF_000635795.1_De_novo_assembly_of_single-cell_genomes/GCF_000635795.1_De_novo_assembly_of_single-cell_genomes_protein.faa

Prochlorococcus_sp._scB245a_520F22,ftp://ftp.ncbi.nlm.nih.gov/genomes/all/GCF/000/635/815/GCF_000635815.1_De_novo_assembly_of_single-cell_genomes/GCF_000635815.1_De_novo_assembly_of_single-cell_genomes_protein.faa

Prochlorococcus_sp._scB245a_520K10,ftp://ftp.ncbi.nlm.nih.gov/genomes/all/GCF/000/634/515/GCF_000634515.1_De_novo_assembly_of_single-cell_genomes/GCF_000634515.1_De_novo_assembly_of_single-cell_genomes_protein.faa

Prochlorococcus_sp._scB245a_520M11,ftp://ftp.ncbi.nlm.nih.gov/genomes/all/GCF/000/634/535/GCF_000634535.1_De_novo_assembly_of_single-cell_genomes/GCF_000634535.1_De_novo_assembly_of_single-cell_genomes_protein.faa

Prochlorococcus_sp._scB245a_521B10,ftp://ftp.ncbi.nlm.nih.gov/genomes/all/GCF/000/635/835/GCF_000635835.1_De_novo_assembly_of_single-cell_genomes/GCF_000635835.1_De_novo_assembly_of_single-cell_genomes_protein.faa

Prochlorococcus_sp._scB245a_521C8,ftp://ftp.ncbi.nlm.nih.gov/genomes/all/GCF/000/635/215/GCF_000635215.1_De_novo_assembly_of_single-cell_genomes/GCF_000635215.1_De_novo_assembly_of_single-cell_genomes_protein.faa

Prochlorococcus_sp._scB245a_521K15,ftp://ftp.ncbi.nlm.nih.gov/genomes/all/GCF/000/635/235/GCF_000635235.1_De_novo_assembly_of_single-cell_genomes/GCF_000635235.1_De_novo_assembly_of_single-cell_genomes_protein.faa

Prochlorococcus_sp._scB245a_521M10,ftp://ftp.ncbi.nlm.nih.gov/genomes/all/GCF/000/635/855/GCF_000635855.1_De_novo_assembly_of_single-cell_genomes/GCF_000635855.1_De_novo_assembly_of_single-cell_genomes_protein.faa

Prochlorococcus_sp._scB245a_521N3,ftp://ftp.ncbi.nlm.nih.gov/genomes/all/GCF/000/634/575/GCF_000634575.1_De_novo_assembly_of_single-cell_genomes/GCF_000634575.1_De_novo_assembly_of_single-cell_genomes_protein.faa

Prochlorococcus_sp._scB245a_521N5,ftp://ftp.ncbi.nlm.nih.gov/genomes/all/GCF/000/635/875/GCF_000635875.1_De_novo_assembly_of_single-cell_genomes/GCF_000635875.1_De_novo_assembly_of_single-cell_genomes_protein.faa

Prochlorococcus_sp._scB245a_521O20,ftp://ftp.ncbi.nlm.nih.gov/genomes/all/GCF/000/635/255/GCF_000635255.1_De_novo_assembly_of_single-cell_genomes/GCF_000635255.1_De_novo_assembly_of_single-cell_genomes_protein.faa

Prochlorococcus_sp._scB245a_521O23,ftp://ftp.ncbi.nlm.nih.gov/genomes/all/GCF/000/634/595/GCF_000634595.1_De_novo_assembly_of_single-cell_genomes/GCF_000634595.1_De_novo_assembly_of_single-cell_genomes_protein.faa

Prochloron_didemni_P2-Fiji,ftp://ftp.ncbi.nlm.nih.gov/genomes/all/GCF/000/252/425/GCF_000252425.1_ASM25242v1/GCF_000252425.1_ASM25242v1_protein.faa

Prochloron_didemni_P3-Solomon,ftp://ftp.ncbi.nlm.nih.gov/genomes/all/GCF/000/252/465/GCF_000252465.1_ASM25246v1/GCF_000252465.1_ASM25246v1_protein.faa

Prochloron_didemni_P4-Papua_New_Guinea,ftp://ftp.ncbi.nlm.nih.gov/genomes/all/GCF/000/252/485/GCF_000252485.1_ASM25248v1/GCF_000252485.1_ASM25248v1_protein.faa

Prochlorothrix_hollandica_PCC_9006_=_CALU_1027,ftp://ftp.ncbi.nlm.nih.gov/genomes/all/GCF/000/341/585/GCF_000341585.2_ASM34158v2/GCF_000341585.2_ASM34158v2_protein.faa

Pseudanabaena_biceps_PCC_7429,ftp://ftp.ncbi.nlm.nih.gov/genomes/all/GCF/000/332/215/GCF_000332215.1_ASM33221v1/GCF_000332215.1_ASM33221v1_translated_cds.faa

Pseudanabaena_sp._'Roaring_Creek',ftp://ftp.ncbi.nlm.nih.gov/genomes/all/GCF/001/402/795/GCF_001402795.1_ASM140279v1/GCF_001402795.1_ASM140279v1_translated_cds.faa

Pseudanabaena_sp._PCC_6802,ftp://ftp.ncbi.nlm.nih.gov/genomes/all/GCF/000/332/175/GCF_000332175.1_ASM33217v1/GCF_000332175.1_ASM33217v1_translated_cds.faa

Pseudanabaena_sp._PCC_7367,ftp://ftp.ncbi.nlm.nih.gov/genomes/all/GCF/000/317/065/GCF_000317065.1_ASM31706v1/GCF_000317065.1_ASM31706v1_translated_cds.faa

Pseudanabaena_sp._SR411,ftp://ftp.ncbi.nlm.nih.gov/genomes/all/GCF/002/251/945/GCF_002251945.1_ASM225194v1/GCF_002251945.1_ASM225194v1_translated_cds.faa

Raphidiopsis_brookii_D9,ftp://ftp.ncbi.nlm.nih.gov/genomes/all/GCF/000/175/855/GCF_000175855.1_ASM17585v1/GCF_000175855.1_ASM17585v1_translated_cds.faa

Raphidiopsis_curvata_NIES-932,ftp://ftp.ncbi.nlm.nih.gov/genomes/all/GCF/002/368/135/GCF_002368135.1_ASM236813v1/GCF_002368135.1_ASM236813v1_translated_cds.faa

Richelia_intracellularis,ftp://ftp.ncbi.nlm.nih.gov/genomes/all/GCF/000/613/065/GCF_000613065.1_RintRC_1/GCF_000613065.1_RintRC_1_protein.faa

Richelia_intracellularis_HH01,ftp://ftp.ncbi.nlm.nih.gov/genomes/all/GCF/000/350/105/GCF_000350105.1_ASM35010v1/GCF_000350105.1_ASM35010v1_translated_cds.faa

Richelia_intracellularis_HM01,ftp://ftp.ncbi.nlm.nih.gov/genomes/all/GCF/000/350/125/GCF_000350125.1_ASM35012v1/GCF_000350125.1_ASM35012v1_translated_cds.faa

Rivularia_sp._PCC_7116,ftp://ftp.ncbi.nlm.nih.gov/genomes/all/GCF/000/316/665/GCF_000316665.1_ASM31666v1/GCF_000316665.1_ASM31666v1_translated_cds.faa

Roseofilum_reptotaenium_AO1-A,ftp://ftp.ncbi.nlm.nih.gov/genomes/all/GCA/001/890/975/GCA_001890975.1_ASM189097v1/GCA_001890975.1_ASM189097v1_translated_cds.faa

Rubidibacter_lacunae_KORDI_51-2,ftp://ftp.ncbi.nlm.nih.gov/genomes/all/GCF/000/473/895/GCF_000473895.1_KS51_v1/GCF_000473895.1_KS51_v1_translated_cds.faa

Scytonema_hofmannii_PCC_7110,ftp://ftp.ncbi.nlm.nih.gov/genomes/all/GCF/000/346/485/GCF_000346485.2_ASM34648v2/GCF_000346485.2_ASM34648v2_translated_cds.faa

Scytonema_millei_VB511283,ftp://ftp.ncbi.nlm.nih.gov/genomes/all/GCF/000/817/735/GCF_000817735.1_ASM81773v1/GCF_000817735.1_ASM81773v1_protein.faa

Scytonema_sp._HK-05,ftp://ftp.ncbi.nlm.nih.gov/genomes/all/GCF/001/904/675/GCF_001904675.1_ASM190467v1/GCF_001904675.1_ASM190467v1_translated_cds.faa

Scytonema_sp._NIES-4073,ftp://ftp.ncbi.nlm.nih.gov/genomes/all/GCF/002/368/435/GCF_002368435.1_ASM236843v1/GCF_002368435.1_ASM236843v1_translated_cds.faa

Scytonema_tolypothrichoides_VB-61278,ftp://ftp.ncbi.nlm.nih.gov/genomes/all/GCF/000/828/085/GCF_000828085.2_ASM82808v2/GCF_000828085.2_ASM82808v2_translated_cds.faa

Sphaerospermopsis_kisseleviana_NIES-73,ftp://ftp.ncbi.nlm.nih.gov/genomes/all/GCF/002/368/075/GCF_002368075.1_ASM236807v1/GCF_002368075.1_ASM236807v1_translated_cds.faa

Spirulina_major_PCC_6313,ftp://ftp.ncbi.nlm.nih.gov/genomes/all/GCF/001/890/765/GCF_001890765.1_ASM189076v1/GCF_001890765.1_ASM189076v1_translated_cds.faa

Spirulina_subsalsa_PCC_9445,ftp://ftp.ncbi.nlm.nih.gov/genomes/all/GCF/000/314/005/GCF_000314005.1_ASM31400v1/GCF_000314005.1_ASM31400v1_translated_cds.faa

Stanieria_cyanosphaera_PCC_7437,ftp://ftp.ncbi.nlm.nih.gov/genomes/all/GCF/000/317/575/GCF_000317575.1_ASM31757v1/GCF_000317575.1_ASM31757v1_translated_cds.faa

Stanieria_sp._NIES-3757,ftp://ftp.ncbi.nlm.nih.gov/genomes/all/GCF/002/355/455/GCF_002355455.1_ASM235545v1/GCF_002355455.1_ASM235545v1_translated_cds.faa

Synechococcus_elongatus_PCC_6301,ftp://ftp.ncbi.nlm.nih.gov/genomes/all/GCF/000/010/065/GCF_000010065.1_ASM1006v1/GCF_000010065.1_ASM1006v1_translated_cds.faa

Synechococcus_elongatus_PCC_7942,ftp://ftp.ncbi.nlm.nih.gov/genomes/all/GCF/000/012/525/GCF_000012525.1_ASM1252v1/GCF_000012525.1_ASM1252v1_translated_cds.faa

Synechococcus_lacustris_str._Tous,ftp://ftp.ncbi.nlm.nih.gov/genomes/all/GCA/002/006/815/GCA_002006815.1_ASM200681v1/GCA_002006815.1_ASM200681v1_translated_cds.faa

Synechococcus_lividus_PCC_6715,ftp://ftp.ncbi.nlm.nih.gov/genomes/all/GCF/002/754/935/GCF_002754935.1_ASM275493v1/GCF_002754935.1_ASM275493v1_translated_cds.faa

Synechococcus_sp._1G10,ftp://ftp.ncbi.nlm.nih.gov/genomes/all/GCF/002/252/625/GCF_002252625.1_ASM225262v1/GCF_002252625.1_ASM225262v1_translated_cds.faa

Synechococcus_sp._60AY4M2,ftp://ftp.ncbi.nlm.nih.gov/genomes/all/GCF/002/760/375/GCF_002760375.1_ASM276037v1/GCF_002760375.1_ASM276037v1_translated_cds.faa

Synechococcus_sp._63AY4M1,ftp://ftp.ncbi.nlm.nih.gov/genomes/all/GCF/002/760/395/GCF_002760395.1_ASM276039v1/GCF_002760395.1_ASM276039v1_translated_cds.faa

Synechococcus_sp._63AY4M2,ftp://ftp.ncbi.nlm.nih.gov/genomes/all/GCF/002/760/475/GCF_002760475.1_ASM276047v1/GCF_002760475.1_ASM276047v1_translated_cds.faa

Synechococcus_sp._65AY640,ftp://ftp.ncbi.nlm.nih.gov/genomes/all/GCF/002/760/445/GCF_002760445.1_ASM276044v1/GCF_002760445.1_ASM276044v1_translated_cds.faa

Synechococcus_sp._65AY6A5,ftp://ftp.ncbi.nlm.nih.gov/genomes/all/GCF/002/760/415/GCF_002760415.1_ASM276041v1/GCF_002760415.1_ASM276041v1_translated_cds.faa

Synechococcus_sp._65AY6Li,ftp://ftp.ncbi.nlm.nih.gov/genomes/all/GCF/002/760/345/GCF_002760345.1_ASM276034v1/GCF_002760345.1_ASM276034v1_translated_cds.faa

Synechococcus_sp._7002,ftp://ftp.ncbi.nlm.nih.gov/genomes/all/GCF/900/177/825/GCF_900177825.1_IMG-taxon_2708742468_annotated_assembly/GCF_900177825.1_IMG-taxon_2708742468_annotated_assembly_translated_cds.faa

Synechococcus_sp._8F6,ftp://ftp.ncbi.nlm.nih.gov/genomes/all/GCF/002/252/665/GCF_002252665.1_ASM225266v1/GCF_002252665.1_ASM225266v1_translated_cds.faa

Synechococcus_sp._ARS1019,ftp://ftp.ncbi.nlm.nih.gov/genomes/all/GCA/002/690/325/GCA_002690325.1_ASM269032v1/GCA_002690325.1_ASM269032v1_translated_cds.faa

Synechococcus_sp._BDU_130192,ftp://ftp.ncbi.nlm.nih.gov/genomes/all/GCF/002/721/235/GCF_002721235.1_ASM272123v1/GCF_002721235.1_ASM272123v1_translated_cds.faa

Synechococcus_sp._BL107,ftp://ftp.ncbi.nlm.nih.gov/genomes/all/GCF/000/153/805/GCF_000153805.1_ASM15380v1/GCF_000153805.1_ASM15380v1_translated_cds.faa

Synechococcus_sp._BO_8801,ftp://ftp.ncbi.nlm.nih.gov/genomes/all/GCF/002/252/675/GCF_002252675.1_ASM225267v1/GCF_002252675.1_ASM225267v1_translated_cds.faa

Synechococcus_sp._Baikal-G1,ftp://ftp.ncbi.nlm.nih.gov/genomes/all/GCA/002/737/305/GCA_002737305.1_ASM273730v1/GCA_002737305.1_ASM273730v1_translated_cds.faa

Synechococcus_sp._CB0101,ftp://ftp.ncbi.nlm.nih.gov/genomes/all/GCF/000/179/235/GCF_000179235.1_ASM17923v1/GCF_000179235.1_ASM17923v1_translated_cds.faa

Synechococcus_sp._CB0205,ftp://ftp.ncbi.nlm.nih.gov/genomes/all/GCF/000/179/255/GCF_000179255.1_ASM17925v1/GCF_000179255.1_ASM17925v1_translated_cds.faa

Synechococcus_sp._CC9311,ftp://ftp.ncbi.nlm.nih.gov/genomes/all/GCF/000/014/585/GCF_000014585.1_ASM1458v1/GCF_000014585.1_ASM1458v1_translated_cds.faa

Synechococcus_sp._CC9605,ftp://ftp.ncbi.nlm.nih.gov/genomes/all/GCF/000/012/625/GCF_000012625.1_ASM1262v1/GCF_000012625.1_ASM1262v1_translated_cds.faa

Synechococcus_sp._CC9616,ftp://ftp.ncbi.nlm.nih.gov/genomes/all/GCF/000/515/235/GCF_000515235.1_ASM51523v1/GCF_000515235.1_ASM51523v1_translated_cds.faa

Synechococcus_sp._CC9902,ftp://ftp.ncbi.nlm.nih.gov/genomes/all/GCF/000/012/505/GCF_000012505.1_ASM1250v1/GCF_000012505.1_ASM1250v1_translated_cds.faa

Synechococcus_sp._CPC100,ftp://ftp.ncbi.nlm.nih.gov/genomes/all/GCA/002/685/515/GCA_002685515.1_ASM268551v1/GCA_002685515.1_ASM268551v1_translated_cds.faa

Synechococcus_sp._CPC35,ftp://ftp.ncbi.nlm.nih.gov/genomes/all/GCA/002/684/175/GCA_002684175.1_ASM268417v1/GCA_002684175.1_ASM268417v1_translated_cds.faa

Synechococcus_sp._EAC657,ftp://ftp.ncbi.nlm.nih.gov/genomes/all/GCA/002/693/285/GCA_002693285.1_ASM269328v1/GCA_002693285.1_ASM269328v1_translated_cds.faa

Synechococcus_sp._GFB01,ftp://ftp.ncbi.nlm.nih.gov/genomes/all/GCF/001/039/265/GCF_001039265.1_ASM103926v1/GCF_001039265.1_ASM103926v1_translated_cds.faa

Synechococcus_sp._JA-2-3B'a(2-13),ftp://ftp.ncbi.nlm.nih.gov/genomes/all/GCF/000/013/225/GCF_000013225.1_ASM1322v1/GCF_000013225.1_ASM1322v1_translated_cds.faa

Synechococcus_sp._JA-3-3Ab,ftp://ftp.ncbi.nlm.nih.gov/genomes/all/GCF/000/013/205/GCF_000013205.1_ASM1320v1/GCF_000013205.1_ASM1320v1_translated_cds.faa

Synechococcus_sp._KORDI-100,ftp://ftp.ncbi.nlm.nih.gov/genomes/all/GCF/000/737/535/GCF_000737535.1_ASM73753v1/GCF_000737535.1_ASM73753v1_translated_cds.faa

Synechococcus_sp._KORDI-49,ftp://ftp.ncbi.nlm.nih.gov/genomes/all/GCF/000/737/575/GCF_000737575.1_ASM73757v1/GCF_000737575.1_ASM73757v1_translated_cds.faa

Synechococcus_sp._KORDI-52,ftp://ftp.ncbi.nlm.nih.gov/genomes/all/GCF/000/737/595/GCF_000737595.1_ASM73759v1/GCF_000737595.1_ASM73759v1_translated_cds.faa

Synechococcus_sp._Lanier,ftp://ftp.ncbi.nlm.nih.gov/genomes/all/GCA/002/151/525/GCA_002151525.1_ASM215152v1/GCA_002151525.1_ASM215152v1_translated_cds.faa

Synechococcus_sp._MED650,ftp://ftp.ncbi.nlm.nih.gov/genomes/all/GCA/002/691/345/GCA_002691345.1_ASM269134v1/GCA_002691345.1_ASM269134v1_translated_cds.faa

Synechococcus_sp._MED850,ftp://ftp.ncbi.nlm.nih.gov/genomes/all/GCA/002/700/765/GCA_002700765.1_ASM270076v1/GCA_002700765.1_ASM270076v1_translated_cds.faa

Synechococcus_sp._MIT_S9504,ftp://ftp.ncbi.nlm.nih.gov/genomes/all/GCF/001/632/105/GCF_001632105.1_ASM163210v1/GCF_001632105.1_ASM163210v1_translated_cds.faa

Synechococcus_sp._MIT_S9508,ftp://ftp.ncbi.nlm.nih.gov/genomes/all/GCF/001/632/165/GCF_001632165.1_ASM163216v1/GCF_001632165.1_ASM163216v1_translated_cds.faa

Synechococcus_sp._MIT_S9509,ftp://ftp.ncbi.nlm.nih.gov/genomes/all/GCF/001/631/935/GCF_001631935.1_ASM163193v1/GCF_001631935.1_ASM163193v1_translated_cds.faa

Synechococcus_sp._MW101C3,ftp://ftp.ncbi.nlm.nih.gov/genomes/all/GCF/002/252/635/GCF_002252635.1_ASM225263v1/GCF_002252635.1_ASM225263v1_translated_cds.faa

Synechococcus_sp._NAT40,ftp://ftp.ncbi.nlm.nih.gov/genomes/all/GCA/002/698/505/GCA_002698505.1_ASM269850v1/GCA_002698505.1_ASM269850v1_translated_cds.faa

Synechococcus_sp._NIES-970,ftp://ftp.ncbi.nlm.nih.gov/genomes/all/GCF/002/356/215/GCF_002356215.1_ASM235621v1/GCF_002356215.1_ASM235621v1_translated_cds.faa

Synechococcus_sp._NKBG042902,ftp://ftp.ncbi.nlm.nih.gov/genomes/all/GCF/000/715/475/GCF_000715475.1_ASM71547v1/GCF_000715475.1_ASM71547v1_translated_cds.faa

Synechococcus_sp._NKBG15041c,ftp://ftp.ncbi.nlm.nih.gov/genomes/all/GCF/000/485/815/GCF_000485815.1_ASM48581v1/GCF_000485815.1_ASM48581v1_translated_cds.faa

Synechococcus_sp._NP17,ftp://ftp.ncbi.nlm.nih.gov/genomes/all/GCA/002/729/835/GCA_002729835.1_ASM272983v1/GCA_002729835.1_ASM272983v1_translated_cds.faa

Synechococcus_sp._OG1,ftp://ftp.ncbi.nlm.nih.gov/genomes/all/GCF/900/177/365/GCF_900177365.1_IMG-taxon_2708742537_annotated_assembly/GCF_900177365.1_IMG-taxon_2708742537_annotated_assembly_translated_cds.faa

Synechococcus_sp._PCC_6312,ftp://ftp.ncbi.nlm.nih.gov/genomes/all/GCF/000/316/685/GCF_000316685.1_ASM31668v1/GCF_000316685.1_ASM31668v1_translated_cds.faa

Synechococcus_sp._PCC_7002,ftp://ftp.ncbi.nlm.nih.gov/genomes/all/GCF/000/019/485/GCF_000019485.1_ASM1948v1/GCF_000019485.1_ASM1948v1_translated_cds.faa

Synechococcus_sp._PCC_7003,ftp://ftp.ncbi.nlm.nih.gov/genomes/all/GCF/001/693/255/GCF_001693255.1_ASM169325v1/GCF_001693255.1_ASM169325v1_translated_cds.faa

Synechococcus_sp._PCC_7117,ftp://ftp.ncbi.nlm.nih.gov/genomes/all/GCF/001/693/275/GCF_001693275.1_ASM169327v1/GCF_001693275.1_ASM169327v1_translated_cds.faa

Synechococcus_sp._PCC_73109,ftp://ftp.ncbi.nlm.nih.gov/genomes/all/GCF/001/521/855/GCF_001521855.1_ASM152185v1/GCF_001521855.1_ASM152185v1_translated_cds.faa

Synechococcus_sp._PCC_7335,ftp://ftp.ncbi.nlm.nih.gov/genomes/all/GCF/000/155/595/GCF_000155595.1_ASM15559v1/GCF_000155595.1_ASM15559v1_translated_cds.faa

Synechococcus_sp._PCC_7336,ftp://ftp.ncbi.nlm.nih.gov/genomes/all/GCF/000/332/275/GCF_000332275.1_ASM33227v1/GCF_000332275.1_ASM33227v1_translated_cds.faa

Synechococcus_sp._PCC_7502,ftp://ftp.ncbi.nlm.nih.gov/genomes/all/GCF/000/317/085/GCF_000317085.1_ASM31708v1/GCF_000317085.1_ASM31708v1_translated_cds.faa

Synechococcus_sp._PCC_8807,ftp://ftp.ncbi.nlm.nih.gov/genomes/all/GCF/001/693/295/GCF_001693295.1_ASM169329v1/GCF_001693295.1_ASM169329v1_translated_cds.faa

Synechococcus_sp._RCC307,ftp://ftp.ncbi.nlm.nih.gov/genomes/all/GCF/000/063/525/GCF_000063525.1_ASM6352v1/GCF_000063525.1_ASM6352v1_translated_cds.faa

Synechococcus_sp._RS344,ftp://ftp.ncbi.nlm.nih.gov/genomes/all/GCA/002/724/845/GCA_002724845.1_ASM272484v1/GCA_002724845.1_ASM272484v1_translated_cds.faa

Synechococcus_sp._RS9916,ftp://ftp.ncbi.nlm.nih.gov/genomes/all/GCF/000/153/825/GCF_000153825.1_ASM15382v1/GCF_000153825.1_ASM15382v1_translated_cds.faa

Synechococcus_sp._RS9917,ftp://ftp.ncbi.nlm.nih.gov/genomes/all/GCF/000/153/065/GCF_000153065.1_ASM15306v1/GCF_000153065.1_ASM15306v1_translated_cds.faa

Synechococcus_sp._SAT82,ftp://ftp.ncbi.nlm.nih.gov/genomes/all/GCA/002/706/715/GCA_002706715.1_ASM270671v1/GCA_002706715.1_ASM270671v1_translated_cds.faa

Synechococcus_sp._SynAce01,ftp://ftp.ncbi.nlm.nih.gov/genomes/all/GCF/001/885/215/GCF_001885215.1_ASM188521v1/GCF_001885215.1_ASM188521v1_translated_cds.faa

Synechococcus_sp._TMED155,ftp://ftp.ncbi.nlm.nih.gov/genomes/all/GCA/002/171/075/GCA_002171075.2_ASM217107v2/GCA_002171075.2_ASM217107v2_translated_cds.faa

Synechococcus_sp._TMED169,ftp://ftp.ncbi.nlm.nih.gov/genomes/all/GCA/002/172/175/GCA_002172175.1_ASM217217v1/GCA_002172175.1_ASM217217v1_translated_cds.faa

Synechococcus_sp._TMED185,ftp://ftp.ncbi.nlm.nih.gov/genomes/all/GCA/002/172/045/GCA_002172045.1_ASM217204v1/GCA_002172045.1_ASM217204v1_translated_cds.faa

Synechococcus_sp._TMED187,ftp://ftp.ncbi.nlm.nih.gov/genomes/all/GCA/002/171/995/GCA_002171995.1_ASM217199v1/GCA_002171995.1_ASM217199v1_translated_cds.faa

Synechococcus_sp._TMED19,ftp://ftp.ncbi.nlm.nih.gov/genomes/all/GCA/002/168/155/GCA_002168155.1_ASM216815v1/GCA_002168155.1_ASM216815v1_translated_cds.faa

Synechococcus_sp._TMED20,ftp://ftp.ncbi.nlm.nih.gov/genomes/all/GCA/002/167/425/GCA_002167425.2_ASM216742v2/GCA_002167425.2_ASM216742v2_translated_cds.faa

Synechococcus_sp._TMED205,ftp://ftp.ncbi.nlm.nih.gov/genomes/all/GCA/002/170/705/GCA_002170705.1_ASM217070v1/GCA_002170705.1_ASM217070v1_translated_cds.faa

Synechococcus_sp._TMED66,ftp://ftp.ncbi.nlm.nih.gov/genomes/all/GCA/002/171/685/GCA_002171685.2_ASM217168v2/GCA_002171685.2_ASM217168v2_translated_cds.faa

Synechococcus_sp._TMED90,ftp://ftp.ncbi.nlm.nih.gov/genomes/all/GCA/002/172/935/GCA_002172935.1_ASM217293v1/GCA_002172935.1_ASM217293v1_translated_cds.faa

Synechococcus_sp._UTEX_2973,ftp://ftp.ncbi.nlm.nih.gov/genomes/all/GCF/000/817/325/GCF_000817325.1_ASM81732v1/GCF_000817325.1_ASM81732v1_translated_cds.faa

Synechococcus_sp._WH_5701,ftp://ftp.ncbi.nlm.nih.gov/genomes/all/GCF/000/153/045/GCF_000153045.1_ASM15304v1/GCF_000153045.1_ASM15304v1_translated_cds.faa

Synechococcus_sp._WH_7803,ftp://ftp.ncbi.nlm.nih.gov/genomes/all/GCF/000/063/505/GCF_000063505.1_ASM6350v1/GCF_000063505.1_ASM6350v1_translated_cds.faa

Synechococcus_sp._WH_7805,ftp://ftp.ncbi.nlm.nih.gov/genomes/all/GCF/000/153/285/GCF_000153285.1_ASM15328v1/GCF_000153285.1_ASM15328v1_translated_cds.faa

Synechococcus_sp._WH_8016,ftp://ftp.ncbi.nlm.nih.gov/genomes/all/GCF/000/230/675/GCF_000230675.1_ASM23067v1/GCF_000230675.1_ASM23067v1_translated_cds.faa

Synechococcus_sp._WH_8020,ftp://ftp.ncbi.nlm.nih.gov/genomes/all/GCF/001/040/845/GCF_001040845.1_ASM104084v1/GCF_001040845.1_ASM104084v1_translated_cds.faa

Synechococcus_sp._WH_8102,ftp://ftp.ncbi.nlm.nih.gov/genomes/all/GCF/000/195/975/GCF_000195975.1_ASM19597v1/GCF_000195975.1_ASM19597v1_translated_cds.faa

Synechococcus_sp._WH_8103,ftp://ftp.ncbi.nlm.nih.gov/genomes/all/GCF/001/182/765/GCF_001182765.1_WH8103.1/GCF_001182765.1_WH8103.1_translated_cds.faa

Synechococcus_sp._WH_8109,ftp://ftp.ncbi.nlm.nih.gov/genomes/all/GCF/000/161/795/GCF_000161795.2_ASM16179v2/GCF_000161795.2_ASM16179v2_translated_cds.faa

Synechocystis_sp._PCC_6714,ftp://ftp.ncbi.nlm.nih.gov/genomes/all/GCF/000/478/825/GCF_000478825.2_ASM47882v2/GCF_000478825.2_ASM47882v2_translated_cds.faa

Synechocystis_sp._PCC_6803,ftp://ftp.ncbi.nlm.nih.gov/genomes/all/GCF/000/270/265/GCF_000270265.1_ASM27026v1/GCF_000270265.1_ASM27026v1_translated_cds.faa

Synechocystis_sp._PCC_6803_substr._GT-I,ftp://ftp.ncbi.nlm.nih.gov/genomes/all/GCF/000/284/135/GCF_000284135.1_ASM28413v1/GCF_000284135.1_ASM28413v1_translated_cds.faa

Synechocystis_sp._PCC_6803_substr._PCC-N,ftp://ftp.ncbi.nlm.nih.gov/genomes/all/GCF/000/284/215/GCF_000284215.1_ASM28421v1/GCF_000284215.1_ASM28421v1_translated_cds.faa

Synechocystis_sp._PCC_6803_substr._PCC-P,ftp://ftp.ncbi.nlm.nih.gov/genomes/all/GCF/000/284/455/GCF_000284455.1_ASM28445v1/GCF_000284455.1_ASM28445v1_translated_cds.faa

Synechocystis_sp._PCC_7509,ftp://ftp.ncbi.nlm.nih.gov/genomes/all/GCF/000/332/075/GCF_000332075.2_ASM33207v2/GCF_000332075.2_ASM33207v2_translated_cds.faa

Thermosynechococcus_elongatus_BP-1,ftp://ftp.ncbi.nlm.nih.gov/genomes/all/GCF/000/011/345/GCF_000011345.1_ASM1134v1/GCF_000011345.1_ASM1134v1_translated_cds.faa

Thermosynechococcus_sp._NK55a,ftp://ftp.ncbi.nlm.nih.gov/genomes/all/GCF/000/505/665/GCF_000505665.1_ASM50566v1/GCF_000505665.1_ASM50566v1_translated_cds.faa

Thermosynechococcus_vulcanus_NIES-2134,ftp://ftp.ncbi.nlm.nih.gov/genomes/all/GCF/003/990/665/GCF_003990665.1_ASM399066v2/GCF_003990665.1_ASM399066v2_translated_cds.faa

Tolypothrix_bouteillei_VB521301,ftp://ftp.ncbi.nlm.nih.gov/genomes/all/GCF/000/760/695/GCF_000760695.2_ASM76069v2/GCF_000760695.2_ASM76069v2_translated_cds.faa

Tolypothrix_campylonemoides_VB511288,ftp://ftp.ncbi.nlm.nih.gov/genomes/all/GCF/000/828/075/GCF_000828075.2_ASM82807v2/GCF_000828075.2_ASM82807v2_translated_cds.faa

Tolypothrix_sp._NIES-4075,ftp://ftp.ncbi.nlm.nih.gov/genomes/all/GCF/002/218/085/GCF_002218085.1_ASM221808v1/GCF_002218085.1_ASM221808v1_translated_cds.faa

Tolypothrix_sp._PCC_7601,ftp://ftp.ncbi.nlm.nih.gov/genomes/all/GCF/000/300/115/GCF_000300115.1_Fremyella_diplosiphon-2.0.1/GCF_000300115.1_Fremyella_diplosiphon-2.0.1_translated_cds.faa

Tolypothrix_tenuis_PCC_7101,ftp://ftp.ncbi.nlm.nih.gov/genomes/all/GCF/002/368/295/GCF_002368295.1_ASM236829v1/GCF_002368295.1_ASM236829v1_translated_cds.faa

Trichodesmium_erythraeum_21-75,ftp://ftp.ncbi.nlm.nih.gov/genomes/all/GCF/000/963/755/GCF_000963755.1_ASM96375v2/GCF_000963755.1_ASM96375v2_protein.faa

Trichodesmium_erythraeum_IMS101,ftp://ftp.ncbi.nlm.nih.gov/genomes/all/GCF/000/014/265/GCF_000014265.1_ASM1426v1/GCF_000014265.1_ASM1426v1_translated_cds.faa

Trichodesmium_thiebautii_H9-4,ftp://ftp.ncbi.nlm.nih.gov/genomes/all/GCF/000/987/385/GCF_000987385.1_H9-4/GCF_000987385.1_H9-4_protein.faa

Trichormus_sp._NMC-1,ftp://ftp.ncbi.nlm.nih.gov/genomes/all/GCF/001/858/025/GCF_001858025.1_ASM185802v1/GCF_001858025.1_ASM185802v1_translated_cds.faa

Trichormus_variabilis_ATCC_29413,ftp://ftp.ncbi.nlm.nih.gov/genomes/all/GCF/000/204/075/GCF_000204075.1_ASM20407v1/GCF_000204075.1_ASM20407v1_translated_cds.faa

Trichormus_variabilis_NIES-23,ftp://ftp.ncbi.nlm.nih.gov/genomes/all/GCF/002/368/015/GCF_002368015.1_ASM236801v1/GCF_002368015.1_ASM236801v1_translated_cds.faa

Tychonema_bourrellyi_FEM_GT703,ftp://ftp.ncbi.nlm.nih.gov/genomes/all/GCF/002/412/335/GCF_002412335.2_ASM241233v2/GCF_002412335.2_ASM241233v2_translated_cds.faa

Vulcanococcus_limneticus_LL,ftp://ftp.ncbi.nlm.nih.gov/genomes/all/GCF/002/252/705/GCF_002252705.1_ASM225270v1/GCF_002252705.1_ASM225270v1_translated_cds.faa

Xenococcus_sp._PCC_7305,ftp://ftp.ncbi.nlm.nih.gov/genomes/all/GCF/000/332/055/GCF_000332055.1_ASM33205v1/GCF_000332055.1_ASM33205v1_translated_cds.faa

cyanobacterium_BACL30_MAG-120619-bin27,ftp://ftp.ncbi.nlm.nih.gov/genomes/all/GCA/001/438/415/GCA_001438415.1_ASM143841v1/GCA_001438415.1_ASM143841v1_translated_cds.faa

cyanobacterium_PCC_7702,ftp://ftp.ncbi.nlm.nih.gov/genomes/all/GCF/000/332/255/GCF_000332255.1_ASM33225v1/GCF_000332255.1_ASM33225v1_translated_cds.faa

cyanobacterium_TDX16,ftp://ftp.ncbi.nlm.nih.gov/genomes/all/GCA/002/213/405/GCA_002213405.1_ASM221340v1/GCA_002213405.1_ASM221340v1_translated_cds.faa

cyanobacterium_endosymbiont_of_Epithemia_turgida_isolate_EtSB_Lake_Yunoko,ftp://ftp.ncbi.nlm.nih.gov/genomes/all/GCF/000/829/235/GCF_000829235.1_ASM82923v1/GCF_000829235.1_ASM82923v1_translated_cds.faa

cyanobacterium_endosymbiont_of_Rhopalodia_gibberula,ftp://ftp.ncbi.nlm.nih.gov/genomes/all/GCF/003/574/135/GCF_003574135.1_ASM357413v1/GCF_003574135.1_ASM357413v1_translated_cds.faa

filamentous_cyanobacterium_ESFC-1,ftp://ftp.ncbi.nlm.nih.gov/genomes/all/GCF/000/380/225/GCF_000380225.1_ASM38022v1/GCF_000380225.1_ASM38022v1_translated_cds.faa

unicellular_cyanobacterium_SU2,ftp://ftp.ncbi.nlm.nih.gov/genomes/all/GCF/002/110/465/GCF_002110465.1_croco123/GCF_002110465.1_croco123_translated_cds.faa

unicellular_cyanobacterium_SU3,ftp://ftp.ncbi.nlm.nih.gov/genomes/all/GCF/002/172/675/GCF_002172675.1_croco123Su3/GCF_002172675.1_croco123Su3_translated_cds.faa

[Leptolyngbya]_sp._JSC-1,ftp://ftp.ncbi.nlm.nih.gov/genomes/all/GCF/000/733/415/GCF_000733415.1_ASM73341v1/GCF_000733415.1_ASM73341v1_translated_cds.faa

[Oscillatoria]_sp._PCC_6506,ftp://ftp.ncbi.nlm.nih.gov/genomes/all/GCF/000/180/455/GCF_000180455.1_ASM18045v1/GCF_000180455.1_ASM18045v1_translated_cds.faa

[Scytonema_hofmanni]_UTEX_2349,ftp://ftp.ncbi.nlm.nih.gov/genomes/all/GCF/000/582/685/GCF_000582685.1_ASM58268v1/GCF_000582685.1_ASM58268v1_translated_cds.faa

**Supplementary Data 3: Final HMM profile generated at the end of the iterative search for homologs of calcyanin in cyanobacterial genomes.**

```
HMMER3/f [3.3 | Nov 2019]
NAME  alignment_0
LENG  226
ALPH  amino
RF    no
MM    no
CONS  yes
CS    no
MAP   yes
DATE  Tue Jul 21 14:42:24 2020
NSEQ  32
EFFN  1.535156
CKSUM 4009382736
STATS LOCAL MSV      -11.2853  0.70396
STATS LOCAL VITERBI  -11.9627  0.70396
STATS LOCAL FORWARD   -5.2243  0.70396
HMM          A        C        D        E        F        G        H        I        K        L        M        N        P        Q        R        S        T        V        W        Y
            m->m     m->i     m->d     i->m     i->i     d->m     d->d
COMPO  2.39555  4.60090  3.17697  2.72838  3.67304  1.687048  3.95144  2.84231  2.79592  2.56153  3.64724  3.23445  3.43804  3.01503  3.09120  2.66370  2.78762  2.55577  4.55821  4.02755
2.68399  4.42305  2.77578  2.73121  3.46430  1.924312  3.72570  3.29403  2.67778  2.69365  4.24686  2.90393  2.73805  3.18003  2.89881  2.37906  2.77408  2.98442  4.58523  3.61583
          0.41999  1.91702  1.63014  2.99841  0.05115  0.00000        *

1  2.70919  5.15510  2.42026  1.73216  4.46595  2.208576  3.62900  3.93447  2.41754  3.44426  4.22815  2.81101  3.80428  2.65179  2.49382  2.66433  2.95214  3.53218  5.60417  4.19859  84  e  -  -  -
2.68618  4.42225  2.77519  2.73123  3.46354  1.924104  3.72494  3.29354  2.67741  2.69355  4.24690  2.90347  2.73739  3.18146  2.89801  2.37887  2.77519  2.98518  4.58477  3.61503
          0.02644  4.04194  4.76429  0.61958  0.77255  0.71057  0.67603

2  2.32449  4.95446  2.76631  1.91512  4.25733  2.69704  3.64373  3.67135  2.40499  3.24686  4.06226  2.89372  3.81952  2.16347  2.84723  2.66855  2.91319  3.12906  5.45918  4.10357  85  e  -  -  -
2.68618  4.42225  2.77519  2.73123  3.46354  1.924104  3.72494  3.29354  2.67741  2.69355  4.24690  2.90347  2.73739  3.18146  2.89801  2.37887  2.77519  2.98518  4.58477  3.61503
          0.02644  4.04194  4.76429  0.61958  0.77255  0.44049  1.03204

3  2.53965  4.20814  4.05187  2.87529  2.79047  3.06484  4.14749  1.75809  3.39016  2.29192  3.29711  3.75515  4.19333  3.64055  3.62388  3.11846  2.95356  1.77611  4.89687  3.69430  86  i  -  -  -
2.68618  4.42225  2.77519  2.73123  3.46354  1.924104  3.72494  3.29354  2.67741  2.69355  4.24690  2.90347  2.73739  3.18146  2.89801  2.37887  2.77519  2.98518  4.58477  3.61503
          0.02243  4.20435  4.92670  0.61958  0.77255  0.47062  0.97981

4  2.40273  5.08588  2.81806  2.14903  4.40826  2.330752  3.64080  3.87172  2.40974  3.39097  4.15879  2.61112  3.15084  2.24847  2.89818  2.38404  2.90583  3.46979  5.55686  4.16373  87  e  -  -  -
2.68618  4.42225  2.77519  2.73123  3.46354  1.924104  3.72494  3.29354  2.67741  2.69355  4.24690  2.90347  2.73739  3.18146  2.89801  2.37887  2.77519  2.98518  4.58477  3.61503
          0.02161  4.24114  4.96349  0.61958  0.77255  0.49930  0.93384

5  2.68276  5.13210  2.50877  2.16298  4.44661  1.929152  3.47193  3.51222  2.41056  3.42466  4.18510  2.73471  3.83869  2.48493  2.90391  2.28405  2.91746  3.50707  5.58215  4.18193  88  e  -  -  -
2.68618  4.42225  2.77519  2.73123  3.46354  1.924104  3.72494  3.29354  2.67741  2.69355  4.24690  2.90347  2.73739  3.18146  2.89801  2.37887  2.77519  2.98518  4.58477  3.61503
          0.02014  4.31090  5.03324  0.61958  0.77255  0.49930  0.93384

6  2.74242  5.08935  3.04387  2.48359  4.39785  2.811816  3.08426  3.83690  1.84925  3.36237  2.92620  2.66233  3.90420  2.10089  2.75249  2.73199  2.96561  3.46379  5.52403  4.17587  89  k  -  -  -
2.68618  4.42225  2.77519  2.73123  3.46354  1.924104  3.72494  3.29354  2.67741  2.69355  4.24690  2.90347  2.73739  3.18146  2.89801  2.37887  2.77519  2.98518  4.58477  3.61503
          0.01886  4.37610  5.09845  0.61958  0.77255  0.49930  0.93384

7  2.21050  4.43029  4.72267  4.16162  3.45677  3.440864  4.70236  1.71609  4.03989  1.33128  3.32287  4.35004  4.61820  4.23635  4.19309  3.63041  3.30469  1.80458  5.24159  4.08228  90  l  -  -  -
2.68618  4.42225  2.77519  2.73123  3.46354  1.924104  3.72494  3.29354  2.67741  2.69355  4.24690  2.90347  2.73739  3.18146  2.89801  2.37887  2.77519  2.98518  4.58477  3.61503
          0.01886  4.37610  5.09845  0.61958  0.77255  0.49930  0.93384

8  2.69997  5.13826  2.97159  2.23336  4.46174  2.41644  3.65968  3.92550  2.16595  3.42964  4.18685  2.95631  3.87028  2.31569  2.19458  2.44406  2.70520  3.51773  5.57971  3.49201  91  k  -  -  -
2.68618  4.42225  2.77519  2.73123  3.46354  1.924104  3.72494  3.29354  2.67741  2.69355  4.24690  2.90347  2.73739  3.18146  2.89801  2.37887  2.77519  2.98518  4.58477  3.61503
          0.01773  4.43731  5.15966  0.61958  0.77255  0.49930  0.93384

9  2.51447  5.12561  2.40736  2.22067  4.43668  2.77284  3.67267  3.89969  2.42999  2.97620  4.17937  2.94790  3.86947  2.13451  2.91799  2.44162  2.41190  3.49844  5.58102  4.18928  92  q  -  -  -
2.68566  4.42235  2.77512  2.73111  3.46363  1.924176  3.72490  3.29364  2.67750  2.69365  4.24699  2.90356  2.73728  3.18156  2.89810  2.37896  2.77529  2.98500  4.58487  3.61513
          0.14656  2.03576  5.15966  1.06937  0.42041  0.49930  0.93384

10  2.60927  4.33788  4.82290  4.24216  2.79203  3.387936  4.60375  1.39065  4.08833  2.13750  2.38950  4.34022  4.55401  4.24370  4.16898  3.55956  3.22681  1.70156  5.08837  3.92482  99  i  -  -  -
2.68618  4.42225  2.77519  2.73123  3.46354  1.924104  3.72494  3.29354  2.67741  2.69355  4.24690  2.90347  2.73739  3.18146  2.89801  2.37887  2.77519  2.98518  4.58477  3.61503
          0.01773  4.43731  5.15966  0.61958  0.77255  0.49930  0.93384

11  2.07295  4.49746  3.79245  3.69373  4.90192  0.512248  4.80924  4.32725  3.89849  4.05243  4.89374  3.74498  4.04444  4.13435  4.14025  2.76182  3.11335  3.66866  6.20307  5.06485  100  G  -  -  -
2.68618  4.42225  2.77519  2.73123  3.46354  1.924104  3.72494  3.29354  2.67741  2.69355  4.24690  2.90347  2.73739  3.18146  2.89801  2.37887  2.77519  2.98518  4.58477  3.61503
          0.01773  4.43731  5.15966  0.61958  0.77255  0.49930  0.93384

12  1.97581  4.67211  3.24857  2.89212  4.48500  1.062688  4.10609  3.91270  2.52453  3.53240  4.34606  3.28411  3.95429  3.29631  3.35758  2.31855  3.00464  3.45207  5.75510  4.46798  101  g  -  -  -
2.68618  4.42225  2.77519  2.73123  3.46354  1.924104  3.72494  3.29354  2.67741  2.69355  4.24690  2.90347  2.73739  3.18146  2.89801  2.37887  2.77519  2.98518  4.58477  3.61503
          0.01773  4.43731  5.15966  0.61958  0.77255  0.49930  0.93384

13  2.02879  4.58783  3.33242  2.79827  2.73188  2.06584  3.88892  3.30070  2.78721  2.96957  3.82305  2.78527  3.95295  3.11326  3.20251  2.22248  2.50558  3.00492  5.27747  4.00279  102  a  -  -  -
2.68618  4.42225  2.77519  2.73123  3.46354  1.924104  3.72494  3.29354  2.67741  2.69355  4.24690  2.90347  2.73739  3.18146  2.89801  2.37887  2.77519  2.98518  4.58477  3.61503
          0.01773  4.43731  5.15966  0.61958  0.77255  0.49930  0.93384

14  3.06516  4.47946  5.05161  4.46392  2.70910  3.537688  4.77153  1.64669  4.30169  1.47155  2.28355  4.55179  4.69073  4.38911  4.33735  3.75546  3.38814  1.75233  5.14515  4.03243  103  l  -  -  -
2.68618  4.42225  2.77519  2.73123  3.46354  1.924104  3.72494  3.29354  2.67741  2.69355  4.24690  2.90347  2.73739  3.18146  2.89801  2.37887  2.77519  2.98518  4.58477  3.61503
          0.01773  4.43731  5.15966  0.61958  0.77255  0.49930  0.93384

15  3.33972  4.61309  5.24404  4.68673  3.47154  3.767192  5.12334  1.66866  4.55858  1.44195  2.45857  4.82669  4.93330  4.66084  4.62770  4.06774  3.57337  1.28671  5.44738  4.34416  104  v  -  -  -
2.68620  4.42227  2.77522  2.73107  3.46356  1.92412  3.72497  3.29356  2.67743  2.69357  4.24692  2.90349  2.73742  3.18149  2.89803  2.37889  2.77503  2.98521  4.58479  3.61505
          0.07092  2.76905  5.15966  0.65228  0.73575  0.49930  0.93384

16  2.21040  4.43091  3.69650  3.59603  4.87756  0.548064  4.73847  4.32728  3.81957  4.02983  4.83963  3.65358  3.97827  4.04636  4.07734  2.49676  3.02729  3.62932  6.20776  5.02795  107  G  -  -  -
2.68618  4.42225  2.77519  2.73123  3.46354  1.924104  3.72494  3.29354  2.67741  2.69355  4.24690  2.90347  2.73739  3.18146  2.89801  2.37887  2.77519  2.98518  4.58477  3.61503
          0.01773  4.43731  5.15966  0.61958  0.77255  0.49930  0.93384

17  1.82319  4.39211  3.67719  3.13606  3.61486  1.679272  4.03597  2.90294  3.07393  2.12925  2.95858  3.49044  4.06252  3.37982  3.40434  2.90502  2.94509  2.67809  5.08502  3.85909  108  a  -  -  -
2.68618  4.42225  2.77519  2.73123  3.46354  1.924104  3.72494  3.29354  2.67741  2.69355  4.24690  2.90347  2.73739  3.18146  2.89801  2.37887  2.77519  2.98518  4.58477  3.61503
          0.01773  4.43731  5.15966  0.61958  0.77255  0.49930  0.93384

18  1.93870  4.64048  3.25569  2.69674  3.82528  2.852816  3.79390  3.20965  2.67499  2.87658  3.15416  2.85106  3.95431  2.62127  3.09938  2.31185  2.90857  2.63827  5.18941  3.37625  109  a  -  -  -
2.68618  4.42225  2.77519  2.73123  3.46354  1.924104  3.72494  3.29354  2.67741  2.69355  4.24690  2.90347  2.73739  3.18146  2.89801  2.37887  2.77519  2.98518  4.58477  3.61503
          0.08007  4.43731  2.73152  0.61958  0.77255  0.49930  0.93384

19  1.95278  4.28195  3.86282  3.30605  3.49017  2.40592  4.08964  2.55108  3.23213  2.32343  3.46197  3.61022  4.10772  3.50806  3.51791  2.97472  2.55927  1.70399  4.98845  3.77678  110  v  -  -  -
2.68618  4.42225  2.77519  2.73123  3.46354  1.924104  3.72494  3.29354  2.67741  2.69355  4.24690  2.90347  2.73739  3.18146  2.89801  2.37887  2.77519  2.98518  4.58477  3.61503
          0.01886  4.37610  5.09845  0.61958  0.77255  0.58923  0.80913

20  3.41139  5.10676  4.18445  4.15943  5.23579  0.202024  5.21727  4.98871  4.40665  4.56494  5.56531  4.35148  4.50952  4.69177  4.55873  3.60189  3.92983  4.41912  6.16776  5.36462  111  G  -  -  -
2.68618  4.42225  2.77519  2.73123  3.46354  1.924104  3.72494  3.29354  2.67741  2.69355  4.24690  2.90347  2.73739  3.18146  2.89801  2.37887  2.77519  2.98518  4.58477  3.61503
          0.01886  4.37610  5.09845  0.61958  0.77255  0.58923  0.80913

21  2.90195  5.45540  1.67375  1.89824  4.76715  2.742992  2.99482  4.27097  2.63533  3.74545  4.52456  2.88453  3.93939  1.99200  3.16374  2.82534  3.15413  3.83362  5.88343  4.42944  112  d  -  -  -
2.68618  4.42225  2.77519  2.73123  3.46354  1.924104  3.72494  3.29354  2.67741  2.69355  4.24690  2.90347  2.73739  3.18146  2.89801  2.37887  2.77519  2.98518  4.58477  3.61503
          0.01886  4.37610  5.09845  0.61958  0.77255  0.58923  0.80913

22  3.28063  4.55514  5.15347  4.60906  3.55386  3.738424  5.12413  1.66425  4.49385  1.81831  2.58339  4.77028  4.91573  4.64869  4.60439  4.03465  3.52061  1.03698  5.50608  4.36961  113  v  -  -  -
2.68618  4.42225  2.77519  2.73123  3.46354  1.924104  3.72494  3.29354  2.67741  2.69355  4.24690  2.90347  2.73739  3.18146  2.89801  2.37887  2.77519  2.98518  4.58477  3.61503
          0.01886  4.37610  5.09845  0.61958  0.77255  0.44510  1.02377

23  2.57133  4.39853  4.74802  4.17916  2.78405  3.400232  4.62167  2.21224  4.04270  1.53513  3.30416  4.32468  4.57461  4.21656  4.16204  3.57624  3.26994  1.28275  5.13058  3.95308  114  v  -  -  -
2.68618  4.42225  2.77519  2.73123  3.46354  1.924104  3.72494  3.29354  2.67741  2.69355  4.24690  2.90347  2.73739  3.18146  2.89801  2.37887  2.77519  2.98518  4.58477  3.61503
          0.01773  4.43731  5.15966  0.61958  0.77255  0.49930  0.93384

24  2.82137  4.86789  2.56031  2.71994  4.23282  1.01864  4.02117  3.63359  2.94610  2.31316  4.19090  3.20448  4.04014  3.23648  3.38677  2.89907  3.13403  3.31482  5.58115  4.25380  115  g  -  -  -
2.68618  4.42225  2.77519  2.73123  3.46354  1.924104  3.72494  3.29354  2.67741  2.69355  4.24690  2.90347  2.73739  3.18146  2.89801  2.37887  2.77519  2.98518  4.58477  3.61503
          0.01773  4.43731  5.15966  0.61958  0.77255  0.49930  0.93384

25  2.03193  4.39725  3.78611  3.58642  4.82046  0.737512  4.67360  4.27610  3.72902  3.94878  4.73976  3.63776  3.94808  3.95996  4.01309  1.86781  2.97516  3.58415  6.14329  4.96597  116  g  -  -  -
2.68618  4.42225  2.77519  2.73123  3.46354  1.924104  3.72494  3.29354  2.67741  2.69355  4.24690  2.90347  2.73739  3.18146  2.89801  2.37887  2.77519  2.98518  4.58477  3.61503
          0.01773  4.43731  5.15966  0.61958  0.77255  0.47739  0.96864

26  1.69927  4.28781  3.89591  3.33101  3.44938  2.139584  4.08610  2.38137  3.25287  2.48919  2.66109  3.63605  4.13302  3.52246  3.52904  3.01334  2.94921  2.18566  4.95135  3.74307  117  a  -  -  -
2.68580  4.42242  2.77537  2.73141  3.46371  1.924136  3.72495  3.29371  2.67758  2.69337  4.24707  2.90313  2.73720  3.18164  2.89818  2.37891  2.77495  2.98536  4.58494  3.61491
          0.07937  2.65065  5.16811  1.93619  0.15578  0.48576  0.95510

27  2.43927  4.17927  4.29673  3.71132  3.30175  2.512552  4.19838  1.66891  3.58137  2.13023  2.82280  3.88869  4.23693  3.79202  3.73902  3.16750  2.60854  1.97392  4.84073  3.65115  132  i  -  -  -
2.68618  4.42225  2.77519  2.73123  3.46354  1.924104  3.72494  3.29354  2.67741  2.69355  4.24690  2.90347  2.73739  3.18146  2.89801  2.37887  2.77519  2.98518  4.58477  3.61503
          0.01758  4.44576  5.16811  0.61958  0.77255  0.48576  0.95510

28  2.07922  4.50411  3.80310  3.70622  4.91317  0.505872  4.82090  4.33974  3.91217  4.06465  4.90560  3.75467  4.05188  4.14697  4.15250  2.76901  3.12102  3.67861  6.21363  5.07691  133  G  -  -  -
2.68618  4.42225  2.77519  2.73123  3.46354  1.924104  3.72494  3.29354  2.67741  2.69355  4.24690  2.90347  2.73739  3.18146  2.89801  2.37887  2.77519  2.98518  4.58477  3.61503
          0.01758  4.44576  5.16811  0.61958  0.77255  0.48576  0.95510

29  1.93300  5.07425  2.83443  2.23433  4.47564  1.369408  3.85429  3.91527  2.67660  3.48498  4.29526  3.03860  3.96503  2.58380  3.14432  2.81292  3.08311  3.53404  5.69104  4.32623  134  g  -  -  -
2.68618  4.42225  2.77519  2.73123  3.46354  1.924104  3.72494  3.29354  2.67741  2.69355  4.24690  2.90347  2.73739  3.18146  2.89801  2.37887  2.77519  2.98518  4.58477  3.61503
          0.01758  4.44576  5.16811  0.61958  0.77255  0.48576  0.95510

30  1.53545  4.35366  3.88502  3.39806  3.94563  2.715952  4.28577  3.17844  3.34498  3.00436  3.89707  3.60543  4.01815  3.62743  3.65052  2.36126  1.90468  1.94062  5.38961  4.18032  135  a  -  -  -
2.68618  4.42225  2.77519  2.73123  3.46354  1.924104  3.72494  3.29354  2.67741  2.69355  4.24690  2.90347  2.73739  3.18146  2.89801  2.37887  2.77519  2.98518  4.58477  3.61503
          0.01758  4.44576  5.16811  0.61958  0.77255  0.48576  0.95510

31  1.69645  4.30566  4.56842  3.99118  2.82379  3.28556  4.46202  1.84865  3.86083  2.02441  3.33332  4.15258  4.44855  4.05608  4.00077  3.41850  3.14918  1.74635  5.03943  3.85726  136  a  -  -  -
2.68618  4.42225  2.77519  2.73123  3.46354  1.924104  3.72494  3.29354  2.67741  2.69355  4.24690  2.90347  2.73739  3.18146  2.89801  2.37887  2.77519  2.98518  4.58477  3.61503
          0.01758  4.44576  5.16811  0.61958  0.77255  0.48576  0.95510

32  3.52943  5.20981  4.31481  4.30055  5.36069  0.17488  5.34468  5.13918  4.55586  4.70143  5.70785  4.48216  4.61282  4.83556  4.69240  3.72289  4.05210  4.55636  6.26633  5.49441  137  G  -  -  -
2.68618  4.42225  2.77519  2.73123  3.46354  1.924104  3.72494  3.29354  2.67741  2.69355  4.24690  2.90347  2.73739  3.18146  2.89801  2.37887  2.77519  2.98518  4.58477  3.61503
          0.01758  4.44576  5.16811  0.61958  0.77255  0.48576  0.95510

33  1.45979  4.44792  3.61896  3.14295  3.03399  1.649992  4.16088  3.45122  3.13399  3.14339  4.00018  3.44546  3.96839  3.43353  3.49816  2.06215  2.93000  3.09349  5.44769  4.20327  138  a  -  -  -
2.68618  4.42225  2.77519  2.73123  3.46354  1.924104  3.72494  3.29354  2.67741  2.69355  4.24690  2.90347  2.73739  3.18146  2.89801  2.37887  2.77519  2.98518  4.58477  3.61503
          0.01758  4.44576  5.16811  0.61958  0.77255  0.48576  0.95510

34  1.90848  4.11729  4.47238  3.87183  2.75987  3.091968  4.18733  2.25074  3.70163  2.12749  2.76526  3.95654  4.22494  3.86774  3.78213  3.16849  2.94501  1.83144  3.47929  3.55791  139  v  -  -  -
2.68618  4.42225  2.77519  2.73123  3.46354  1.924104  3.72494  3.29354  2.67741  2.69355  4.24690  2.90347  2.73739  3.18146  2.89801  2.37887  2.77519  2.98518  4.58477  3.61503
          0.07937  4.44576  2.73997  0.61958  0.77255  0.48576  0.95510

35  2.45260  4.19946  4.43535  3.85003  2.78880  3.163856  4.29260  2.09653  3.71158  1.78773  3.26854  4.00073  4.30807  3.90590  3.84448  3.25899  2.32568  1.73089  4.88572  3.70328  140  v  -  -  -
2.68618  4.42225  2.77519  2.73123  3.46354  1.924104  3.72494  3.29354  2.67741  2.69355  4.24690  2.90347  2.73739  3.18146  2.89801  2.37887  2.77519  2.98518  4.58477  3.61503
          0.01869  4.38508  5.10742  0.61958  0.77255  0.43274  1.04620

36  2.60789  4.45378  4.87369  4.27809  2.32882  3.437344  4.60881  2.15199  4.11831  1.27693  2.03114  4.39331  4.57961  4.21757  4.17576  3.61600  3.32090  2.44072  5.02176  3.90433  141  l  -  -  -
2.68618  4.42225  2.77519  2.73123  3.46354  1.924104  3.72494  3.29354  2.67741  2.69355  4.24690  2.90347  2.73739  3.18146  2.89801  2.37887  2.77519  2.98518  4.58477  3.61503
          0.01758  4.44576  5.16811  0.61958  0.77255  0.48576  0.95510

37  2.68893  4.56169  3.74739  3.46564  4.17656  0.657952  4.46161  3.51795  3.42573  2.62105  4.28134  3.67285  4.11372  3.79098  3.70300  2.88308  3.13672  3.05654  5.65330  4.35347  142  g  -  -  -
2.68618  4.42225  2.77519  2.73123  3.46354  1.924104  3.72494  3.29354  2.67741  2.69355  4.24690  2.90347  2.73739  3.18146  2.89801  2.37887  2.77519  2.98518  4.58477  3.61503
          0.01758  4.44576  5.16811  0.61958  0.77255  0.48576  0.95510

38  2.09934  4.44290  3.74819  3.47165  4.58945  2.583736  4.53124  3.91155  3.50238  3.65499  4.50910  3.60779  0.94579  3.80755  3.79816  2.69175  2.39829  3.39568  5.94501  4.74429  143  p  -  -  -
2.68618  4.42225  2.77519  2.73123  3.46354  1.924104  3.72494  3.29354  2.67741  2.69355  4.24690  2.90347  2.73739  3.18146  2.89801  2.37887  2.77519  2.98518  4.58477  3.61503
          0.01758  4.44576  5.16811  0.61958  0.77255  0.48576  0.95510

39  1.99608  4.88143  3.10098  2.55196  4.17475  2.03604  3.74121  3.59245  2.30827  3.19066  4.00095  3.06267  2.83625  2.89009  2.65393  2.72047  2.92737  2.85032  5.42600  4.09403  144  a  -  -  -
2.68618  4.42225  2.77519  2.73123  3.46354  1.924104  3.72494  3.29354  2.67741  2.69355  4.24690  2.90347  2.73739  3.18146  2.89801  2.37887  2.77519  2.98518  4.58477  3.61503
          0.01758  4.44576  5.16811  0.61958  0.77255  0.48576  0.95510

40  3.02915  5.02711  3.24230  3.08350  4.86169  0.562048  4.35896  4.35605  2.39689  3.94766  4.82733  3.49605  4.21153  3.58337  3.42282  3.11858  3.43093  3.89354  6.00127  4.79576  145  g  -  -  -
2.68618  4.42225  2.77519  2.73123  3.46354  1.924104  3.72494  3.29354  2.67741  2.69355  4.24690  2.90347  2.73739  3.18146  2.89801  2.37887  2.77519  2.98518  4.58477  3.61503
          0.01758  4.44576  5.16811  0.61958  0.77255  0.48576  0.95510

41  1.26293  4.39795  4.16829  3.64299  3.64192  3.0438  4.40477  2.49774  3.54665  2.48423  2.74604  3.88223  4.28908  3.82381  3.80387  3.15068  3.10853  1.86237  5.25120  4.05219  146  a  -  -  -
2.68618  4.42225  2.77519  2.73123  3.46354  1.924104  3.72494  3.29354  2.67741  2.69355  4.24690  2.90347  2.73739  3.18146  2.89801  2.37887  2.77519  2.98518  4.58477  3.61503
          0.01758  4.44576  5.16811  0.61958  0.77255  0.48576  0.95510

42  2.64887  4.50233  5.05924  4.52219  3.70658  3.665984  5.07064  1.58337  4.41052  1.79216  3.53882  4.67727  4.87456  4.61926  4.55128  3.94042  3.45678  1.07310  5.55260  4.36863  147  v  -  -  -
2.68618  4.42225  2.77519  2.73123  3.46354  1.924104  3.72494  3.29354  2.67741  2.69355  4.24690  2.90347  2.73739  3.18146  2.89801  2.37887  2.77519  2.98518  4.58477  3.61503
          0.07937  4.44576  2.73997  0.61958  0.77255  0.48576  0.95510

43  2.60221  4.30470  4.53164  3.94761  3.27542  3.254408  4.40201  1.90675  3.80212  1.33579  2.36231  4.11167  4.40128  3.98645  3.93170  3.17550  3.13276  2.24894  4.95786  3.79919  148  l  -  -  -
2.68618  4.42225  2.77519  2.73123  3.46354  1.924104  3.72494  3.29354  2.67741  2.69355  4.24690  2.90347  2.73739  3.18146  2.89801  2.37887  2.77519  2.98518  4.58477  3.61503
          0.01869  4.38508  5.10742  0.61958  0.77255  0.57687  0.82475

44  2.84798  4.69392  3.73740  3.50554  4.27971  0.573224  4.53142  3.71831  3.50795  2.48593  4.44696  3.75089  4.19080  3.87843  3.77782  3.03308  3.29188  3.38321  5.68786  4.42915  149  g  -  -  -
2.68618  4.42225  2.77519  2.73123  3.46354  1.924104  3.72494  3.29354  2.67741  2.69355  4.24690  2.90347  2.73739  3.18146  2.89801  2.37887  2.77519  2.98518  4.58477  3.61503
          0.01869  4.38508  5.10742  0.61958  0.77255  0.57687  0.82475

45  1.95994  4.58628  3.33477  2.82820  4.18489  1.702584  3.96248  3.59448  2.82814  3.23553  4.05880  2.69865  3.90100  3.15580  3.25073  2.32823  2.19717  2.79127  5.49740  4.21229  150  a  -  -  -
2.68618  4.42225  2.77519  2.73123  3.46354  1.924104  3.72494  3.29354  2.67741  2.69355  4.24690  2.90347  2.73739  3.18146  2.89801  2.37887  2.77519  2.98518  4.58477  3.61503
          0.01869  4.38508  5.10742  0.61958  0.77255  0.57687  0.82475

46  2.20700  4.98012  3.10518  2.57644  4.34207  2.809384  3.74380  3.75680  2.41892  3.32027  4.12940  3.07734  3.93636  1.64031  2.48622  2.76888  2.61558  3.40011  5.51414  4.19285  151  q  -  -  -
2.68618  4.42225  2.77519  2.73123  3.46354  1.924104  3.72494  3.29354  2.67741  2.69355  4.24690  2.90347  2.73739  3.18146  2.89801  2.37887  2.77519  2.98518  4.58477  3.61503
          0.01869  4.38508  5.10742  0.61958  0.77255  0.43274  1.04620

47  2.55568  3.24253  4.51447  3.93216  3.36558  3.18816  4.35193  2.06675  3.78786  2.35328  2.82096  4.05904  4.34839  3.98274  3.90989  3.29669  2.34471  1.38438  4.93894  3.75095  152  v  -  -  -
2.68618  4.42225  2.77519  2.73123  3.46354  1.924104  3.72494  3.29354  2.67741  2.69355  4.24690  2.90347  2.73739  3.18146  2.89801  2.37887  2.77519  2.98518  4.58477  3.61503
          0.01758  4.44576  5.16811  0.61958  0.77255  0.48576  0.95510

48  3.52943  5.20981  4.31481  4.30055  5.36069  0.17488  5.34468  5.13918  4.55586  4.70143  5.70785  4.48216  4.61282  4.83556  4.69240  3.72289  4.05210  4.55636  6.26633  5.49441  153  G  -  -  -
2.68618  4.42225  2.77519  2.73123  3.46354  1.924104  3.72494  3.29354  2.67741  2.69355  4.24690  2.90347  2.73739  3.18146  2.89801  2.37887  2.77519  2.98518  4.58477  3.61503
          0.07937  4.44576  2.73997  0.61958  0.77255  0.48576  0.95510

49  1.92400  4.40204  3.63776  3.13984  4.05002  1.83984  4.14451  3.42019  3.11244  3.12319  3.97700  3.42823  3.93144  3.41394  3.47396  2.02070  2.12286  2.19743  5.43651  4.20259  154  a  -  -  -
2.68618  4.42225  2.77519  2.73123  3.46354  1.924104  3.72494  3.29354  2.67741  2.69355  4.24690  2.90347  2.73739  3.18146  2.89801  2.37887  2.77519  2.98518  4.58477  3.61503
          0.01869  4.38508  5.10742  0.61958  0.77255  0.57687  0.82475

50  3.71246  4.98507  5.11555  4.70096  0.93023  3.791624  4.33294  2.73449  4.52933  1.47525  3.34740  4.66055  4.98066  4.53117  4.55657  4.13887  3.94227  2.86181  4.47736  2.87820  155  f  -  -  -
2.68618  4.42225  2.77519  2.73123  3.46354  1.924104  3.72494  3.29354  2.67741  2.69355  4.24690  2.90347  2.73739  3.18146  2.89801  2.37887  2.77519  2.98518  4.58477  3.61503
          0.01869  4.38508  5.10742  0.61958  0.77255  0.43274  1.04620

51  2.22128  4.21318  4.44209  3.85688  2.56328  3.166352  4.29775  2.29061  3.71757  1.89806  3.29217  4.00596  4.31538  3.91285  3.85067  3.26221  2.63253  1.57715  4.89447  3.71008  156  v  -  -  -
2.68618  4.42225  2.77519  2.73123  3.46354  1.924104  3.72494  3.29354  2.67741  2.69355  4.24690  2.90347  2.73739  3.18146  2.89801  2.37887  2.77519  2.98518  4.58477  3.61503
          0.01758  4.44576  5.16811  0.61958  0.77255  0.48576  0.95510

52  3.52943  5.20981  4.31481  4.30055  5.36069  0.17488  5.34468  5.13918  4.55586  4.70143  5.70785  4.48216  4.61282  4.83556  4.69240  3.72289  4.05210  4.55636  6.26633  5.49441  157  G  -  -  -
2.68618  4.42225  2.77519  2.73123  3.46354  1.924104  3.72494  3.29354  2.67741  2.69355  4.24690  2.90347  2.73739  3.18146  2.89801  2.37887  2.77519  2.98518  4.58477  3.61503
          0.01758  4.44576  5.16811  0.61958  0.77255  0.48576  0.95510

53  2.44454  4.94371  3.04334  2.29942  4.23601  1.9462  3.72609  3.66291  2.50426  2.84568  4.04600  3.03199  3.90250  2.86557  2.64956  1.94669  2.93416  3.31490  5.46557  4.11952  158  s  -  -  -
2.68618  4.42225  2.77519  2.73123  3.46354  1.924104  3.72494  3.29354  2.67741  2.69355  4.24690  2.90347  2.73739  3.18146  2.89801  2.37887  2.77519  2.98518  4.58477  3.61503
          0.01758  4.44576  5.16811  0.61958  0.77255  0.48576  0.95510

54  2.68783  4.45704  4.76831  4.25177  3.80283  3.443384  4.91215  1.79275  4.14955  2.51919  3.65889  4.42645  4.70174  4.40756  4.34714  3.66984  2.72847  0.88643  5.55493  4.34650  159  v  -  -  -
2.68618  4.42225  2.77519  2.73123  3.46354  1.924104  3.72494  3.29354  2.67741  2.69355  4.24690  2.90347  2.73739  3.18146  2.89801  2.37887  2.77519  2.98518  4.58477  3.61503
          0.01758  4.44576  5.16811  0.61958  0.77255  0.48576  0.95510

55  3.15832  4.47178  4.97006  4.42321  3.65028  3.594456  4.95231  1.06158  4.30292  2.29863  2.81585  4.57887  4.79664  4.50988  4.44239  3.84108  2.76671  1.51678  5.45641  4.27605  160  i  -  -  -
2.68618  4.42225  2.77519  2.73123  3.46354  1.924104  3.72494  3.29354  2.67741  2.69355  4.24690  2.90347  2.73739  3.18146  2.89801  2.37887  2.77519  2.98518  4.58477  3.61503
          0.01758  4.44576  5.16811  0.61958  0.77255  0.48576  0.95510

56  3.52943  5.20981  4.31481  4.30055  5.36069  0.17488  5.34468  5.13918  4.55586  4.70143  5.70785  4.48216  4.61282  4.83556  4.69240  3.72289  4.05210  4.55636  6.26633  5.49441  161  G  -  -  -
2.68601  4.42229  2.77524  2.73127  3.46358  1.924136  3.72499  3.29358  2.67745  2.69359  4.24694  2.90325  2.73744  3.18150  2.89805  2.37891  2.77501  2.98522  4.58481  3.61507
          0.07937  2.65065  5.16811  0.87504  0.53930  0.48576  0.95510

57  0.86028  4.37401  3.92715  3.66222  4.71523  2.34652  4.66568  4.14213  3.71425  3.83662  4.64035  3.67559  3.95197  3.96260  3.98549  1.59943  2.96064  3.50482  6.05769  4.88469  165  a  -  -  -
2.68618  4.42225  2.77519  2.73123  3.46354  1.924104  3.72494  3.29354  2.67741  2.69355  4.24690  2.90347  2.73739  3.18146  2.89801  2.37887  2.77519  2.98518  4.58477  3.61503
          0.01758  4.44576  5.16811  0.61958  0.77255  0.48576  0.95510

58  2.79217  5.03931  3.16972  2.58728  4.33209  2.863688  3.72072  3.74219  1.99502  3.29970  4.10811  3.09325  3.96277  2.11974  2.24094  2.80434  2.65878  2.34554  5.48788  4.17297  166  k  -  -  -
2.68618  4.42225  2.77519  2.73123  3.46354  1.924104  3.72494  3.29354  2.67741  2.69355  4.24690  2.90347  2.73739  3.18146  2.89801  2.37887  2.77519  2.98518  4.58477  3.61503
          0.01758  4.44576  5.16811  0.61958  0.77255  0.48576  0.95510

59  2.04781  4.33447  4.53133  3.96205  3.43820  3.283152  4.48627  1.84095  3.83536  1.44360  3.36411  4.14384  4.45734  4.04791  3.99623  3.31086  3.16746  1.90205  5.09385  3.91353  167  l  -  -  -
2.68618  4.42225  2.77519  2.73123  3.46354  1.924104  3.72494  3.29354  2.67741  2.69355  4.24690  2.90347  2.73739  3.18146  2.89801  2.37887  2.77519  2.98518  4.58477  3.61503
          0.01758  4.44576  5.16811  0.61958  0.77255  0.48576  0.95510

60  2.19780  3.08937  3.95540  3.46628  3.89384  0.984384  4.30128  3.19108  3.38881  2.96106  2.93009  3.63967  4.02052  3.67016  3.67025  2.77916  2.93549  2.88297  5.34688  4.14396  168  g  -  -  -
2.68618  4.42225  2.77519  2.73123  3.46354  1.924104  3.72494  3.29354  2.67741  2.69355  4.24690  2.90347  2.73739  3.18146  2.89801  2.37887  2.77519  2.98518  4.58477  3.61503
          0.07937  4.44576  2.73997  0.61958  0.77255  0.48576  0.95510

61  2.21793  4.51758  3.34569  2.78293  3.67974  2.168752  3.81901  2.95559  2.71146  2.74171  3.18367  3.23829  3.96680  2.77007  3.15466  2.53294  2.59080  2.52842  5.08113  3.18776  169  a  -  -  -
2.68618  4.42225  2.77519  2.73123  3.46354  1.924104  3.72494  3.29354  2.67741  2.69355  4.24690  2.90347  2.73739  3.18146  2.89801  2.37887  2.77519  2.98518  4.58477  3.61503
          0.01869  4.38508  5.10742  0.61958  0.77255  0.57687  0.82475

62  2.27138  5.45324  1.96999  1.32439  4.78290  2.737968  3.83660  4.27709  2.73090  3.77404  4.56891  2.89427  3.96157  2.97404  3.27726  2.61954  3.20040  3.84718  5.93135  4.47382  170  e  -  -  -
2.68618  4.42225  2.77519  2.73123  3.46354  1.924104  3.72494  3.29354  2.67741  2.69355  4.24690  2.90347  2.73739  3.18146  2.89801  2.37887  2.77519  2.98518  4.58477  3.61503
          0.01869  4.38508  5.10742  0.61958  0.77255  0.57687  0.82475

63  2.45443  4.26730  3.86849  3.30305  3.43373  2.967632  4.06286  2.53686  3.22504  2.04336  3.41193  3.61153  4.11280  3.49673  3.50345  2.34519  2.25408  1.88982  4.93321  3.72346  171  v  -  -  -
2.68618  4.42225  2.77519  2.73123  3.46354  1.924104  3.72494  3.29354  2.67741  2.69355  4.24690  2.90347  2.73739  3.18146  2.89801  2.37887  2.77519  2.98518  4.58477  3.61503
          0.01869  4.38508  5.10742  0.61958  0.77255  0.57687  0.82475

64  1.84461  4.24068  4.13339  3.55712  3.35880  3.090584  4.18505  2.18667  3.45643  2.11519  2.72780  3.80980  4.22924  3.69660  3.67573  3.15117  2.39054  2.04548  4.91373  3.71887  172  a  -  -  -
2.68618  4.42225  2.77519  2.73123  3.46354  1.924104  3.72494  3.29354  2.67741  2.69355  4.24690  2.90347  2.73739  3.18146  2.89801  2.37887  2.77519  2.98518  4.58477  3.61503
          0.01869  4.38508  5.10742  0.61958  0.77255  0.57687  0.82475

65  2.72855  5.22141  2.10911  2.16186  4.54867  2.752416  2.88680  4.03004  2.44206  3.51873  4.27202  2.90895  3.18904  2.18453  2.94045  2.68659  2.63314  3.60355  5.66042  4.24680  173  d  -  -  -
2.68618  4.42225  2.77519  2.73123  3.46354  1.924104  3.72494  3.29354  2.67741  2.69355  4.24690  2.90347  2.73739  3.18146  2.89801  2.37887  2.77519  2.98518  4.58477  3.61503
          0.15506  4.38508  2.03124  0.61958  0.77255  0.57687  0.82475

66  2.71354  5.12783  2.80324  2.14804  4.43766  2.73932  3.29569  3.89732  2.40421  3.41412  4.19031  2.30719  3.85185  2.13394  2.87115  2.68488  2.33044  3.50287  5.57590  4.18948  174  q  -  -  -
2.68618  4.42225  2.77519  2.73123  3.46354  1.924104  3.72494  3.29354  2.67741  2.69355  4.24690  2.90347  2.73739  3.18146  2.89801  2.37887  2.77519  2.98518  4.58477  3.61503
          0.02139  4.25141  4.97375  0.61958  0.77255  0.73741  0.65076

67  3.20215  4.51289  5.01912  4.43842  2.59396  3.583904  4.81620  1.80673  4.29730  1.22142  2.72703  4.58013  4.71087  4.37046  4.35249  3.81795  3.42682  1.74044  5.14960  4.05830  175  l  -  -  -
2.68618  4.42225  2.77519  2.73123  3.46354  1.924104  3.72494  3.29354  2.67741  2.69355  4.24690  2.90347  2.73739  3.18146  2.89801  2.37887  2.77519  2.98518  4.58477  3.61503
          0.02139  4.25141  4.97375  0.61958  0.77255  0.73741  0.65076

68  2.62749  4.84672  2.44314  2.45719  3.84209  2.76236  3.65637  3.23471  2.25289  2.75533  3.91618  2.97907  3.84386  2.53188  2.91553  2.31800  2.85890  2.88753  5.34558  4.00608  176  k  -  -  -
2.68618  4.42225  2.77519  2.73123  3.46354  1.924104  3.72494  3.29354  2.67741  2.69355  4.24690  2.90347  2.73739  3.18146  2.89801  2.37887  2.77519  2.98518  4.58477  3.61503
          0.02139  4.25141  4.97375  0.61958  0.77255  0.73741  0.65076

69  2.80973  5.19953  2.99501  2.01144  4.55652  2.812864  3.10915  3.99583  2.23010  3.47033  4.25583  2.99399  3.91599  1.98332  2.03452  2.77790  3.02514  3.60392  5.58064  4.24694  177  q  -  -  -
2.68604  4.42232  2.77504  2.73130  3.46316  1.92416  3.72443  3.29361  2.67727  2.69362  4.24697  2.90353  2.73746  3.18153  2.89808  2.37894  2.77526  2.98525  4.58484  3.61510
          0.26877  1.84986  2.54562  0.82010  0.58051  0.73741  0.65076

70  2.58180  4.28378  3.53725  2.99033  2.88527  2.358032  3.87764  2.80719  2.93942  1.88968  3.42834  3.36306  3.15365  3.23731  3.27248  2.48641  2.83414  2.58154  4.89762  3.65881  182  l  -  -  -
2.68618  4.42225  2.77519  2.73123  3.46354  1.924104  3.72494  3.29354  2.67741  2.69355  4.24690  2.90347  2.73739  3.18146  2.89801  2.37887  2.77519  2.98518  4.58477  3.61503
          0.10514  4.17722  2.47142  0.61958  0.77255  0.80895  0.58938

71  2.64035  4.90395  2.80837  2.17667  4.18446  2.291144  3.64206  3.60784  2.42607  3.19025  3.99771  2.91490  3.81872  2.07126  2.88498  2.65085  2.78124  2.78576  5.40959  4.05807  183  q  -  -  -
2.68618  4.42225  2.77519  2.73123  3.46354  1.924104  3.72494  3.29354  2.67741  2.69355  4.24690  2.90347  2.73739  3.18146  2.89801  2.37887  2.77519  2.98518  4.58477  3.61503
          0.02500  4.09707  4.81942  0.61958  0.77255  0.87571  0.53883

72  2.57387  4.43280  3.27241  2.71635  3.00948  2.793952  3.74722  2.98343  2.68583  2.67143  3.54532  2.74932  3.16305  3.01054  3.08283  2.32400  2.50260  2.73398  4.03734  3.72953  184  s  -  -  -
2.68618  4.42225  2.77519  2.73123  3.46354  1.924104  3.72494  3.29354  2.67741  2.69355  4.24690  2.90347  2.73739  3.18146  2.89801  2.37887  2.77519  2.98518  4.58477  3.61503
          0.02500  4.09707  4.81942  0.61958  0.77255  0.87571  0.53883

73  2.60480  4.60089  2.41953  2.57640  3.80363  2.720856  3.71593  3.13706  2.58621  2.26386  3.70170  3.06512  3.86679  2.82254  3.01624  2.70457  2.58932  2.53571  5.16455  3.87190  185  l  -  -  -
2.68618  4.42225  2.77519  2.73123  3.46354  1.924104  3.72494  3.29354  2.67741  2.69355  4.24690  2.90347  2.73739  3.18146  2.89801  2.37887  2.77519  2.98518  4.58477  3.61503
          0.02500  4.09707  4.81942  0.61958  0.77255  0.60563  0.78906

74  2.04323  3.57731  2.95556  2.28985  4.01172  2.768424  3.68224  3.41977  2.28381  3.03903  3.86173  3.01478  3.20335  2.84438  2.95159  2.60549  2.85944  2.88516  5.29834  3.97520  186  a  -  -  -
2.68618  4.42225  2.77519  2.73123  3.46354  1.924104  3.72494  3.29354  2.67741  2.69355  4.24690  2.90347  2.73739  3.18146  2.89801  2.37887  2.77519  2.98518  4.58477  3.61503
          0.02139  4.25141  4.97375  0.61958  0.77255  0.73741  0.65076

75  2.16796  4.36981  3.16817  2.88679  3.35390  2.869784  3.83979  2.58306  2.84573  2.23556  3.31799  3.05579  3.96866  3.15352  3.21181  2.82922  2.30967  2.58467  4.95402  3.71366  187  a  -  -  -
2.68618  4.42225  2.77519  2.73123  3.46354  1.924104  3.72494  3.29354  2.67741  2.69355  4.24690  2.90347  2.73739  3.18146  2.89801  2.37887  2.77519  2.98518  4.58477  3.61503
          0.02139  4.25141  4.97375  0.61958  0.77255  0.73741  0.65076

76  2.63027  4.90161  2.66246  2.11041  4.16090  2.753648  3.64456  3.29502  2.42721  2.79930  3.96716  2.95556  3.83519  2.45875  2.89866  2.41007  2.70956  2.85010  5.39036  4.03835  188  e  -  -  -
2.68618  4.42225  2.77519  2.73123  3.46354  1.924104  3.72494  3.29354  2.67741  2.69355  4.24690  2.90347  2.73739  3.18146  2.89801  2.37887  2.77519  2.98518  4.58477  3.61503
          0.02139  4.25141  4.97375  0.61958  0.77255  0.73741  0.65076

77  2.21046  5.10475  2.75996  2.24405  4.42221  2.728296  3.62411  3.88995  2.18559  3.39844  4.15835  2.58344  3.17547  2.41047  2.86774  2.38788  2.89391  3.48161  5.55493  4.15823  189  k  -  -  -
2.68618  4.42225  2.77519  2.73123  3.46354  1.924104  3.72494  3.29354  2.67741  2.69355  4.24690  2.90347  2.73739  3.18146  2.89801  2.37887  2.77519  2.98518  4.58477  3.61503
          0.02139  4.25141  4.97375  0.61958  0.77255  0.46733  0.98530

78  2.46892  4.47309  3.48590  2.99825  4.00228  1.24956  4.04945  2.74142  2.98102  3.06006  3.92275  3.35476  3.02383  3.30098  3.35966  2.72241  2.24703  3.01758  5.38384  4.13377  190  g  -  -  -
2.68618  4.42225  2.77519  2.73123  3.46354  1.924104  3.72494  3.29354  2.67741  2.69355  4.24690  2.90347  2.73739  3.18146  2.89801  2.37887  2.77519  2.98518  4.58477  3.61503
          0.01869  4.38508  5.10742  0.61958  0.77255  0.57687  0.82475

79  2.44075  5.09196  2.64041  2.21308  4.40304  2.400704  3.65137  3.86439  2.06914  3.38427  4.14640  2.93542  2.85842  2.76202  2.89181  2.65257  2.62051  3.00969  5.54986  4.16101  191  k  -  -  -
2.68618  4.42225  2.77519  2.73123  3.46354  1.924104  3.72494  3.29354  2.67741  2.69355  4.24690  2.90347  2.73739  3.18146  2.89801  2.37887  2.77519  2.98518  4.58477  3.61503
          0.29184  4.38508  1.42440  0.61958  0.77255  0.57687  0.82475

80  2.67475  4.93135  2.73552  2.40009  4.29778  2.51748  2.91975  3.78353  2.52941  3.34839  4.16023  1.94482  3.83784  2.87500  2.98402  2.05797  2.95157  3.40199  5.52475  4.15155  192  n  -  -  -
2.68618  4.42225  2.77519  2.73123  3.46354  1.924104  3.72494  3.29354  2.67741  2.69355  4.24690  2.90347  2.73739  3.18146  2.89801  2.37887  2.77519  2.98518  4.58477  3.61503
          0.02449  4.11772  4.84007  0.61958  0.77255  0.47751  0.96844

81  2.71063  4.93401  2.53039  2.48454  4.26315  2.496896  3.76624  3.68348  2.44114  3.28146  4.09653  3.00640  1.77353  2.92003  3.05666  2.73366  2.97089  2.82049  5.51226  4.16311  193  p  -  -  -
2.68602  4.42256  2.77550  2.73116  3.46384  1.924344  3.72467  3.29375  2.67726  2.69256  4.24720  2.90352  2.73765  3.18177  2.89793  2.37902  2.77472  2.98521  4.58507  3.61534
          0.68383  0.71546  5.05937  1.05714  0.42686  0.40008  1.10946

82  2.48899  4.96075  3.02755  1.92122  4.22731  2.171304  3.70076  3.12978  2.48318  3.23527  4.03138  2.69899  3.88903  2.76189  2.95565  2.46209  2.78466  2.91035  5.45354  4.10004  206  e  -  -  -
2.68618  4.42225  2.77519  2.73123  3.46354  1.924104  3.72494  3.29354  2.67741  2.69355  4.24690  2.90347  2.73739  3.18146  2.89801  2.37887  2.77519  2.98518  4.58477  3.61503
          0.01758  4.44576  5.16811  0.61958  0.77255  0.48576  0.95510

83  2.62747  5.18473  2.64753  2.27080  4.52250  2.773736  3.65566  3.99965  1.90083  3.48412  4.22920  2.93817  3.86374  2.35817  2.48177  2.43845  2.51018  3.57184  5.61892  4.21742  207  k  -  -  -
2.68618  4.42225  2.77519  2.73123  3.46354  1.924104  3.72494  3.29354  2.67741  2.69355  4.24690  2.90347  2.73739  3.18146  2.89801  2.37887  2.77519  2.98518  4.58477  3.61503
          0.01758  4.44576  5.16811  0.61958  0.77255  0.48576  0.95510

84  2.63750  4.47254  5.03868  4.48969  3.63606  3.624608  4.98205  1.41653  4.36894  1.81780  3.23924  4.62802  4.82114  4.55879  4.48942  3.87998  3.41805  1.24676  5.45326  4.27769  208  v  -  -  -
2.68618  4.42225  2.77519  2.73123  3.46354  1.924104  3.72494  3.29354  2.67741  2.69355  4.24690  2.90347  2.73739  3.18146  2.89801  2.37887  2.77519  2.98518  4.58477  3.61503
          0.01758  4.44576  5.16811  0.61958  0.77255  0.48576  0.95510

85  2.11329  5.08325  2.67144  2.24056  4.38321  2.419184  3.67162  3.24060  2.24678  3.36929  4.13752  2.96156  3.86994  2.53459  2.91282  2.67344  2.41736  3.44962  5.54454  4.16217  209  a  -  -  -
2.68618  4.42225  2.77519  2.73123  3.46354  1.924104  3.72494  3.29354  2.67741  2.69355  4.24690  2.90347  2.73739  3.18146  2.89801  2.37887  2.77519  2.98518  4.58477  3.61503
          0.01758  4.44576  5.16811  0.61958  0.77255  0.48576  0.95510

86  2.27078  5.12926  2.94487  2.07373  4.44764  2.775744  3.66883  3.91214  2.23305  3.42419  4.18322  2.95155  3.24993  2.32287  2.89922  2.44255  2.41634  3.50676  5.58269  4.19156  210  e  -  -  -
2.68618  4.42225  2.77519  2.73123  3.46354  1.924104  3.72494  3.29354  2.67741  2.69355  4.24690  2.90347  2.73739  3.18146  2.89801  2.37887  2.77519  2.98518  4.58477  3.61503
          0.01758  4.44576  5.16811  0.61958  0.77255  0.48576  0.95510

87  2.23832  4.55170  3.35068  2.78778  3.27399  2.878288  3.20382  3.09099  2.75728  2.44658  3.64762  3.24678  3.98241  3.08406  2.52223  2.33172  2.61109  2.56032  5.11265  3.84859  211  a  -  -  -
2.68618  4.42225  2.77519  2.73123  3.46354  1.924104  3.72494  3.29354  2.67741  2.69355  4.24690  2.90347  2.73739  3.18146  2.89801  2.37887  2.77519  2.98518  4.58477  3.61503
          0.01758  4.44576  5.16811  0.61958  0.77255  0.48576  0.95510

88  2.68064  4.22771  3.96204  3.38892  3.35464  2.474608  4.06286  2.66466  3.29293  1.52417  2.87171  3.66329  4.12510  3.54815  3.53522  2.65462  2.58638  2.23983  4.85594  3.65501  212  l  -  -  -
2.68618  4.42225  2.77519  2.73123  3.46354  1.924104  3.72494  3.29354  2.67741  2.69355  4.24690  2.90347  2.73739  3.18146  2.89801  2.37887  2.77519  2.98518  4.58477  3.61503
          0.01758  4.44576  5.16811  0.61958  0.77255  0.48576  0.95510

89  2.68331  4.92863  3.05314  2.13838  4.18105  2.430512  3.70441  3.60554  2.49027  3.19365  3.31913  3.02283  3.89418  2.19319  2.95710  2.27815  2.63075  3.26887  3.80652  4.07707  213  e  -  -  -
2.68618  4.42225  2.77519  2.73123  3.46354  1.924104  3.72494  3.29354  2.67741  2.69355  4.24690  2.90347  2.73739  3.18146  2.89801  2.37887  2.77519  2.98518  4.58477  3.61503
          0.01758  4.44576  5.16811  0.61958  0.77255  0.48576  0.95510

90  2.73399  5.01283  3.08998  2.52322  4.29633  2.825344  3.70482  2.84983  1.79253  3.28268  4.08058  2.70406  3.92056  2.83598  2.33158  2.27487  2.96059  3.24019  5.48016  4.14147  214  k  -  -  -
2.68618  4.42225  2.77519  2.73123  3.46354  1.924104  3.72494  3.29354  2.67741  2.69355  4.24690  2.90347  2.73739  3.18146  2.89801  2.37887  2.77519  2.98518  4.58477  3.61503
          0.01758  4.44576  5.16811  0.61958  0.77255  0.48576  0.95510

91  3.12798  5.25645  3.62446  2.91823  4.61861  3.063504  3.29710  3.98939  1.73573  2.43393  4.32708  3.34286  4.17737  2.91744  1.27605  3.13671  3.29998  3.67759  5.56877  4.36341  215  r  -  -  -
2.68618  4.42225  2.77519  2.73123  3.46354  1.924104  3.72494  3.29354  2.67741  2.69355  4.24690  2.90347  2.73739  3.18146  2.89801  2.37887  2.77519  2.98518  4.58477  3.61503
          0.01758  4.44576  5.16811  0.61958  0.77255  0.48576  0.95510

92  1.91615  4.31904  3.78338  3.22398  3.50052  2.936728  4.04149  2.46326  3.15506  1.93307  3.47628  3.55521  4.09059  3.43902  3.45748  2.39470  2.38207  2.55276  4.98134  3.76581  216  a  -  -  -
2.68618  4.42225  2.77519  2.73123  3.46354  1.924104  3.72494  3.29354  2.67741  2.69355  4.24690  2.90347  2.73739  3.18146  2.89801  2.37887  2.77519  2.98518  4.58477  3.61503
          0.01758  4.44576  5.16811  0.61958  0.77255  0.48576  0.95510

93  2.04386  4.40941  3.78642  3.54574  4.80370  0.958648  4.63133  4.26118  3.66085  3.92438  4.71198  3.62176  3.94791  3.90335  3.96259  1.40709  2.97175  3.57958  6.11911  4.93260  217  g  -  -  -
2.68618  4.42225  2.77519  2.73123  3.46354  1.924104  3.72494  3.29354  2.67741  2.69355  4.24690  2.90347  2.73739  3.18146  2.89801  2.37887  2.77519  2.98518  4.58477  3.61503
          0.01758  4.44576  5.16811  0.61958  0.77255  0.48576  0.95510

94  2.41490  5.24127  2.11428  1.85419  4.58216  2.349208  3.73052  4.06137  2.52107  3.56065  4.32276  2.93672  3.90286  2.84504  3.02641  2.22574  2.65309  3.63786  5.71192  4.29734  218  e  -  -  -
2.68618  4.42225  2.77519  2.73123  3.46354  1.924104  3.72494  3.29354  2.67741  2.69355  4.24690  2.90347  2.73739  3.18146  2.89801  2.37887  2.77519  2.98518  4.58477  3.61503
          0.01758  4.44576  5.16811  0.61958  0.77255  0.48576  0.95510

95  2.87699  5.17553  3.22613  2.62558  4.51409  2.9024  3.71992  3.92972  1.33420  3.43488  4.23661  3.12332  4.00465  2.51219  2.33263  2.86800  2.68132  3.56939  5.56750  3.30170  219  k  -  -  -
2.68618  4.42225  2.77519  2.73123  3.46354  1.924104  3.72494  3.29354  2.67741  2.69355  4.24690  2.90347  2.73739  3.18146  2.89801  2.37887  2.77519  2.98518  4.58477  3.61503
          0.01758  4.44576  5.16811  0.61958  0.77255  0.48576  0.95510

96  2.60496  4.42902  4.90502  4.32713  3.38721  3.481008  4.71990  1.73754  4.18114  1.40475  2.42765  4.44798  4.64696  4.32387  4.26572  3.67995  3.32587  1.71545  5.17337  4.03183  220  l  -  -  -
2.68618  4.42225  2.77519  2.73123  3.46354  1.924104  3.72494  3.29354  2.67741  2.69355  4.24690  2.90347  2.73739  3.18146  2.89801  2.37887  2.77519  2.98518  4.58477  3.61503
          0.01758  4.44576  5.16811  0.61958  0.77255  0.48576  0.95510

97  3.52943  5.20981  4.31481  4.30055  5.36069  0.17488  5.34468  5.13918  4.55586  4.70143  5.70785  4.48216  4.61282  4.83556  4.69240  3.72289  4.05210  4.55636  6.26633  5.49441  221  G  -  -  -
2.68618  4.42225  2.77519  2.73123  3.46354  1.924104  3.72494  3.29354  2.67741  2.69355  4.24690  2.90347  2.73739  3.18146  2.89801  2.37887  2.77519  2.98518  4.58477  3.61503
          0.01758  4.44576  5.16811  0.61958  0.77255  0.48576  0.95510

98  2.44737  5.18806  2.70516  1.10129  4.69192  2.444128  3.91866  4.15543  2.78848  3.69992  4.50341  3.01468  3.98367  3.06780  3.28864  2.43829  3.16182  3.72379  5.87988  4.47849  222  e  -  -  -
2.68618  4.42225  2.77519  2.73123  3.46354  1.924104  3.72494  3.29354  2.67741  2.69355  4.24690  2.90347  2.73739  3.18146  2.89801  2.37887  2.77519  2.98518  4.58477  3.61503
          0.01758  4.44576  5.16811  0.61958  0.77255  0.48576  0.95510

99  2.53526  4.79389  3.12858  1.98816  4.12490  2.790632  3.81646  3.50104  2.64498  3.15194  3.98234  3.12060  3.93485  2.98769  3.09187  2.45746  1.79707  2.67945  5.42119  4.10541  223  t  -  -  -
2.68618  4.42225  2.77519  2.73123  3.46354  1.924104  3.72494  3.29354  2.67741  2.69355  4.24690  2.90347  2.73739  3.18146  2.89801  2.37887  2.77519  2.98518  4.58477  3.61503
          0.01758  4.44576  5.16811  0.61958  0.77255  0.48576  0.95510

100  1.90508  4.23851  4.21978  3.64032  3.35125  3.114104  4.21620  2.20290  3.52946  2.12136  2.50226  3.86606  4.25664  3.75773  3.72561  3.18403  2.34936  2.01789  4.90719  3.71588  224  a  -  -  -
2.68618  4.42225  2.77519  2.73123  3.46354  1.924104  3.72494  3.29354  2.67741  2.69355  4.24690  2.90347  2.73739  3.18146  2.89801  2.37887  2.77519  2.98518  4.58477  3.61503
          0.01758  4.44576  5.16811  0.61958  0.77255  0.48576  0.95510

101  3.52943  5.20981  4.31481  4.30055  5.36069  0.17488  5.34468  5.13918  4.55586  4.70143  5.70785  4.48216  4.61282  4.83556  4.69240  3.72289  4.05210  4.55636  6.26633  5.49441  225  G  -  -  -
2.68618  4.42225  2.77519  2.73123  3.46354  1.924104  3.72494  3.29354  2.67741  2.69355  4.24690  2.90347  2.73739  3.18146  2.89801  2.37887  2.77519  2.98518  4.58477  3.61503
          0.01758  4.44576  5.16811  0.61958  0.77255  0.48576  0.95510

102  2.26141  4.96126  3.02664  1.85762  4.22633  2.42092  3.70580  3.65569  2.48669  3.23470  4.03301  3.01365  3.89429  2.57039  2.95544  2.70612  2.40681  3.31003  3.79489  4.10258  226  e  -  -  -
2.68618  4.42225  2.77519  2.73123  3.46354  1.924104  3.72494  3.29354  2.67741  2.69355  4.24690  2.90347  2.73739  3.18146  2.89801  2.37887  2.77519  2.98518  4.58477  3.61503
          0.01758  4.44576  5.16811  0.61958  0.77255  0.48576  0.95510

103  2.41537  4.18118  4.06064  3.47755  3.29042  3.0172  4.07052  2.11206  3.36933  2.04103  2.84805  3.71898  4.14204  3.60481  3.01991  2.68687  2.58853  2.02681  4.79668  3.60073  227  v  -  -  -
2.68618  4.42225  2.77519  2.73123  3.46354  1.924104  3.72494  3.29354  2.67741  2.69355  4.24690  2.90347  2.73739  3.18146  2.89801  2.37887  2.77519  2.98518  4.58477  3.61503
          0.01758  4.44576  5.16811  0.61958  0.77255  0.48576  0.95510

104  2.16756  4.30505  3.75793  3.19694  2.94640  2.426096  4.01454  2.82514  3.13375  2.52720  3.47058  3.52775  4.06456  3.41302  3.43839  2.58138  2.11763  1.98849  4.95801  3.74204  228  v  -  -  -
2.68618  4.42225  2.77519  2.73123  3.46354  1.924104  3.72494  3.29354  2.67741  2.69355  4.24690  2.90347  2.73739  3.18146  2.89801  2.37887  2.77519  2.98518  4.58477  3.61503
          0.01758  4.44576  5.16811  0.61958  0.77255  0.48576  0.95510

105  3.52943  5.20981  4.31481  4.30055  5.36069  0.17488  5.34468  5.13918  4.55586  4.70143  5.70785  4.48216  4.61282  4.83556  4.69240  3.72289  4.05210  4.55636  6.26633  5.49441  229  G  -  -  -
2.68618  4.42225  2.77519  2.73123  3.46354  1.924104  3.72494  3.29354  2.67741  2.69355  4.24690  2.90347  2.73739  3.18146  2.89801  2.37887  2.77519  2.98518  4.58477  3.61503
          0.01758  4.44576  5.16811  0.61958  0.77255  0.48576  0.95510

106  1.84863  4.68623  3.25414  2.72246  4.00873  1.875136  3.84881  3.39224  2.41217  2.66941  3.89377  3.18850  3.94950  3.04525  3.11347  2.76168  2.33233  3.08681  5.33894  4.04905  230  a  -  -  -
2.68618  4.42225  2.77519  2.73123  3.46354  1.924104  3.72494  3.29354  2.67741  2.69355  4.24690  2.90347  2.73739  3.18146  2.89801  2.37887  2.77519  2.98518  4.58477  3.61503
          0.01758  4.44576  5.16811  0.61958  0.77255  0.48576  0.95510

107  2.20967  4.18902  4.13258  3.54884  3.29779  3.049088  4.11676  2.24797  3.43414  2.01875  2.83771  3.77562  4.17875  3.66439  3.03632  3.09681  2.59870  1.83549  4.81911  3.62571  231  v  -  -  -
2.68618  4.42225  2.77519  2.73123  3.46354  1.924104  3.72494  3.29354  2.67741  2.69355  4.24690  2.90347  2.73739  3.18146  2.89801  2.37887  2.77519  2.98518  4.58477  3.61503
          0.01758  4.44576  5.16811  0.61958  0.77255  0.48576  0.95510

108  2.28358  4.46966  4.97057  4.42998  3.70669  3.604824  4.98538  1.45945  4.31917  2.01817  3.55578  4.59050  4.81459  4.53781  4.47030  3.85708  3.40832  1.18375  5.50619  4.31332  232  v  -  -  -
2.68618  4.42225  2.77519  2.73123  3.46354  1.924104  3.72494  3.29354  2.67741  2.69355  4.24690  2.90347  2.73739  3.18146  2.89801  2.37887  2.77519  2.98518  4.58477  3.61503
          0.01758  4.44576  5.16811  0.61958  0.77255  0.48576  0.95510

109  2.10355  4.41801  3.84579  3.69481  4.78219  0.559672  4.75088  4.14482  3.81698  3.90454  4.74917  3.70869  3.99344  4.06825  4.05937  2.68341  3.02530  3.32686  6.14019  4.96331  233  g  -  -  -
2.68618  4.42225  2.77519  2.73123  3.46354  1.924104  3.72494  3.29354  2.67741  2.69355  4.24690  2.90347  2.73739  3.18146  2.89801  2.37887  2.77519  2.98518  4.58477  3.61503
          0.01758  4.44576  5.16811  0.61958  0.77255  0.48576  0.95510

110  2.46135  5.00696  3.04054  2.48362  4.28818  1.972024  3.69436  3.72178  2.24913  2.88543  4.07317  3.01102  3.89598  2.34271  2.60847  2.70829  2.40973  3.36434  5.48317  4.12938  234  k  -  -  -
2.68618  4.42225  2.77519  2.73123  3.46354  1.924104  3.72494  3.29354  2.67741  2.69355  4.24690  2.90347  2.73739  3.18146  2.89801  2.37887  2.77519  2.98518  4.58477  3.61503
          0.01758  4.44576  5.16811  0.61958  0.77255  0.48576  0.95510

111  2.02521  4.36897  4.56720  4.00437  3.55793  3.353792  4.58811  2.00049  3.89390  2.12140  3.24742  4.20989  4.53761  4.12501  4.07945  3.50961  3.06273  1.20360  5.22292  4.02893  235  v  -  -  -
2.68618  4.42225  2.77519  2.73123  3.46354  1.924104  3.72494  3.29354  2.67741  2.69355  4.24690  2.90347  2.73739  3.18146  2.89801  2.37887  2.77519  2.98518  4.58477  3.61503
          0.01758  4.44576  5.16811  0.61958  0.77255  0.48576  0.95510

112  2.37310  4.35014  4.70180  4.13816  3.55076  3.388216  4.64244  1.57795  4.01409  1.91575  3.46438  4.29662  4.57445  4.22652  4.16181  3.56044  2.70704  1.42456  5.22073  4.02894  236  v  -  -  -
2.68618  4.42225  2.77519  2.73123  3.46354  1.924104  3.72494  3.29354  2.67741  2.69355  4.24690  2.90347  2.73739  3.18146  2.89801  2.37887  2.77519  2.98518  4.58477  3.61503
          0.01758  4.44576  5.16811  0.61958  0.77255  0.48576  0.95510

113  1.84833  4.38674  4.53850  3.95336  2.72907  3.297656  4.43173  2.38071  3.82345  1.31703  2.68914  4.14253  4.44242  3.99391  3.96314  3.42605  3.19903  2.41382  4.97660  3.82490  237  l  -  -  -
2.68618  4.42225  2.77519  2.73123  3.46354  1.924104  3.72494  3.29354  2.67741  2.69355  4.24690  2.90347  2.73739  3.18146  2.89801  2.37887  2.77519  2.98518  4.58477  3.61503
          0.01758  4.44576  5.16811  0.61958  0.77255  0.48576  0.95510

114  3.52943  5.20981  4.31481  4.30055  5.36069  0.17488  5.34468  5.13918  4.55586  4.70143  5.70785  4.48216  4.61282  4.83556  4.69240  3.72289  4.05210  4.55636  6.26633  5.49441  238  G  -  -  -
2.68618  4.42225  2.77519  2.73123  3.46354  1.924104  3.72494  3.29354  2.67741  2.69355  4.24690  2.90347  2.73739  3.18146  2.89801  2.37887  2.77519  2.98518  4.58477  3.61503
          0.01758  4.44576  5.16811  0.61958  0.77255  0.48576  0.95510

115  3.69038  5.30981  4.36085  4.30438  5.19401  3.1738  5.29726  4.93654  4.42045  4.47609  5.57288  4.53911  0.22728  4.78704  4.56597  3.88312  4.17986  4.50607  6.17811  5.33292  239  P  -  -  -
2.68618  4.42225  2.77519  2.73123  3.46354  1.924104  3.72494  3.29354  2.67741  2.69355  4.24690  2.90347  2.73739  3.18146  2.89801  2.37887  2.77519  2.98518  4.58477  3.61503
          0.01758  4.44576  5.16811  0.61958  0.77255  0.48576  0.95510

116  1.98454  4.33396  3.80959  3.25702  3.55594  2.929552  4.08604  2.74697  3.19046  2.32250  3.52406  3.57941  3.07947  3.47563  3.49512  2.95815  2.30777  1.77137  5.04082  3.82467  240  v  -  -  -
2.68618  4.42225  2.77519  2.73123  3.46354  1.924104  3.72494  3.29354  2.67741  2.69355  4.24690  2.90347  2.73739  3.18146  2.89801  2.37887  2.77519  2.98518  4.58477  3.61503
          0.01758  4.44576  5.16811  0.61958  0.77255  0.48576  0.95510

117  3.52943  5.20981  4.31481  4.30055  5.36069  0.17488  5.34468  5.13918  4.55586  4.70143  5.70785  4.48216  4.61282  4.83556  4.69240  3.72289  4.05210  4.55636  6.26633  5.49441  241  G  -  -  -
2.68618  4.42225  2.77519  2.73123  3.46354  1.924104  3.72494  3.29354  2.67741  2.69355  4.24690  2.90347  2.73739  3.18146  2.89801  2.37887  2.77519  2.98518  4.58477  3.61503
          0.01758  4.44576  5.16811  0.61958  0.77255  0.48576  0.95510

118  1.56278  4.92507  2.98130  1.89918  4.35716  2.01128  3.85037  3.77203  2.67024  3.37441  3.97879  3.08178  3.94064  3.00457  3.13089  2.75888  3.00792  3.39961  5.60092  4.25688  242  a  -  -  -
2.68618  4.42225  2.77519  2.73123  3.46354  1.924104  3.72494  3.29354  2.67741  2.69355  4.24690  2.90347  2.73739  3.18146  2.89801  2.37887  2.77519  2.98518  4.58477  3.61503
          0.01758  4.44576  5.16811  0.61958  0.77255  0.48576  0.95510

119  2.61107  4.09924  4.41906  3.81962  3.19559  3.066424  4.15602  1.84662  3.65351  2.08371  2.78367  3.91515  4.19762  3.82867  3.74465  2.73292  2.65800  1.87737  3.49395  3.54286  243  i  -  -  -
2.68618  4.42225  2.77519  2.73123  3.46354  1.924104  3.72494  3.29354  2.67741  2.69355  4.24690  2.90347  2.73739  3.18146  2.89801  2.37887  2.77519  2.98518  4.58477  3.61503
          0.01758  4.44576  5.16811  0.61958  0.77255  0.48576  0.95510

120  2.23365  4.24996  4.55831  3.97850  2.84069  3.245296  4.41713  1.84838  3.84008  2.21109  3.35773  4.11786  4.40724  4.03668  3.96821  3.36821  2.65816  1.42784  4.99851  3.81083  244  v  -  -  -
2.68618  4.42225  2.77519  2.73123  3.46354  1.924104  3.72494  3.29354  2.67741  2.69355  4.24690  2.90347  2.73739  3.18146  2.89801  2.37887  2.77519  2.98518  4.58477  3.61503
          0.01758  4.44576  5.16811  0.61958  0.77255  0.48576  0.95510

121  3.52943  5.20981  4.31481  4.30055  5.36069  0.17488  5.34468  5.13918  4.55586  4.70143  5.70785  4.48216  4.61282  4.83556  4.69240  3.72289  4.05210  4.55636  6.26633  5.49441  245  G  -  -  -
2.68618  4.42225  2.77519  2.73123  3.46354  1.924104  3.72494  3.29354  2.67741  2.69355  4.24690  2.90347  2.73739  3.18146  2.89801  2.37887  2.77519  2.98518  4.58477  3.61503
          0.01758  4.44576  5.16811  0.61958  0.77255  0.48576  0.95510

122  2.25155  4.73468  3.18149  2.62235  3.27552  2.836032  3.76141  3.33332  2.60571  2.97801  3.21233  2.77350  3.93347  2.64903  2.72114  2.50464  2.21332  3.04710  5.26614  3.96411  246  t  -  -  -
2.68618  4.42225  2.77519  2.73123  3.46354  1.924104  3.72494  3.29354  2.67741  2.69355  4.24690  2.90347  2.73739  3.18146  2.89801  2.37887  2.77519  2.98518  4.58477  3.61503
          0.01758  4.44576  5.16811  0.61958  0.77255  0.48576  0.95510

123  2.42197  4.13659  4.31572  3.72265  2.79045  3.06628  4.14798  2.08825  3.57899  1.92422  2.79389  3.87112  4.19654  3.10961  3.71018  3.12843  2.59236  1.96792  4.75896  3.57533  247  l  -  -  -
2.68618  4.42225  2.77519  2.73123  3.46354  1.924104  3.72494  3.29354  2.67741  2.69355  4.24690  2.90347  2.73739  3.18146  2.89801  2.37887  2.77519  2.98518  4.58477  3.61503
          0.01758  4.44576  5.16811  0.61958  0.77255  0.48576  0.95510

124  2.60950  4.42599  4.93906  4.37558  3.52810  3.52724  4.81862  1.68363  4.24211  1.78730  2.77884  4.50352  4.71086  4.41719  4.34892  3.74576  3.34474  1.23600  5.29969  4.13262  248  v  -  -  -
2.68618  4.42225  2.77519  2.73123  3.46354  1.924104  3.72494  3.29354  2.67741  2.69355  4.24690  2.90347  2.73739  3.18146  2.89801  2.37887  2.77519  2.98518  4.58477  3.61503
          0.01758  4.44576  5.16811  0.61958  0.77255  0.48576  0.95510

125  3.52943  5.20981  4.31481  4.30055  5.36069  0.17488  5.34468  5.13918  4.55586  4.70143  5.70785  4.48216  4.61282  4.83556  4.69240  3.72289  4.05210  4.55636  6.26633  5.49441  249  G  -  -  -
2.68618  4.42225  2.77519  2.73123  3.46354  1.924104  3.72494  3.29354  2.67741  2.69355  4.24690  2.90347  2.73739  3.18146  2.89801  2.37887  2.77519  2.98518  4.58477  3.61503
          0.01758  4.44576  5.16811  0.61958  0.77255  0.48576  0.95510

126  2.80340  5.27104  2.43782  2.14486  4.60801  1.655224  3.75572  4.08760  2.56379  3.59026  4.35938  2.44054  3.91938  2.87522  3.07416  2.16792  2.65896  3.66683  5.74590  4.32749  250  g  -  -  -
2.68618  4.42225  2.77519  2.73123  3.46354  1.924104  3.72494  3.29354  2.67741  2.69355  4.24690  2.90347  2.73739  3.18146  2.89801  2.37887  2.77519  2.98518  4.58477  3.61503
          0.01758  4.44576  5.16811  0.61958  0.77255  0.48576  0.95510

127  1.83718  4.35634  4.36661  3.79787  3.40779  3.220592  4.39701  2.37146  3.68181  1.99628  1.96739  4.02475  4.38962  3.91025  3.87776  3.33360  3.14194  1.84151  5.06145  3.88483  251  a  -  -  -
2.68623  4.42230  2.77525  2.73129  3.46315  1.924024  3.72500  3.29359  2.67746  2.69360  4.24695  2.90352  2.73745  3.18152  2.89806  2.37892  2.77503  2.98496  4.58482  3.61508
          0.07937  2.65065  5.16811  1.05522  0.42789  0.48576  0.95510

128  3.40202  4.64427  5.35322  4.82604  3.54593  3.889664  5.35966  1.39658  4.72298  1.48099  3.42474  4.98299  5.07575  4.86420  4.82399  4.24483  3.64447  1.19308  5.67205  4.53805  256  v  -  -  -
2.68618  4.42225  2.77519  2.73123  3.46354  1.924104  3.72494  3.29354  2.67741  2.69355  4.24690  2.90347  2.73739  3.18146  2.89801  2.37887  2.77519  2.98518  4.58477  3.61503
          0.01758  4.44576  5.16811  0.61958  0.77255  0.48576  0.95510

129  1.88658  4.46255  3.70410  3.35501  4.28068  0.87676  4.38285  3.57608  3.35724  2.62172  4.23559  3.57032  4.01184  3.67149  3.67279  2.75153  3.00521  3.18552  5.68475  4.45395  257  g  -  -  -
2.68618  4.42225  2.77519  2.73123  3.46354  1.924104  3.72494  3.29354  2.67741  2.69355  4.24690  2.90347  2.73739  3.18146  2.89801  2.37887  2.77519  2.98518  4.58477  3.61503
          0.01758  4.44576  5.16811  0.61958  0.77255  0.48576  0.95510

130  1.73971  4.39104  3.83609  3.64488  4.82222  0.713952  4.71055  4.26935  3.78148  3.95296  4.74712  3.66720  3.95609  4.00773  4.05093  2.25318  2.98016  3.57959  6.15270  4.98376  258  g  -  -  -
2.68618  4.42225  2.77519  2.73123  3.46354  1.924104  3.72494  3.29354  2.67741  2.69355  4.24690  2.90347  2.73739  3.18146  2.89801  2.37887  2.77519  2.98518  4.58477  3.61503
          0.01758  4.44576  5.16811  0.61958  0.77255  0.48576  0.95510

131  3.21897  5.35689  3.39587  2.92408  4.72102  3.033024  3.90124  4.20231  2.28426  3.64789  4.53665  3.36456  4.22828  0.94010  2.16452  3.21539  3.42985  3.87209  5.69767  4.46028  259  q  -  -  -
2.68618  4.42225  2.77519  2.73123  3.46354  1.924104  3.72494  3.29354  2.67741  2.69355  4.24690  2.90347  2.73739  3.18146  2.89801  2.37887  2.77519  2.98518  4.58477  3.61503
          0.01758  4.44576  5.16811  0.61958  0.77255  0.48576  0.95510

132  2.65600  4.55031  4.97499  4.40490  3.15364  3.5884  4.85158  1.57301  4.27127  1.09053  3.22015  4.56494  4.75337  4.40116  4.36922  3.82106  3.45153  2.10107  5.26534  4.14313  260  l  -  -  -
2.68618  4.42225  2.77519  2.73123  3.46354  1.924104  3.72494  3.29354  2.67741  2.69355  4.24690  2.90347  2.73739  3.18146  2.89801  2.37887  2.77519  2.98518  4.58477  3.61503
          0.01758  4.44576  5.16811  0.61958  0.77255  0.48576  0.95510

133  2.07939  4.38703  3.93396  3.65169  4.51285  0.70224  4.62513  3.73932  3.67276  3.57785  4.45462  3.69724  3.99369  3.94170  3.93745  2.69068  2.98796  2.52665  5.91512  4.72695  261  g  -  -  -
2.68618  4.42225  2.77519  2.73123  3.46354  1.924104  3.72494  3.29354  2.67741  2.69355  4.24690  2.90347  2.73739  3.18146  2.89801  2.37887  2.77519  2.98518  4.58477  3.61503
          0.01758  4.44576  5.16811  0.61958  0.77255  0.48576  0.95510

134  2.88221  5.19922  2.71780  1.07246  4.66975  2.75712  3.91788  4.11051  2.76676  3.67438  4.48935  3.02587  3.99768  3.06942  3.24785  2.45189  2.57826  3.69876  5.86064  4.46652  262  e  -  -  -
2.68618  4.42225  2.77519  2.73123  3.46354  1.924104  3.72494  3.29354  2.67741  2.69355  4.24690  2.90347  2.73739  3.18146  2.89801  2.37887  2.77519  2.98518  4.58477  3.61503
          0.01758  4.44576  5.16811  0.61958  0.77255  0.48576  0.95510

135  3.31717  5.89270  0.70779  2.36043  5.21745  2.769944  4.13006  4.84455  3.26985  4.32802  5.22941  2.34907  4.14710  3.32157  3.92770  3.18043  3.64702  4.37613  6.42321  4.86670  263  d  -  -  -
2.68517  4.42242  2.77536  2.73140  3.46370  1.924232  3.72453  3.29371  2.67757  2.69351  4.24692  2.90363  2.73756  3.18163  2.89792  2.37903  2.77508  2.98484  4.58493  3.61520
          0.37369  1.39741  2.73997  1.01223  0.45163  0.48576  0.95510

136  2.03484  4.49207  3.45889  2.89285  3.57309  2.902544  3.86948  2.98952  2.78813  1.78752  3.57887  3.32251  4.01435  2.75181  2.74830  2.87526  2.92615  2.68344  5.06328  3.81838  268  l  -  -  -
2.68618  4.42225  2.77519  2.73123  3.46354  1.924104  3.72494  3.29354  2.67741  2.69355  4.24690  2.90347  2.73739  3.18146  2.89801  2.37887  2.77519  2.98518  4.58477  3.61503
          0.01869  4.38508  5.10742  0.61958  0.77255  0.57687  0.82475

137  2.41484  4.54788  3.31224  2.75107  3.71639  2.526096  3.80663  2.73903  2.72541  2.77545  3.64366  2.82299  3.95800  2.71035  3.13394  2.46622  2.59249  2.33386  5.10764  3.20066  269  v  -  -  -
2.68618  4.42225  2.77519  2.73123  3.46354  1.924104  3.72494  3.29354  2.67741  2.69355  4.24690  2.90347  2.73739  3.18146  2.89801  2.37887  2.77519  2.98518  4.58477  3.61503
          0.01869  4.38508  5.10742  0.61958  0.77255  0.57687  0.82475

138  2.45251  5.14107  2.94572  2.21433  4.46878  2.771824  3.10880  3.93367  2.36682  3.43303  4.19004  2.63954  3.86058  2.12443  2.34181  2.66883  2.62753  3.52318  5.57871  4.19154  270  q  -  -  -
2.68618  4.42225  2.77519  2.73123  3.46354  1.924104  3.72494  3.29354  2.67741  2.69355  4.24690  2.90347  2.73739  3.18146  2.89801  2.37887  2.77519  2.98518  4.58477  3.61503
          0.01869  4.38508  5.10742  0.61958  0.77255  0.57687  0.82475

139  2.78611  4.98060  2.91958  2.62708  4.57360  2.190496  3.93343  4.04158  2.69375  3.59895  4.41671  3.11309  3.97642  1.20610  3.09560  2.39249  3.10945  3.61285  5.76134  4.41981  271  q  -  -  -
2.68618  4.42225  2.77519  2.73123  3.46354  1.924104  3.72494  3.29354  2.67741  2.69355  4.24690  2.90347  2.73739  3.18146  2.89801  2.37887  2.77519  2.98518  4.58477  3.61503
          0.01869  4.38508  5.10742  0.61958  0.77255  0.57687  0.82475

140  2.65713  4.10400  4.23309  3.64145  2.76399  3.021344  3.25627  2.08654  3.50155  2.18991  3.21780  3.80069  3.28427  3.70232  3.64348  3.00504  2.88918  1.94009  2.95088  3.50252  272  v  -  -  -
2.68618  4.42225  2.77519  2.73123  3.46354  1.924104  3.72494  3.29354  2.67741  2.69355  4.24690  2.90347  2.73739  3.18146  2.89801  2.37887  2.77519  2.98518  4.58477  3.61503
          0.08455  4.38508  2.67929  0.61958  0.77255  0.57687  0.82475

141  2.18000  4.94358  3.00493  2.25307  4.21424  2.39516  3.66133  3.10122  2.06753  3.21536  4.01041  2.97744  3.85852  2.79371  2.59090  2.67059  2.67352  3.29409  5.42616  4.07508  273  k  -  -  -
2.68618  4.42225  2.77519  2.73123  3.46354  1.924104  3.72494  3.29354  2.67741  2.69355  4.24690  2.90347  2.73739  3.18146  2.89801  2.37887  2.77519  2.98518  4.58477  3.61503
          0.01995  4.32048  5.04282  0.61958  0.77255  0.66036  0.72705

142  2.70605  5.16655  2.90311  2.02195  4.50386  2.761616  3.07104  3.97161  2.34346  3.45863  4.21529  2.60695  3.85491  2.22751  2.32543  2.41688  2.93491  3.55454  5.59301  4.20503  274  e  -  -  -
2.68618  4.42225  2.77519  2.73123  3.46354  1.924104  3.72494  3.29354  2.67741  2.69355  4.24690  2.90347  2.73739  3.18146  2.89801  2.37887  2.77519  2.98518  4.58477  3.61503
          0.01995  4.32048  5.04282  0.61958  0.77255  0.66036  0.72705

143  2.69893  5.12786  2.53277  2.36786  4.44744  2.349264  3.66333  3.91221  2.17106  3.42705  4.19393  2.91380  2.31598  2.49121  2.91389  2.67144  2.60742  3.50858  5.58909  4.19442  275  k  -  -  -
2.68618  4.42225  2.77519  2.73123  3.46354  1.924104  3.72494  3.29354  2.67741  2.69355  4.24690  2.90347  2.73739  3.18146  2.89801  2.37887  2.77519  2.98518  4.58477  3.61503
          0.09045  4.32048  2.61468  0.61958  0.77255  0.51623  0.90824

144  2.66309  5.13336  2.59984  2.35569  4.46274  2.380768  3.62585  3.93700  2.18774  3.43161  4.18123  2.61097  3.82825  2.47948  2.57641  2.25931  2.54513  3.51619  5.57571  4.17472  276  k  -  -  -
2.68618  4.42225  2.77519  2.73123  3.46354  1.924104  3.72494  3.29354  2.67741  2.69355  4.24690  2.90347  2.73739  3.18146  2.89801  2.37887  2.77519  2.98518  4.58477  3.61503
          0.01995  4.32048  5.04282  0.61958  0.77255  0.66036  0.72705

145  2.42274  5.04486  2.92616  2.16425  4.00091  2.385272  3.64219  3.79826  2.21301  3.33196  4.10314  2.93025  2.62024  2.50049  2.88084  2.47337  2.89132  3.41223  5.50955  4.12957  277  e  -  -  -
2.68618  4.42225  2.77519  2.73123  3.46354  1.924104  3.72494  3.29354  2.67741  2.69355  4.24690  2.90347  2.73739  3.18146  2.89801  2.37887  2.77519  2.98518  4.58477  3.61503
          0.01995  4.32048  5.04282  0.61958  0.77255  0.51623  0.90824

146  2.53736  4.95359  3.01022  2.25451  4.22077  2.781672  3.67925  3.65211  2.23740  3.22609  4.02062  2.98913  3.24253  2.54718  2.92439  2.44476  2.39233  2.55345  5.44060  4.08630  278  k  -  -  -
2.68618  4.42225  2.77519  2.73123  3.46354  1.924104  3.72494  3.29354  2.67741  2.69355  4.24690  2.90347  2.73739  3.18146  2.89801  2.37887  2.77519  2.98518  4.58477  3.61503
          0.01869  4.38508  5.10742  0.61958  0.77255  0.57687  0.82475

147  1.84933  4.70548  3.04893  2.68438  4.23650  1.86328  3.86800  3.52761  2.69877  3.27148  4.08078  3.14155  2.64409  3.03379  3.14546  2.35663  2.57033  3.27328  5.51044  4.19657  279  a  -  -  -
2.68622  4.42229  2.77501  2.73125  3.46358  1.924136  3.72499  3.29358  2.67745  2.69359  4.24694  2.90351  2.73744  3.18120  2.89805  2.37891  2.77524  2.98495  4.58481  3.61507
          0.60961  1.60872  1.36143  0.20862  1.66976  0.57687  0.82475

148  2.58504  4.40763  3.00712  2.05469  3.99217  2.720664  3.67190  3.39507  2.48696  3.02280  3.85034  2.99102  3.82315  2.83954  2.93271  2.13723  2.37926  3.07834  5.28222  3.67032  281  e  -  -  -
2.68618  4.42225  2.77519  2.73123  3.46354  1.924104  3.72494  3.29354  2.67741  2.69355  4.24690  2.90347  2.73739  3.18146  2.89801  2.37887  2.77519  2.98518  4.58477  3.61503
          0.02500  4.09707  4.81942  0.61958  0.77255  0.87571  0.53883

149  2.01978  4.74578  2.89849  2.46565  4.03604  2.710784  3.02517  3.45057  2.48626  2.99613  3.88644  2.69334  3.81896  2.83387  2.93813  2.48910  2.50854  3.12434  5.31459  3.98824  282  a  -  -  -
2.68618  4.42225  2.77519  2.73123  3.46354  1.924104  3.72494  3.29354  2.67741  2.69355  4.24690  2.90347  2.73739  3.18146  2.89801  2.37887  2.77519  2.98518  4.58477  3.61503
          0.02500  4.09707  4.81942  0.61958  0.77255  0.83768  0.56689

150  1.92911  4.44456  3.17697  2.73410  3.77734  2.706328  3.80446  2.99550  2.71437  2.83187  3.69345  3.16253  3.01070  3.04014  3.11964  2.36846  2.61149  2.52207  5.15517  3.88845  283  a  -  -  -
2.68618  4.42225  2.77519  2.73123  3.46354  1.924104  3.72494  3.29354  2.67741  2.69355  4.24690  2.90347  2.73739  3.18146  2.89801  2.37887  2.77519  2.98518  4.58477  3.61503
          0.02449  4.11772  4.84007  0.61958  0.77255  0.74944  0.63986

151  2.63676  4.87347  2.92829  2.45114  4.21838  2.717552  3.68228  3.64613  2.23881  3.22441  4.02936  2.75794  2.39796  2.77028  2.89582  2.34609  2.39478  3.28792  5.43816  4.09636  284  k  -  -  -
2.68618  4.42225  2.77519  2.73123  3.46354  1.924104  3.72494  3.29354  2.67741  2.69355  4.24690  2.90347  2.73739  3.18146  2.89801  2.37887  2.77519  2.98518  4.58477  3.61503
          0.02305  4.17722  4.89956  0.61958  0.77255  0.80895  0.58938

152  2.37232  4.57506  3.16757  2.56061  3.90365  2.721688  3.76882  3.28396  2.62821  2.73896  3.79380  3.10472  2.40035  2.97022  3.05147  2.36716  2.33232  2.91330  5.24219  3.95415  285  t  -  -  -
2.68618  4.42225  2.77519  2.73123  3.46354  1.924104  3.72494  3.29354  2.67741  2.69355  4.24690  2.90347  2.73739  3.18146  2.89801  2.37887  2.77519  2.98518  4.58477  3.61503
          0.02305  4.17722  4.89956  0.61958  0.77255  0.80895  0.58938

153  2.62900  5.00803  2.78222  2.16160  4.30673  2.344216  3.61304  3.75886  2.37837  3.08802  4.07083  2.89208  3.29028  2.25292  2.86071  2.31460  2.81433  3.29522  5.47743  4.09763  286  e  -  -  -
2.68618  4.42225  2.77519  2.73123  3.46354  1.924104  3.72494  3.29354  2.67741  2.69355  4.24690  2.90347  2.73739  3.18146  2.89801  2.37887  2.77519  2.98518  4.58477  3.61503
          0.02305  4.17722  4.89956  0.61958  0.77255  0.80895  0.58938

154  2.62962  5.00337  2.89625  2.14804  4.30336  2.726416  3.60894  3.75303  2.25887  3.16439  4.06421  2.59289  3.29522  2.72703  2.67491  2.20943  2.56403  3.37118  5.46816  4.09355  287  e  -  -  -
2.68618  4.42225  2.77519  2.73123  3.46354  1.924104  3.72494  3.29354  2.67741  2.69355  4.24690  2.90347  2.73739  3.18146  2.89801  2.37887  2.77519  2.98518  4.58477  3.61503
          0.02305  4.17722  4.89956  0.61958  0.77255  0.80895  0.58938

155  2.60707  4.58032  2.76473  2.62136  3.77582  2.7902  3.72892  3.13295  2.56518  2.65643  3.07616  3.10263  3.88353  2.94194  3.03006  1.99279  2.84369  2.55346  5.14013  3.85440  288  s  -  -  -
2.68618  4.42225  2.77519  2.73123  3.46354  1.924104  3.72494  3.29354  2.67741  2.69355  4.24690  2.90347  2.73739  3.18146  2.89801  2.37887  2.77519  2.98518  4.58477  3.61503
          0.10514  4.17722  2.47142  0.61958  0.77255  0.80895  0.58938

156  2.32540  4.39181  3.33420  2.77998  3.55027  2.815088  3.52884  2.91037  2.72260  2.20456  3.49816  3.21407  3.16098  2.81713  3.09600  2.76504  2.82118  2.27775  4.96986  3.71968  289  l  -  -  -
2.68618  4.42225  2.77519  2.73123  3.46354  1.924104  3.72494  3.29354  2.67741  2.69355  4.24690  2.90347  2.73739  3.18146  2.89801  2.37887  2.77519  2.98518  4.58477  3.61503
          0.02500  4.09707  4.81942  0.61958  0.77255  0.87571  0.53883

157  2.36923  5.04868  2.65511  2.27807  4.34836  2.695152  3.25755  3.80481  2.39633  3.33693  4.11763  2.44689  3.80152  2.22824  2.87959  2.35188  2.89339  3.41845  5.51444  4.12702  290  q  -  -  -
2.68618  4.42225  2.77519  2.73123  3.46354  1.924104  3.72494  3.29354  2.67741  2.69355  4.24690  2.90347  2.73739  3.18146  2.89801  2.37887  2.77519  2.98518  4.58477  3.61503
          0.02500  4.09707  4.81942  0.61958  0.77255  0.87571  0.53883

158  2.58995  4.69491  2.69054  2.49109  3.50100  2.7518  3.65211  2.78085  2.23453  2.93975  3.77381  2.99765  3.82991  2.56933  2.92518  2.65240  2.52943  2.91014  5.21626  3.90293  291  k  -  -  -
2.68618  4.42225  2.77519  2.73123  3.46354  1.924104  3.72494  3.29354  2.67741  2.69355  4.24690  2.90347  2.73739  3.18146  2.89801  2.37887  2.77519  2.98518  4.58477  3.61503
          0.02500  4.09707  4.81942  0.61958  0.77255  0.39320  1.12360

159  2.66403  4.73504  2.79049  2.59335  3.54903  2.820392  3.73809  3.33357  2.19939  2.66542  3.81630  3.09418  3.91339  2.92050  2.65500  2.73778  2.37986  2.36221  5.26220  3.95691  292  k  -  -  -
2.68618  4.42225  2.77519  2.73123  3.46354  1.924104  3.72494  3.29354  2.67741  2.69355  4.24690  2.90347  2.73739  3.18146  2.89801  2.37887  2.77519  2.98518  4.58477  3.61503
          0.01869  4.38508  5.10742  0.61958  0.77255  0.43274  1.04620

160  2.71450  4.90106  3.11983  2.55804  4.14643  2.429432  3.72573  3.55780  2.44394  2.31048  3.97454  2.72879  3.92586  2.36349  2.22588  2.74988  2.94285  2.84486  5.39472  4.07126  293  r  -  -  -
2.68618  4.42225  2.77519  2.73123  3.46354  1.924104  3.72494  3.29354  2.67741  2.69355  4.24690  2.90347  2.73739  3.18146  2.89801  2.37887  2.77519  2.98518  4.58477  3.61503
          0.01758  4.44576  5.16811  0.61958  0.77255  0.48576  0.95510

161  2.70494  5.16428  2.63014  2.06139  4.48790  2.70968  3.66794  3.95909  2.41378  3.00470  4.21312  2.94200  3.86917  2.31634  2.61576  1.97500  2.93718  3.54387  5.60849  4.20997  294  s  -  -  -
2.68618  4.42225  2.77519  2.73123  3.46354  1.924104  3.72494  3.29354  2.67741  2.69355  4.24690  2.90347  2.73739  3.18146  2.89801  2.37887  2.77519  2.98518  4.58477  3.61503
          0.02617  4.44576  4.26113  0.61958  0.77255  0.48576  0.95510

162  4.42929  5.56869  5.04499  4.91290  3.47088  3.558472  4.76827  4.53143  4.65879  3.81153  5.12892  4.96815  5.01598  5.01146  4.67385  4.62846  4.76585  4.43253  0.24923  3.44989  295  W  -  -  -
2.68618  4.42225  2.77519  2.73123  3.46354  1.924104  3.72494  3.29354  2.67741  2.69355  4.24690  2.90347  2.73739  3.18146  2.89801  2.37887  2.77519  2.98518  4.58477  3.61503
          0.01773  4.43731  5.15966  0.61958  0.77255  0.49930  0.93384

163  2.42930  4.13464  4.44013  3.84226  2.58546  3.09368  4.17900  2.26609  3.67794  1.57002  3.19321  3.94367  4.22598  3.84896  3.77073  3.16907  2.59680  2.34725  2.92602  3.54513  296  l  -  -  -
2.68618  4.42225  2.77519  2.73123  3.46354  1.924104  3.72494  3.29354  2.67741  2.69355  4.24690  2.90347  2.73739  3.18146  2.89801  2.37887  2.77519  2.98518  4.58477  3.61503
          0.01773  4.43731  5.15966  0.61958  0.77255  0.49930  0.93384

164  2.67544  4.59811  2.88426  2.47961  3.77120  2.499776  3.80890  2.51891  2.71153  2.74988  3.02744  2.82500  3.96574  3.04103  3.12845  2.80401  2.61180  2.27714  5.15153  3.87735  297  v  -  -  -
2.68618  4.42225  2.77519  2.73123  3.46354  1.924104  3.72494  3.29354  2.67741  2.69355  4.24690  2.90347  2.73739  3.18146  2.89801  2.37887  2.77519  2.98518  4.58477  3.61503
          0.01773  4.43731  5.15966  0.61958  0.77255  0.49930  0.93384

165  2.73455  5.21768  2.62987  2.40942  4.56407  2.786144  3.66720  4.03804  1.77445  3.51424  4.26358  2.55046  3.88364  2.39410  2.33218  2.44830  2.96343  3.61026  5.64015  4.24606  298  k  -  -  -
2.68618  4.42225  2.77519  2.73123  3.46354  1.924104  3.72494  3.29354  2.67741  2.69355  4.24690  2.90347  2.73739  3.18146  2.89801  2.37887  2.77519  2.98518  4.58477  3.61503
          0.01773  4.43731  5.15966  0.61958  0.77255  0.49930  0.93384

166  2.62629  4.44549  3.55683  2.99681  3.58069  2.926256  3.19910  2.92613  2.91031  2.37270  2.99734  3.40208  4.05143  3.24732  3.25550  2.91544  1.54379  2.70676  5.03158  3.78896  299  t  -  -  -
2.68618  4.42225  2.77519  2.73123  3.46354  1.924104  3.72494  3.29354  2.67741  2.69355  4.24690  2.90347  2.73739  3.18146  2.89801  2.37887  2.77519  2.98518  4.58477  3.61503
          0.01773  4.43731  5.15966  0.61958  0.77255  0.49930  0.93384

167  1.96064  4.44778  3.67242  3.19670  4.29667  1.574544  4.24022  3.69379  3.17924  3.37112  4.19994  3.45868  3.93641  3.48183  3.54827  1.58051  2.37636  2.62192  5.64549  4.40845  300  s  -  -  -
2.68618  4.42225  2.77519  2.73123  3.46354  1.924104  3.72494  3.29354  2.67741  2.69355  4.24690  2.90347  2.73739  3.18146  2.89801  2.37887  2.77519  2.98518  4.58477  3.61503
          0.08007  4.43731  2.73152  0.61958  0.77255  0.49930  0.93384

168  2.67835  3.84115  2.64366  2.20808  4.45233  2.762792  3.64031  3.91962  2.19353  3.42099  4.17538  2.93058  3.84802  2.39256  2.10820  2.42267  2.90858  3.50758  5.56946  4.17781  301  r  -  -  -
2.68618  4.42225  2.77519  2.73123  3.46354  1.924104  3.72494  3.29354  2.67741  2.69355  4.24690  2.90347  2.73739  3.18146  2.89801  2.37887  2.77519  2.98518  4.58477  3.61503
          0.01886  4.37610  5.09845  0.61958  0.77255  0.56732  0.83711

169  2.39375  5.18242  2.06431  2.33842  4.51234  2.357032  3.68468  3.98673  2.24425  3.48960  4.24943  2.72334  3.86863  2.79636  2.95655  2.17120  2.78126  3.56956  5.64273  4.23612  302  d  -  -  -
2.68618  4.42225  2.77519  2.73123  3.46354  1.924104  3.72494  3.29354  2.67741  2.69355  4.24690  2.90347  2.73739  3.18146  2.89801  2.37887  2.77519  2.98518  4.58477  3.61503
          0.01869  4.38508  5.10742  0.61958  0.77255  0.57687  0.82475

170  2.38685  4.37425  3.63058  3.10712  1.63923  2.910512  3.96074  2.83309  3.04341  2.60070  3.52111  3.46716  4.06119  2.83543  3.36171  2.56101  2.94046  2.67180  4.92643  3.64086  303  f  -  -  -
2.68618  4.42225  2.77519  2.73123  3.46354  1.924104  3.72494  3.29354  2.67741  2.69355  4.24690  2.90347  2.73739  3.18146  2.89801  2.37887  2.77519  2.98518  4.58477  3.61503
          0.01869  4.38508  5.10742  0.61958  0.77255  0.57687  0.82475

171  1.64006  4.40777  4.37029  3.85877  3.68047  3.174856  4.58865  2.22184  3.75552  2.14887  3.57727  4.07277  4.43122  4.03130  3.99065  3.32793  3.19354  1.32061  5.35874  4.15267  304  v  -  -  -
2.68618  4.42225  2.77519  2.73123  3.46354  1.924104  3.72494  3.29354  2.67741  2.69355  4.24690  2.90347  2.73739  3.18146  2.89801  2.37887  2.77519  2.98518  4.58477  3.61503
          0.01869  4.38508  5.10742  0.61958  0.77255  0.43274  1.04620

172  3.03231  4.98166  3.34417  3.23588  4.91968  0.458096  4.52562  4.44738  2.81743  4.06022  4.95321  3.61130  4.24530  3.78421  3.67472  3.14726  3.47484  3.94866  6.07323  4.90424  305  G  -  -  -
2.68618  4.42225  2.77519  2.73123  3.46354  1.924104  3.72494  3.29354  2.67741  2.69355  4.24690  2.90347  2.73739  3.18146  2.89801  2.37887  2.77519  2.98518  4.58477  3.61503
          0.01758  4.44576  5.16811  0.61958  0.77255  0.48576  0.95510

173  3.60549  5.71692  2.91213  0.42385  5.16119  3.01128  4.45341  4.74762  3.42375  4.28213  5.27360  3.44361  4.40752  3.68362  3.86541  3.54963  3.92747  4.38484  6.21232  4.99465  306  E  -  -  -
2.68618  4.42225  2.77519  2.73123  3.46354  1.924104  3.72494  3.29354  2.67741  2.69355  4.24690  2.90347  2.73739  3.18146  2.89801  2.37887  2.77519  2.98518  4.58477  3.61503
          0.01758  4.44576  5.16811  0.61958  0.77255  0.48576  0.95510

174  1.99964  4.57192  3.54273  3.06953  4.22617  2.721216  4.11432  3.56763  2.92219  3.26327  4.12288  3.40772  3.99504  3.34044  2.66236  2.77009  1.20139  3.20182  5.55587  4.31099  307  t  -  -  -
2.68618  4.42225  2.77519  2.73123  3.46354  1.924104  3.72494  3.29354  2.67741  2.69355  4.24690  2.90347  2.73739  3.18146  2.89801  2.37887  2.77519  2.98518  4.58477  3.61503
          0.01758  4.44576  5.16811  0.61958  0.77255  0.48576  0.95510

175  1.57487  4.46306  3.54110  3.01093  3.84466  2.040528  4.01395  3.11746  2.97682  2.31453  3.78307  3.38245  3.98650  3.29066  3.34989  2.44081  2.53435  2.91426  5.25066  4.00647  308  a  -  -  -
2.68618  4.42225  2.77519  2.73123  3.46354  1.924104  3.72494  3.29354  2.67741  2.69355  4.24690  2.90347  2.73739  3.18146  2.89801  2.37887  2.77519  2.98518  4.58477  3.61503
          0.01758  4.44576  5.16811  0.61958  0.77255  0.48576  0.95510

176  2.66241  4.45557  3.52361  2.97358  3.66328  2.78164  3.93802  3.03759  2.91486  2.27287  3.63118  3.37394  4.01812  3.23999  3.27606  1.66116  2.14812  2.79067  4.45126  3.84202  309  s  -  -  -
2.68618  4.42225  2.77519  2.73123  3.46354  1.924104  3.72494  3.29354  2.67741  2.69355  4.24690  2.90347  2.73739  3.18146  2.89801  2.37887  2.77519  2.98518  4.58477  3.61503
          0.01758  4.44576  5.16811  0.61958  0.77255  0.48576  0.95510

177  1.59933  5.00815  2.94787  1.91938  4.32185  2.713816  3.77037  3.74341  2.56051  3.32763  4.13469  3.03167  3.93067  2.57731  3.02194  2.76008  2.99643  3.11097  5.54631  4.19312  310  a  -  -  -
2.68618  4.42225  2.77519  2.73123  3.46354  1.924104  3.72494  3.29354  2.67741  2.69355  4.24690  2.90347  2.73739  3.18146  2.89801  2.37887  2.77519  2.98518  4.58477  3.61503
          0.01758  4.44576  5.16811  0.61958  0.77255  0.48576  0.95510

178  2.39507  4.41673  3.55802  3.00393  3.61600  2.887576  3.94719  2.94714  2.95900  2.19724  3.57486  2.90428  4.02917  3.26639  3.31586  2.55447  1.96923  2.21390  5.05543  3.81911  311  t  -  -  -
2.68618  4.42225  2.77519  2.73123  3.46354  1.924104  3.72494  3.29354  2.67741  2.69355  4.24690  2.90347  2.73739  3.18146  2.89801  2.37887  2.77519  2.98518  4.58477  3.61503
          0.01758  4.44576  5.16811  0.61958  0.77255  0.48576  0.95510

179  3.27287  4.66641  5.28910  4.74727  3.53591  3.842056  5.25355  1.50448  4.62780  1.09071  3.31902  4.91394  5.01659  4.74780  4.71909  4.17440  3.64113  1.59074  5.56433  4.46078  312  l  -  -  -
2.68618  4.42225  2.77519  2.73123  3.46354  1.924104  3.72494  3.29354  2.67741  2.69355  4.24690  2.90347  2.73739  3.18146  2.89801  2.37887  2.77519  2.98518  4.58477  3.61503
          0.01758  4.44576  5.16811  0.61958  0.77255  0.48576  0.95510

180  2.34292  4.42069  3.71592  3.21985  2.92613  1.125416  4.10783  2.99331  3.17580  2.38024  3.67718  3.54078  4.08155  3.47416  3.49839  2.90596  2.98599  2.75853  5.12422  3.84802  313  g  -  -  -
2.68618  4.42225  2.77519  2.73123  3.46354  1.924104  3.72494  3.29354  2.67741  2.69355  4.24690  2.90347  2.73739  3.18146  2.89801  2.37887  2.77519  2.98518  4.58477  3.61503
          0.01758  4.44576  5.16811  0.61958  0.77255  0.48576  0.95510

181  2.06703  4.43737  3.68603  3.47656  4.76562  0.68744  4.59585  4.21528  3.60970  3.88186  4.68349  3.59515  3.96198  3.86967  3.74717  2.18106  2.99496  3.56916  6.08570  4.88988  314  g  -  -  -
2.68618  4.42225  2.77519  2.73123  3.46354  1.924104  3.72494  3.29354  2.67741  2.69355  4.24690  2.90347  2.73739  3.18146  2.89801  2.37887  2.77519  2.98518  4.58477  3.61503
          0.01758  4.44576  5.16811  0.61958  0.77255  0.48576  0.95510

182  2.00350  4.39748  3.56960  3.00355  3.54857  2.461936  3.92378  2.72534  2.95062  2.09435  3.51319  3.40360  4.03856  3.25683  2.86589  2.90088  2.35935  2.47854  4.99497  3.76164  315  a  -  -  -
2.68618  4.42225  2.77519  2.73123  3.46354  1.924104  3.72494  3.29354  2.67741  2.69355  4.24690  2.90347  2.73739  3.18146  2.89801  2.37887  2.77519  2.98518  4.58477  3.61503
          0.01758  4.44576  5.16811  0.61958  0.77255  0.48576  0.95510

183  3.25885  4.64797  5.28514  4.74617  3.56722  3.836016  5.25817  1.49076  4.62907  1.16306  3.35263  4.90824  5.01588  4.76007  4.72486  4.16767  3.62615  1.49057  5.58231  4.46806  316  l  -  -  -
2.68618  4.42225  2.77519  2.73123  3.46354  1.924104  3.72494  3.29354  2.67741  2.69355  4.24690  2.90347  2.73739  3.18146  2.89801  2.37887  2.77519  2.98518  4.58477  3.61503
          0.01758  4.44576  5.16811  0.61958  0.77255  0.48576  0.95510

184  2.25655  4.44516  3.68326  3.29545  4.21693  0.832424  4.31222  3.55715  3.27934  3.28625  2.97866  3.53083  3.99149  3.59679  3.60328  2.60668  2.97356  3.16839  5.61031  4.38297  317  g  -  -  -
2.68618  4.42225  2.77519  2.73123  3.46354  1.924104  3.72494  3.29354  2.67741  2.69355  4.24690  2.90347  2.73739  3.18146  2.89801  2.37887  2.77519  2.98518  4.58477  3.61503
          0.01758  4.44576  5.16811  0.61958  0.77255  0.48576  0.95510

185  2.77675  5.09893  3.10827  2.54798  4.46030  2.045048  3.71981  3.89308  1.99127  3.41563  4.20269  3.06072  3.94303  2.84083  1.88757  2.19776  3.00864  3.32750  5.57158  4.23525  318  r  -  -  -
2.68618  4.42225  2.77519  2.73123  3.46354  1.924104  3.72494  3.29354  2.67741  2.69355  4.24690  2.90347  2.73739  3.18146  2.89801  2.37887  2.77519  2.98518  4.58477  3.61503
          0.01758  4.44576  5.16811  0.61958  0.77255  0.48576  0.95510

186  2.41878  4.25009  4.04226  3.47431  3.39064  2.290192  4.13788  2.31906  3.37578  1.46074  3.37267  3.73421  4.17420  3.62802  3.61025  3.07090  2.64257  2.29895  4.91420  3.71591  319  l  -  -  -
2.68618  4.42225  2.77519  2.73123  3.46354  1.924104  3.72494  3.29354  2.67741  2.69355  4.24690  2.90347  2.73739  3.18146  2.89801  2.37887  2.77519  2.98518  4.58477  3.61503
          0.01758  4.44576  5.16811  0.61958  0.77255  0.48576  0.95510

187  2.55200  4.31996  4.80834  4.23139  2.82801  3.381712  4.60767  1.58335  4.08094  1.83349  3.35619  4.33034  4.55397  4.25137  4.17055  3.55247  3.05808  1.46555  5.11304  3.93719  320  v  -  -  -
2.68618  4.42225  2.77519  2.73123  3.46354  1.924104  3.72494  3.29354  2.67741  2.69355  4.24690  2.90347  2.73739  3.18146  2.89801  2.37887  2.77519  2.98518  4.58477  3.61503
          0.01758  4.44576  5.16811  0.61958  0.77255  0.48576  0.95510

188  2.02767  4.35188  3.77653  3.30742  3.74838  2.2568  4.17875  3.07349  3.25800  1.34006  3.72319  3.57053  4.04387  3.55012  3.55354  2.44232  2.94847  2.80465  5.20764  3.99347  321  l  -  -  -
2.68618  4.42225  2.77519  2.73123  3.46354  1.924104  3.72494  3.29354  2.67741  2.69355  4.24690  2.90347  2.73739  3.18146  2.89801  2.37887  2.77519  2.98518  4.58477  3.61503
          0.01758  4.44576  5.16811  0.61958  0.77255  0.48576  0.95510

189  2.97243  4.99236  3.12565  3.00752  4.78419  0.58028  4.35327  4.35288  3.20636  3.93062  4.81672  3.43059  4.17313  2.61703  3.55054  3.05931  3.38817  3.87587  5.99803  4.73066  322  g  -  -  -
2.68618  4.42225  2.77519  2.73123  3.46354  1.924104  3.72494  3.29354  2.67741  2.69355  4.24690  2.90347  2.73739  3.18146  2.89801  2.37887  2.77519  2.98518  4.58477  3.61503
          0.01758  4.44576  5.16811  0.61958  0.77255  0.48576  0.95510

190  2.06546  4.76193  3.18622  2.70958  4.41574  2.237056  3.92541  3.85720  2.74582  3.44431  4.23533  2.35018  2.07037  3.07887  3.19817  2.35136  2.53558  3.42020  5.65286  4.32655  323  a  -  -  -
2.68618  4.42225  2.77519  2.73123  3.46354  1.924104  3.72494  3.29354  2.67741  2.69355  4.24690  2.90347  2.73739  3.18146  2.89801  2.37887  2.77519  2.98518  4.58477  3.61503
          0.02617  4.44576  4.26113  0.61958  0.77255  0.48576  0.95510

191  2.46636  5.14212  2.93655  2.41887  4.46361  2.781248  3.50842  3.92532  2.25037  3.43697  4.20064  2.64962  2.84154  1.89076  2.88889  2.44757  2.95018  3.52258  5.59412  4.20759  324  q  -  -  -
2.68603  4.42230  2.77525  2.73107  3.46359  1.924144  3.72500  3.29359  2.67746  2.69360  4.24695  2.90352  2.73723  3.18152  2.89806  2.37892  2.77525  2.98524  4.58346  3.61508
          0.14656  2.03576  5.15966  0.66749  0.71948  0.49930  0.93384

192  1.49874  4.43535  3.85919  3.70213  4.83601  0.70848  4.76580  4.24806  3.84768  3.96965  4.79329  3.72238  4.00128  4.08142  4.09742  2.69844  3.04261  3.59388  6.15722  5.00932  327  g  -  -  -
2.68618  4.42225  2.77519  2.73123  3.46354  1.924104  3.72494  3.29354  2.67741  2.69355  4.24690  2.90347  2.73739  3.18146  2.89801  2.37887  2.77519  2.98518  4.58477  3.61503
          0.01773  4.43731  5.15966  0.61958  0.77255  0.49930  0.93384

193  2.27819  5.22669  3.04839  2.08478  4.58440  2.834056  3.68442  4.03446  1.97940  3.50960  4.27667  3.01797  3.93317  2.27392  2.04652  2.76681  3.01834  3.62685  5.62681  4.27108  328  k  -  -  -
2.68618  4.42225  2.77519  2.73123  3.46354  1.924104  3.72494  3.29354  2.67741  2.69355  4.24690  2.90347  2.73739  3.18146  2.89801  2.37887  2.77519  2.98518  4.58477  3.61503
          0.01773  4.43731  5.15966  0.61958  0.77255  0.49930  0.93384

194  2.31463  5.15571  2.63503  1.94775  4.47884  2.770256  3.66185  3.95001  2.22666  3.45094  3.41555  2.93823  3.86327  2.14909  2.89622  2.60447  2.92852  3.53477  5.60047  4.20184  329  e  -  -  -
2.68618  4.42225  2.77519  2.73123  3.46354  1.924104  3.72494  3.29354  2.67741  2.69355  4.24690  2.90347  2.73739  3.18146  2.89801  2.37887  2.77519  2.98518  4.58477  3.61503
          0.01773  4.43731  5.15966  0.61958  0.77255  0.49930  0.93384

195  2.31145  4.14115  4.39788  3.80468  3.24446  3.098616  4.19838  2.02417  3.65345  2.08682  2.79265  3.93183  4.23461  3.84104  3.76894  3.17374  2.59956  1.71704  4.78776  3.00495  330  v  -  -  -
2.68618  4.42225  2.77519  2.73123  3.46354  1.924104  3.72494  3.29354  2.67741  2.69355  4.24690  2.90347  2.73739  3.18146  2.89801  2.37887  2.77519  2.98518  4.58477  3.61503
          0.01773  4.43731  5.15966  0.61958  0.77255  0.49930  0.93384

196  3.51500  5.19723  4.29892  4.28337  5.34554  0.177952  5.32922  5.12090  4.53773  4.68488  5.69054  4.46624  4.60024  4.81808  4.67618  3.70810  4.03718  4.53966  6.25439  5.47866  331  G  -  -  -
2.68618  4.42225  2.77519  2.73123  3.46354  1.924104  3.72494  3.29354  2.67741  2.69355  4.24690  2.90347  2.73739  3.18146  2.89801  2.37887  2.77519  2.98518  4.58477  3.61503
          0.01773  4.43731  5.15966  0.61958  0.77255  0.49930  0.93384

197  2.24221  4.61665  3.28672  2.47149  3.79296  2.863736  3.80249  2.88184  2.57601  2.29765  3.13765  3.19806  3.96431  2.99656  2.41810  2.80235  2.61384  2.91172  5.16544  3.88957  332  a  -  -  -
2.68618  4.42225  2.77519  2.73123  3.46354  1.924104  3.72494  3.29354  2.67741  2.69355  4.24690  2.90347  2.73739  3.18146  2.89801  2.37887  2.77519  2.98518  4.58477  3.61503
          0.01773  4.43731  5.15966  0.61958  0.77255  0.49930  0.93384

198  3.05022  5.33751  3.40001  2.76291  4.77072  2.990048  3.74636  4.15404  1.44719  3.59158  4.40665  3.22758  4.10499  2.65100  1.61972  2.27752  3.23752  3.78721  5.65413  4.41031  333  k  -  -  -
2.68618  4.42225  2.77519  2.73123  3.46354  1.924104  3.72494  3.29354  2.67741  2.69355  4.24690  2.90347  2.73739  3.18146  2.89801  2.37887  2.77519  2.98518  4.58477  3.61503
          0.01773  4.43731  5.15966  0.61958  0.77255  0.47739  0.96864

199  2.27824  4.47966  4.82327  4.26146  3.47319  3.498176  4.77535  1.74641  4.13483  1.20643  3.33160  4.43687  4.68102  4.31725  4.27233  3.70572  3.36474  1.84578  5.28209  4.13168  334  l  -  -  -
2.68618  4.42225  2.77519  2.73123  3.46354  1.924104  3.72494  3.29354  2.67741  2.69355  4.24690  2.90347  2.73739  3.18146  2.89801  2.37887  2.77519  2.98518  4.58477  3.61503
          0.01758  4.44576  5.16811  0.61958  0.77255  0.48576  0.95510

200  3.52943  5.20981  4.31481  4.30055  5.36069  0.17488  5.34468  5.13918  4.55586  4.70143  5.70785  4.48216  4.61282  4.83556  4.69240  3.72289  4.05210  4.55636  6.26633  5.49441  335  G  -  -  -
2.68618  4.42225  2.77519  2.73123  3.46354  1.924104  3.72494  3.29354  2.67741  2.69355  4.24690  2.90347  2.73739  3.18146  2.89801  2.37887  2.77519  2.98518  4.58477  3.61503
          0.01758  4.44576  5.16811  0.61958  0.77255  0.48576  0.95510

201  2.45825  4.70640  3.20350  2.64423  3.90490  2.444104  3.77178  3.16357  2.40000  2.64726  2.91713  2.78352  3.93983  2.96722  3.06416  2.20312  2.50008  3.01615  5.24437  3.94824  336  s  -  -  -
2.68618  4.42225  2.77519  2.73123  3.46354  1.924104  3.72494  3.29354  2.67741  2.69355  4.24690  2.90347  2.73739  3.18146  2.89801  2.37887  2.77519  2.98518  4.58477  3.61503
          0.01758  4.44576  5.16811  0.61958  0.77255  0.48576  0.95510

202  2.69570  4.40915  3.58210  3.01299  3.24214  2.938936  3.91107  2.68002  2.94452  2.52129  3.50973  3.41242  4.05220  3.25891  1.97923  2.60313  2.92711  2.52218  4.95065  2.45878  337  r  -  -  -
2.68617  4.42226  2.77520  2.73122  3.46355  1.924104  3.72495  3.29355  2.67742  2.69356  4.24690  2.90347  2.73740  3.18144  2.89797  2.37886  2.77517  2.98519  4.58478  3.61504
          0.02258  4.09627  5.16811  0.82705  0.57507  0.48576  0.95510

203  2.24814  4.28828  4.74251  4.15541  2.78734  3.31748  4.50135  1.80518  3.99665  1.78317  2.97576  4.24867  4.47634  4.15360  4.07575  3.46514  3.16168  1.58175  5.00336  3.83726  346  v  -  -  -
2.68618  4.42225  2.77519  2.73123  3.46354  1.924104  3.72494  3.29354  2.67741  2.69355  4.24690  2.90347  2.73739  3.18146  2.89801  2.37887  2.77519  2.98518  4.58477  3.61503
          0.02617  4.44576  4.26113  0.61958  0.77255  0.48576  0.95510

204  2.52089  4.47559  3.65624  3.52205  4.86557  0.701176  4.68345  4.37603  3.73867  4.03591  4.83928  3.63253  3.99207  3.97487  4.02626  1.63689  3.05141  3.67567  6.17116  4.97887  347  g  -  -  -
2.68618  4.42225  2.77519  2.73123  3.46354  1.924104  3.72494  3.29354  2.67741  2.69355  4.24690  2.90347  2.73739  3.18146  2.89801  2.37887  2.77519  2.98518  4.58477  3.61503
          0.01773  4.43731  5.15966  0.61958  0.77255  0.49930  0.93384

205  3.47163  5.53394  3.98152  3.11535  5.10193  3.21356  3.83087  4.38180  1.59233  3.74513  4.63267  3.52852  4.35166  2.95925  0.85969  3.44891  3.59062  4.07292  5.71735  4.62517  348  r  -  -  -
2.68618  4.42225  2.77519  2.73123  3.46354  1.924104  3.72494  3.29354  2.67741  2.69355  4.24690  2.90347  2.73739  3.18146  2.89801  2.37887  2.77519  2.98518  4.58477  3.61503
          0.01773  4.43731  5.15966  0.61958  0.77255  0.49930  0.93384

206  2.73399  4.80077  3.23310  2.65635  3.27118  2.865256  3.40430  2.62109  1.83892  2.93960  3.88049  3.14643  3.96384  2.84846  2.17765  2.80414  2.95892  3.12161  5.30226  4.00763  349  k  -  -  -
2.68618  4.42225  2.77519  2.73123  3.46354  1.924104  3.72494  3.29354  2.67741  2.69355  4.24690  2.90347  2.73739  3.18146  2.89801  2.37887  2.77519  2.98518  4.58477  3.61503
          0.01773  4.43731  5.15966  0.61958  0.77255  0.49930  0.93384

207  3.23554  4.52922  5.06834  4.53162  3.68109  3.687136  5.09351  1.02565  4.41630  1.83662  3.50227  4.69811  4.89220  4.62272  4.55914  3.96877  2.86244  1.59853  5.56166  4.38848  350  i  -  -  -
2.68618  4.42225  2.77519  2.73123  3.46354  1.924104  3.72494  3.29354  2.67741  2.69355  4.24690  2.90347  2.73739  3.18146  2.89801  2.37887  2.77519  2.98518  4.58477  3.61503
          0.01773  4.43731  5.15966  0.61958  0.77255  0.49930  0.93384

208  3.11073  5.72439  1.23835  2.11696  5.02858  2.645312  3.94568  4.55729  2.92966  4.02686  4.84701  1.87521  4.04118  3.09730  3.52574  2.49331  3.39044  4.10689  6.17650  4.66421  351  d  -  -  -
2.68618  4.42225  2.77519  2.73123  3.46354  1.924104  3.72494  3.29354  2.67741  2.69355  4.24690  2.90347  2.73739  3.18146  2.89801  2.37887  2.77519  2.98518  4.58477  3.61503
          0.01773  4.43731  5.15966  0.61958  0.77255  0.49930  0.93384

209  3.17707  4.76126  4.20355  3.64354  2.97766  3.230368  3.98127  3.20757  3.08867  2.70933  3.47884  3.86372  4.43607  3.64669  2.89871  3.40173  3.41985  3.06476  1.01454  3.02179  352  w  -  -  -
2.68618  4.42225  2.77519  2.73123  3.46354  1.924104  3.72494  3.29354  2.67741  2.69355  4.24690  2.90347  2.73739  3.18146  2.89801  2.37887  2.77519  2.98518  4.58477  3.61503
          0.06412  4.43731  2.99023  0.61958  0.77255  0.49930  0.93384

210  2.43315  5.14347  2.71561  2.17566  4.46968  2.746112  3.63769  3.94177  2.38191  3.43923  4.19316  2.25726  3.83877  2.48294  2.35121  2.64765  2.61009  3.52475  5.58578  4.18637  353  e  -  -  -
2.68618  4.42225  2.77519  2.73123  3.46354  1.924104  3.72494  3.29354  2.67741  2.69355  4.24690  2.90347  2.73739  3.18146  2.89801  2.37887  2.77519  2.98518  4.58477  3.61503
          0.08978  4.32760  2.62181  0.61958  0.77255  0.56749  0.83689

211  2.27108  4.96258  2.59336  2.40766  4.11369  2.260952  3.66282  3.76255  2.44032  3.31117  4.08979  2.60717  3.39270  2.78700  2.92174  2.15060  2.70962  3.37501  5.50043  4.12861  354  s  -  -  -
2.68600  4.42233  2.77522  2.73120  3.46362  1.924136  3.72502  3.29362  2.67738  2.69363  4.24673  2.90332  2.73726  3.18137  2.89808  2.37894  2.77522  2.98526  4.58485  3.61511
          0.60585  1.44974  1.51524  0.59813  0.79815  0.65174  0.73635

212  2.24423  4.54564  3.12950  2.60712  3.88814  2.689072  3.72509  3.07121  2.25804  2.92341  3.77231  3.06567  3.81950  2.92328  2.95570  2.32022  2.21951  2.94510  5.21546  3.93047  366  t  -  -  -
2.68631  4.42238  2.77532  2.73114  3.46367  1.92408  3.72449  3.29367  2.67724  2.69368  4.24702  2.90359  2.73688  3.18159  2.89813  2.37899  2.77515  2.98503  4.58490  3.61516
          0.66773  0.82151  3.04997  0.66685  0.72015  0.62872  0.76201

213  2.52532  4.76669  3.03142  2.49766  4.06383  2.734144  3.66952  3.37317  2.43801  3.07982  3.90101  2.99895  2.41096  2.50262  2.68289  2.57916  2.45471  3.14612  5.32231  4.00415  369  p  -  -  -
2.68618  4.42225  2.77519  2.73123  3.46354  1.924104  3.72494  3.29354  2.67741  2.69355  4.24690  2.90347  2.73739  3.18146  2.89801  2.37887  2.77519  2.98518  4.58477  3.61503
          0.03599  4.13221  3.94758  0.61958  0.77255  0.77891  0.61416

214  2.60647  4.83460  2.61403  2.12064  4.08272  2.731224  3.63040  2.98483  2.42872  3.09799  3.90752  2.93823  3.81405  2.62537  2.89613  2.38291  2.74507  2.77440  5.33514  3.99216  370  e  -  -  -
2.68618  4.42225  2.77519  2.73123  3.46354  1.924104  3.72494  3.29354  2.67741  2.69355  4.24690  2.90347  2.73739  3.18146  2.89801  2.37887  2.77519  2.98518  4.58477  3.61503
          0.04856  4.12063  3.46834  0.61958  0.77255  0.78902  0.60566

215  2.52331  4.36330  3.38634  2.85787  3.67110  2.727016  3.86303  2.96900  2.80982  2.36497  3.61028  3.25079  2.49062  3.13930  3.17958  2.22164  2.70573  2.36046  5.09112  3.84226  371  s  -  -  -
2.68618  4.42225  2.77519  2.73123  3.46354  1.924104  3.72494  3.29354  2.67741  2.69355  4.24690  2.90347  2.73739  3.18146  2.89801  2.37887  2.77519  2.98518  4.58477  3.61503
          0.06872  4.09707  2.99990  0.61958  0.77255  0.80895  0.58938

216  2.28809  5.01513  2.22712  2.02179  4.29501  2.679552  3.63767  3.73480  2.45688  2.77140  4.09707  2.64647  3.80376  2.77473  2.94956  2.65017  2.91058  3.36877  5.50051  4.11667  372  e  -  -  -
2.68583  4.42231  2.77526  2.73129  3.46360  1.924152  3.72501  3.29313  2.67747  2.69356  4.24696  2.90353  2.73746  3.18152  2.89807  2.37878  2.77526  2.98524  4.58483  3.61509
          0.54060  0.89361  4.77680  0.15697  1.92915  0.63380  0.75624

217  2.60206  3.44830  3.13161  2.58765  3.90507  2.758128  3.71639  3.29837  2.29127  2.94265  3.78469  2.85623  3.28000  2.80935  2.95896  2.68777  1.99876  3.00262  5.22532  3.93075  374  t  -  -  -
2.68618  4.42225  2.77519  2.73123  3.46354  1.924104  3.72494  3.29354  2.67741  2.69355  4.24690  2.90347  2.73739  3.18146  2.89801  2.37887  2.77519  2.98518  4.58477  3.61503
          0.03628  4.17722  3.89784  0.61958  0.77255  0.73741  0.65076

218  2.26165  4.66289  3.08611  2.60409  4.11468  2.679752  3.77462  3.51057  2.26154  3.14054  3.96806  3.06831  2.32813  2.94870  2.98367  2.61560  2.18719  3.15912  5.39151  4.08913  375  t  -  -  -
2.68618  4.42225  2.77519  2.73123  3.46354  1.924104  3.72494  3.29354  2.67741  2.69355  4.24690  2.90347  2.73739  3.18146  2.89801  2.37887  2.77519  2.98518  4.58477  3.61503
          0.02336  4.16430  4.88664  0.61958  0.77255  0.74969  0.63963

219  1.90152  4.70234  3.02393  2.52797  4.12777  2.682208  3.75487  3.53585  2.56267  3.15434  3.97699  3.03590  3.83696  2.29685  2.99423  2.25942  2.48213  3.18300  5.40020  4.08581  376  a  -  -  -
2.68618  4.42225  2.77519  2.73123  3.46354  1.924104  3.72494  3.29354  2.67741  2.69355  4.24690  2.90347  2.73739  3.18146  2.89801  2.37887  2.77519  2.98518  4.58477  3.61503
          0.04353  4.16430  3.61004  0.61958  0.77255  0.74969  0.63963

220  2.20003  4.41919  3.39440  3.20154  4.37795  2.52504  4.33760  3.72303  3.27805  3.44897  4.38869  3.43354  0.99576  3.62444  3.56926  2.67067  2.97609  3.28002  5.72389  4.51104  377  p  -  -  -
2.68618  4.42225  2.77519  2.73123  3.46354  1.924104  3.72494  3.29354  2.67741  2.69355  4.24690  2.90347  2.73739  3.18146  2.89801  2.37887  2.77519  2.98518  4.58477  3.61503
          0.07253  4.14460  2.91671  0.61958  0.77255  0.76784  0.62365

221  2.36109  4.84002  2.93331  2.15580  4.09395  2.690024  3.63253  2.96450  2.35101  3.10496  3.91836  2.94321  3.82086  2.25088  2.85755  2.52728  2.85497  3.18018  5.33836  4.00196  378  e  -  -  -
2.68618  4.42225  2.77519  2.73123  3.46354  1.924104  3.72494  3.29354  2.67741  2.69355  4.24690  2.90347  2.73739  3.18146  2.89801  2.37887  2.77519  2.98518  4.58477  3.61503
          0.02500  4.09707  4.81942  0.61958  0.77255  0.80895  0.58938

222  2.60899  4.49449  3.20936  2.66744  3.75581  2.736256  3.86201  3.08527  2.74049  2.09309  3.73074  3.20796  3.91578  3.12141  3.09696  1.70974  2.90168  2.82841  5.17202  3.88769  379  s  -  -  -
2.68618  4.42225  2.77519  2.73123  3.46354  1.924104  3.72494  3.29354  2.67741  2.69355  4.24690  2.90347  2.73739  3.18146  2.89801  2.37887  2.77519  2.98518  4.58477  3.61503
          0.03216  4.09707  4.19805  0.61958  0.77255  0.80895  0.58938

223  2.62181  4.75239  2.93210  2.49756  4.08355  2.700152  3.70832  3.49936  2.41914  2.81076  3.94799  2.47678  2.18986  2.87962  2.91799  2.49454  2.88724  3.16841  5.35560  4.03530  380  p  -  -  -
2.68618  4.42225  2.77519  2.73123  3.46354  1.924104  3.72494  3.29354  2.67741  2.69355  4.24690  2.90347  2.73739  3.18146  2.89801  2.37887  2.77519  2.98518  4.58477  3.61503
          0.02518  4.09010  4.81244  0.61958  0.77255  0.81468  0.58479

224  2.33557  4.76935  2.99127  2.46619  4.05241  2.491232  3.65155  3.46398  2.36914  3.06975  3.88911  2.97355  2.57352  2.49124  2.77340  2.64276  2.84334  2.90731  5.31234  3.98895  381  a  -  -  -
2.68618  4.42225  2.77519  2.73123  3.46354  1.924104  3.72494  3.29354  2.67741  2.69355  4.24690  2.90347  2.73739  3.18146  2.89801  2.37887  2.77519  2.98518  4.58477  3.61503
          0.03239  4.09010  4.19107  0.61958  0.77255  0.81468  0.58479

225  2.33286  4.79889  2.92987  2.15208  4.08784  2.711352  3.64930  3.50181  2.44426  3.10398  3.91911  2.94805  2.69377  2.71206  2.86358  2.63384  2.51511  3.04121  5.34286  4.00813  382  e  -  -  -
2.68618  4.42225  2.77519  2.73123  3.46354  1.924104  3.72494  3.29354  2.67741  2.69355  4.24690  2.90347  2.73739  3.18146  2.89801  2.37887  2.77519  2.98518  4.58477  3.61503
          0.05472  4.08307  3.31343  0.61958  0.77255  0.55031  0.85984

226  2.35818  5.05185  2.43819  2.08753  4.35764  2.563528  3.59388  3.81975  2.15368  3.24071  4.11344  2.66012  3.70110  2.71171  2.86080  2.60383  2.68660  3.42307  5.51114  4.11651  383  e  -  -  -
2.68636  4.42148  2.77525  2.73055  3.46393  1.924352  3.72533  3.29393  2.67754  2.69311  4.24729  2.90360  2.73692  3.18115  2.89827  2.37883  2.77525  2.98550  4.58516  3.61464
          0.58326  0.81662        *  1.61809  0.22099  0.00000        *

//
```
